# Transcriptional response of the calcification and stress response toolkits in an octocoral under heat and pH stress

**DOI:** 10.1101/2020.07.15.202069

**Authors:** Sergio Vargas, Thorsten Zimmer, Nicola Conci, Martin Lehmann, Gert Wörheide

## Abstract

Up to one-third of all described marine species inhabit coral reefs, but the future of these hyperdiverse ecosystems is insecure due to local and global threats, such as overfishing, eutrophication, ocean warming, and acidification. Although these impacts are expected to have a net detrimental effect on reefs, it has been shown that some organisms like octocorals may remain unaffected, or benefit from, anthropogenically induced environmental change, and may replace stony corals in future reefs. Despite their potential importance in future shallow-water coastal environments, the molecular mechanisms leading to the resilience to anthropogenic-induced stress observed in octocorals remain unknown. Here, we use manipulative experiments, proteomics, and transcriptomics to show that the molecular toolkit used by Pinnigorgia flava, a common Indo-Pacific gorgonian octocoral, to deposit its calcium-carbonate skeleton is resilient to heat and seawater acidification stress. Sublethal heat stress triggered a stress response in P. flava but did not affect the expression of 27 transcripts encoding Skeletal Organic Matrix (SOM) proteins. Exposure to seawater acidification did not cause a stress response but triggered the downregulation of many transcripts, including an osteonidogen homolog present in the SOM. The observed transcriptional decoupling of the skeletogenic and stress-response toolkits provides insights into the mechanisms of resilience to anthropogenically-driven environmental change observed in octocorals.

## Introduction

As a consequence of anthropogenic-induced global climate change, extreme climatic events, like the heat waves affecting the Great Barrier Reef in 2016, 2017, and 2020, are likely to increase in frequency, impacting marine communities in unprecedented ways (Hughes et al., 2017, 2018, 2019). In coral reefs, these phenomena can lead to the displacement of stony corals as the dominant reef-building taxon by other benthic organisms and can cause community phase shifts with ecosystem-level effects (Done, 1992). Both theoretical (Fung, Seymour, & Johnson, 2011) and empirical (Schmitt, Holbrook, Davis, Brooks, & Adam, 2019) studies predict and support coral reef phase shifts leading to the dominance of diverse groups, including algae, sponges, and other cnidarians such as corallimorpharians and octocorals in these ecosystems (see Norström, Nyström, Lokrantz, & Folke, 2009 for a review).

Octocorals (e.g., soft-corals, gorgonians) are common and important community members in many marine ecosystems. They increase the spatial complexity of the habitats they inhabit (Quattrini et al., 2014) and provide refuge to numerous invertebrate species (Buhl-Mortensen & Mortensen, 2005; Cúrdia et al., 2015). In shallow water ecosystems, such as coral reefs, octocorals can outgrow stony corals and become dominant after extreme climatic events, like anomalously strong El Niño events (Ruzicka et al., 2013), or under extreme environmental conditions, like those prevailing in volcanic-seep acidified waters (Inoue, Kayanne, Yamamoto, & Kurihara, 2013). For instance, in Florida, octocorals increased in abundance in the 11 years following the 1997/98 El Niño event, becoming the most abundant taxon in some localities and dominating shallow fore-reefs (Ruzicka et al., 2013). Similarly, the soft coral genus *Rhytisma* became dominant after a significant coral bleaching event in the Aldabra Atoll (Indian ocean) in 1998 (Spencer et al., 2005), and the “blue coral,” *Heliopora coerulea*, a species of the reef-building zooxanthellate octocoral clade Helioporacea, increased its abundance to > 50% over a decade in the Bolinao Reef Complex in the northern Philippines (Atrigenio, Aliño, & Conaco, 2017). Like scleractinians, helioporacean octocorals produce a rigid skeleton made of aragonite (a polymorph of calcium carbonate) (Colgan, 1984), but their calcification machinery appears to be more resilient to environmental stressors than that of most scleractinians (Kayanne, Harii, Ide, & Akimoto, 2002; Shaish, Levy, Katzir, & Rinkevich, 2010). Thus, octocorals may be possible winners in future oceans due to global climate change, potentially replacing scleractinians as the main reef framework-building organisms (Inoue et al., 2013).

Studies on scleractinian corals indicate that these organisms’ response to anthropogenic-driven climate change depends on the stress source and differs among species and ontogenetic stages (Davies, Marchetti, Ries, & Castillo, 2016; Moya et al., 2012; Thomas et al., 2018). In *Acropora hyacinthus*, heat stress triggers a large and dynamic transcriptomic response characterized by the modulation of different metabolic and cell cycle processes in the early stress response and of spliceosome activity, RNA- and DNA-related metabolic processes, cell stress, and cell metabolism before bleaching onset (Seneca & Palumbi, 2015). A meta-analysis of the transcriptomic response to different stress sources in ten *Acropora* species identified an “environmental stress response” toolkit consisting of genes involved in the response to reactive oxygen species, protein folding and degradation, NF-κB signaling, immune response, and cell death (Dixon, Abbott, & Matz, 2020) that corals exposed to environmental stress consistently modulated. In contrast, ocean acidification does not appear to trigger a pronounced transcriptional response in adult corals colonies (Davies et al., 2016; González-Pech, Vargas, Francis, & Wörheide, 2017) but elicit the modulation of several calcification-related genes in the primary polyps of *Acropora millepora* (Moya et al., 2012).

Despite the observed higher tolerance of octocorals to climate change-induced stress, their physiological response to these stimuli remains poorly studied. Wiens et al. (2000) and Shimpi et al. (2016) observed Heat-Shock Protein overexpression in thermally stressed *Dendronephthya klunzingeri* and *Sinularia* cf. *cruciata* colonies, respectively. Similarly, the sea pen *Veretillum cynomorium* displayed elevated heat-shock protein concentrations upon exposure to thermal stress, but this species did not show increased antioxidant or lipid peroxidation activities (Lopes et al., 2018). To date, aside from these limited data on the octocoral response to environmental stress, no assessment of the transcriptomic response of octocorals exist, and the resilience mechanisms used by these organisms to tolerate and, eventually, outgrow and outcompete less resilient organisms, like stony corals, remain largely unknown.

However, since growth in octocorals requires the deposition of new calcium-carbonate skeletal elements (calcite sclerites, or aragonite fibers in case of the blue corals) to support the colony structurally (Lewis & Vonwallis, 1991), resilience to climate change in this group must be linked to their ability to sustain calcification under challenging environmental conditions (Gabay, Benayahu, & Fine, 2013; Gabay, Fine, Barkay, & Benayahu, 2014; Gómez et al., 2015; Inoue et al., 2013). Indeed, except for the Helioporacea, the octocoral tissues effectively isolate sclerocytes from the surrounding seawater (Gabay et al., 2014), putatively allowing these cells to sustain the expression of transcripts involved in calcification under situations of environmental stress, and the colonies to outgrow and outcompete less resilient organisms. To test this hypothesis, we characterized the skeletal proteome of the common Indo-Pacific gorgonian octocoral *Pinnigorgia flava* and used manipulative experiments and transcriptomics to assess the effect of heat and pH stress on the expression of transcripts involved in calcification in this species. Together, our data provide a first assessment of the transcriptional response an octocoral to environmental stress factors induced by anthropogenically driven climate change and gives insights into the octocoral resilience mechanisms.

## Materials and Methods

### Experimental model and subject details

We used clonal explants of a colony of *Pinnigorgia flava* (Nutting, 1910) kept at 25 °C and a 12:12 h light cycle in a 642 L marine aquarium system at the Department of Earth- and Environmental Sciences, Paleontology and Geobiology, Ludwig-Maximilians Universität München (see below). *Pinnigorgia flava* is a colonial, zooxanthellate soft coral (Octocorallia) largely endemic to the Great Barrier Reef but sporadically found in the coral triangle in SE Asia. The studied colony is cultured in our research aquaria for more than a decade, its exact geographic origin is unknown. It forms non-anastomosing, pinnated light purple colonies with brownish polyps. The sclerites are white to yellow bent spindles, capstans, and rods. Currently, *P. flava* belongs to the family Gorgoniidae.

### Determination of calcification hotspots along the body axis of *P. flava*

We used calcein, a calcium-binding fluorescent dye that permanently incorporates into newly formed skeleton, to investigate the distribution of calcification sites along the body axis of *P. flava* (Holcomb, Cohen, & McCorkle, 2013/2). We incubated three colonies of *P. flava* for 72 h in a glass container with 500ml of a 50µg/ml calcein disodium salt (Sigma-Aldrich) in 0.2 µm filtered artificial seawater (Suppl. Fig. 1). We exchanged the seawater with fresh seawater+calcein every 24 h. After staining, we fixed the colonies in 80% EtOH and stored them at ∼5 °C until further processing.

To assess whether calcification preferentially occurs on the tip of the colonies or, on the contrary, the calcification hotspots occur along the colony body axis, we cut the colonies in top and bottom sections using a sterile scalpel and placed each piece in 1.5 ml microcentrifuge tubes containing one ml of sodium hypochlorite (NaOCl 10%; Fluka). After a three-hour bleach incubation, we rinsed the sedimented sclerites six times with distilled water and stored them in 80% ethanol. We then placed a sample of sclerites onto glass slides and embedded them in Eukitt quick-hardening mounting medium (Fluka Analytical) before covering the sample with a glass coverslip.

We observed the sclerites under epifluorescence (excitation band-pass filter 420-490 nm, barrier long-pass filter 515 nm) on a Leica DMLB microscope coupled to a Leica DFC 480 camera and an I3 filter set. We exposed the stained sclerites for ten seconds and acquired pictures using Leica Application Software “LAS V4.5”. To determine the number of stained sclerites per colony region, we sampled the top and bottom fragments of the colonies and counted stained and total sclerites per visual field at a 100X magnification along one horizontal transect crossing the slide from left to right.

### Proteomic analysis of the skeletal organic matrix of *P. flava’s* sclerites

To determine the skeletal proteome of *P. flava*, we sampled four colonies of about four centimeters in length and incubated them in sodium hypochlorite (5%, Fluka) for 72 h under moderate shaking (30 rpm; IKA Rocker 3D digital). We then rinsed the sedimented sclerites six times with Milli-Q water and dried them at 37°C for 24 h. This procedure yielded approximately 0.75 g of dry sclerites, which we ground with a mortar and pestle before incubating again in sodium hypochlorite (2%) for four hours under moderate mixing (30 rpm). After bleaching, we rinsed the powder six times with Milli-Q water and dried it overnight at 37°C. To dissolve the calcitic mineral, we incubated the dry powder in 10% acetic acid overnight under moderate mixing (20 rpm). We centrifuged the resulting solution at 13,500 rpm for 30 min to separate the acetic-insoluble matrix (AIM) from the acetic-soluble matrix (ASM). To isolate the ASM, we centrifuged (4600 rpm for 70 min at 16°C) the supernatant through 15 mL Amicon ultrafiltration devices with a 3 kDa cutoff membrane and added four volumes of methanol, one volume of chloroform, and three volumes of Milli-Q water to one volume of desalted solution to precipitate the proteins by centrifugation at 5,500 rpm for 15 min (Wessel & Flügge, 1984). After discarding the upper phase, adding three volumes of methanol, and centrifuging the sample at 5,500 rpm for 15 min, we air-dried the resulting ASM pellet and resuspended both the ASM and AIM fractions in 95% Laemmli buffer + 5% β-mercaptoethanol. We used a 1-dimensional sodium dodecyl sulfate-polyacrylamide (SDS-PAGE) gel (Mini-PROTEAN Tetra System, Bio-Rad, USA) to separate electrophoretically the skeletal organic matrix proteins before mass spectrometry. To visualize the extracted SOM protein fractions, we ran an SDS-PAGE for 90 min at 80V, increasing the voltage to 100V after the gel front passed the boundary between the stacking and the resolving gel. We used the Precision Plus Protein Dual Xtra Standard (Bio-Rad, 12 band marker, 2kD-250kD) as a size standard and stained the gel after fixing for 20 min in a fixation solution (50% ethanol, 40% Milli-Q, and 10% acetic acid), washing in 30% ethanol for ten minutes, and in Milli-Q water for ten minutes, with silver nitrate using the Proteo Silver Plus Silver Stain Kit (Sigma-Aldrich, USA). For this, we incubated the gel in sensitizer solution for ten minutes, washed it as described above, and equilibrated it for ten minutes in the silver solution. Before developing, we washed the stained gel for one minute in Milli-Q water and submerged it in a developing solution for five minutes. After stopping the development reaction, we washed the gel for 15 min in Milli-Q water. We used an orbital shaker at 60 rpm for all steps described above. For mass spectrometry, we ran the SDS-PAGE with the six technical replicates of the extracted SOM protein fractions for 40 min at 80V until the protein extracts passed the boundary between the stacking and the resolving gel, and manually excised the bands with a sterile scalpel. We then subjected the isolated proteins to alkylation, reduction, and tryptic digestion (0.1 μg/μl trypsin at 37°C, overnight). We used an LTQ Orbitrap mass spectrometer (Thermo Fisher Scientific, Waltham, Massachusetts, USA) coupled with a Rheos Allegro liquid chromatograph (Flux Instruments GmbH, Basel, Switzerland) to analyze three μl of the digested sample after separation using a self-made column of 75 µm diameter, 15 cm length, C18 particles of 2 µm diameter and 100 Å pore size (Dr. Maisch GmbH, Ammerbruch-Entringen, Germany). We prepared the MS grade mobile phases as follows A) water containing 10% acetonitrile (ACN), B) ACN containing 10% water, each combined with 0.1% formic acid. We used the following gradient: 40 min (0-23% B), 40 min (23-85% B), five minutes (85-100% B), 25 min (100% B), three minutes (100-0% B) and 20 min (0 % B) for re-equilibration, and a constant 40 µl/min flow at RT (22°C). We used the programs Xcalibur 2.0 (Thermo Fisher Scientific Inc., 30 Waltham, USA) and MaxQuant Version 1.5.2.8 (Cox andf Mann 2008, doi: 10.1038/nbt.1511) to acquire and analyze the MS/MS data, respectively. We estimated protein abundances as relative iBAQ values (Schwanhäusser et al., 2011) of the detected SOM proteins using the MaxLFQ algorithm (Cox et al., 2014). We filtered the LC-MS/MS results to remove hits from known conventional contamination sources using the common Repository Adventitious Proteins (cRAP) database before mapping the peptides against a transcriptome reference of *P. flava* (Conci, Wörheide, & Vargas, 2019). We translated the transcripts matching proteins present in the SOM fractions and annotated them against the UniProtKB database using blastp (https://blast.ncbi.nlm.nih.gov/) and SignalP 4.0 (Petersen, Brunak, von Heijne, & Nielsen, 2011) to predict the presence of signal peptides, transmembrane regions, and GPI anchors within the predicted protein sequences. Additionally, to assess the distribution of the detected *P. flava* SOM proteins among cnidarians, we screened a database composed of 120 transcriptomes from representatives of this phylum using *P. flava’s* SOM proteins as queries for the search (for details see Eitel et al., 2018).

### In vivo experiments

We used a 360 L marine aquarium under a 12:12 h light cycle controlled by GHL Mitras LX 6200-HV LED lights that yielded ten kLux (PAR = 250 μmol/m^2^s) at the water surface. Based on hourly measurements over one year (2017), the system’s average water temperature and pH are 24.92 ± 0.24 °C and 8.30 ± 0.14, respectively. Based on weekly measurements over one year (2017), the average PO_4_^-3^, NO_2_^-^ and NO_3_^-^ concentrations in the water are 0.092 ± 0.071 mg/L, 0.014 ± 0.072 mg/L, 2.681 ± 3.882 mg/L; the concentration of NH_3_/NH_4_^+^ in the water was consistently below detection (i.e. < 0.05 mg/L). Weekly or daily monitoring in subsequent years revealed that these values are stable and can be taken as a baseline for the system. We measured the PO_4_^-3^, NO_2_^-^, NO_3_^-^ and NH_3_/NH_4_^+^ concentrations every other day during the experiments. The measured concentrations did not differ from the baseline concentrations for the tank.

To assess the effect of heat stress on *P. flava*, we randomly assigned nubbins (n = 18) to six aquaria (10L) filled with approximately six liters of artificial seawater and partially immersed in the 360 L aquarium described above. Water evaporation in the ten-liter tanks was compensated every day with water filtered by reverse osmosis. We used a submersible water pump (300 L/h; Eheim, Germany) to provide adequate water mixing in each tank. For the temperature experiment, after an acclimation period of four days, we randomly selected three tanks and gradually increased the water temperature to 29-30°C for five days (∼1°C per day) using 50 W water heaters (Eheim, Deizisau, Germany). We then kept the *P. flava* colonies at 29-30°C for three days. Although the bleaching threshold of *P. flava* is not known, considering that the colony used for our experiments grows since >ten years at a 25 °C, the final water temperature (i.e., 29-30°C) achieved during the experiment lies >four °C over the mean water temperature this coral is exposed to. Thus, the increase in temperature from 25°C to 29-30°C is expected to cause a heat stress response in *P. flava* without necessarily causing its death. In scleractinian corals, for example, bleaching can be induced by heat stress only about 1°C above the average summer maximum temperature. During the experimental period, we monitored water pH, density, and temperature (manually twice per day) every day (Suppl. Tables 1-3). We also measured the nutrient profile in the ten-liter tanks at the end of the experiment. The observed physicochemical values were similar to those of the main tank, and most nutrients were below the detection limit of the tests used. We also visually monitored the colonies, which had their polyps everted and did not show any visible signs of bleaching during the experiment. In this regard, the bleaching threshold is not known for *P. flava*. At the end of the experiment, we cut octocorals in two (i.e., lower and upper) sections using sterile scissors and flash-froze them in liquid nitrogen before storing them at -80°C until further processing.

We assessed the effect of seawater acidification on twelve colonies of *P. flava* distributed in two 30 L tanks (control and treatment) connected to a 320 L salt-water “mother” tank. To lower (0.1 pH units drop per day) the seawater pH, we pumped CO_2_ into the treatment tank (see details in González-Pech et al., 2017) to achieve a pH of 7.8 (from a starting value of 8.2). We kept the pH stable for three days and lowered it again to 7.6 over three days. After reaching pH = 7.6, we allowed the octocorals to acclimate for three days before decreasing the pH to its final value of 7.3. Gomez et al. (2015) showed that *Eunicea fusca* grows under a broad range of seawater pH values but possibly experience some level of decalcification at pH = 7.1. Therefore, we used a pH = 7.3 to evaluate *P. flava*’s response to seawater acidification as this value represents the lowest reported pH value at which octocorals continue growing without sclerite decalcification. We maintained this pH for two months using an automatic pH computer and monitored it throughout the experiment using a PCE-PHD 1 datalogger (PCE Instruments, Germany). The pH of the “mother” and control tank was 8.2 throughout the experiment. The temperature and nutrient levels in the “mother” tank were as detailed above. During the entire experimental phase, the coral colonies showed everted polyps and no signs of stress (e.g., bleaching). At the end of the experiment, we cut the octocorals at the base using sterile scalpels and then flash-froze them in liquid nitrogen before storing them at -80°C until further processing.

### Analysis of differential gene expression

We extracted total RNA from the upper section of the octocorals using the Direct-zol RNA MiniPrep kit (Zymo Research) following the manufacturer’s protocol. We assessed the purity and integrity of the RNA extracts using a Nanodrop ND-1000 spectrophotometer (Thermo Fisher Scientific, USA) and a Bioanalyzer 2100 (Agilent Inc., USA). In general, all samples showed 260/230 and 260/280 ratios > 2.0 and RIN values above 8.5. We used Lexogen’s SENSE Total RNA-Seq Library Prep Kit according to the manufacturer’s instructions to generate Illumina-ready, stranded transcriptome libraries for all extracted samples. We sequenced the libraries (50bp Pair-End) in an Illumina HiSeq2000 (see Suppl. Table 4), discarded reads with an average quality < Q28, stretches of poly-{ATCG} or Illumina adapters using the program *bl-filter-illumina* from the BioLite suite (Howison, Sinnott-Armstrong, & Dunn, 2012), and mapped them to an available *P. flava* holobiont reference transcriptome (Conci et al., 2019) using Salmon (Patro, Duggal, Love, Irizarry, & Kingsford, 2017). We truncated the resulting (holobiont) pseudocounts matrix and analyzed it using DESeq2 (Love, Huber, & Anders, 2014) to determine differentially expressed genes (DGE) in 1) control vs. heat stress treated and 2) control vs. pH stress treated octocorals. For each stress treatment (i.e., temperature and pH stress) we separated host and symbiont DGEs in R using lists of host and symbiont transcripts determined using PsyTrans (REFS). We used the obtained list of host DEGs for GO-term enrichment analyses with TopGO (Alexa & Rahnenfuhrer, 2016) using a Fisher test on nodes with at least five annotations. To summarize the resulting list of significantly enriched GO-terms, we used the R package GOSemSim (Yu et al., 2010) to calculated the pairwise semantic similarity (Wang) between GO-terms and clustered (Ward) the enriched GO-terms into groups of semantically related GO-terms. To assign a representative GO-term to each cluster of semantically related, significantly enriched GO-terms in the different stress treatments, we took advantage of the hierarchical structure of the gene ontology and retrieved subgraphs including the GO-terms in each cluster. These subgraphs show how semantically similar GO-terms relate to each other in the more broad gene ontology directed acyclic graph. We used these subgraphs graphs to find GO-terms from which other GO-terms in a cluster derived (i.e., “ancestor” GO-terms) and used these more general GO-terms as cluster descriptors.

## Results

### Calcification in *P. flava* occurs along its entire body axis

Visual inspection of stained sclerites in the top vs. bottom parts of *P. flava* colonies revealed that this species produces new sclerites in both the top and bottom sections of the colony, with a slight but significant increase in the proportion of stained sclerites toward the tips of the colony (Welch two-sample t=-2.17, df=92.531, p=0.034). Examination of whole colonies sections revealed that calcification usually occurs at the polyps (Fig. 1). Despite the increased number of stained sclerites at the colony’s top, we could not detect any evident calcification hotspots along the colony axis.

**Figure 1.**
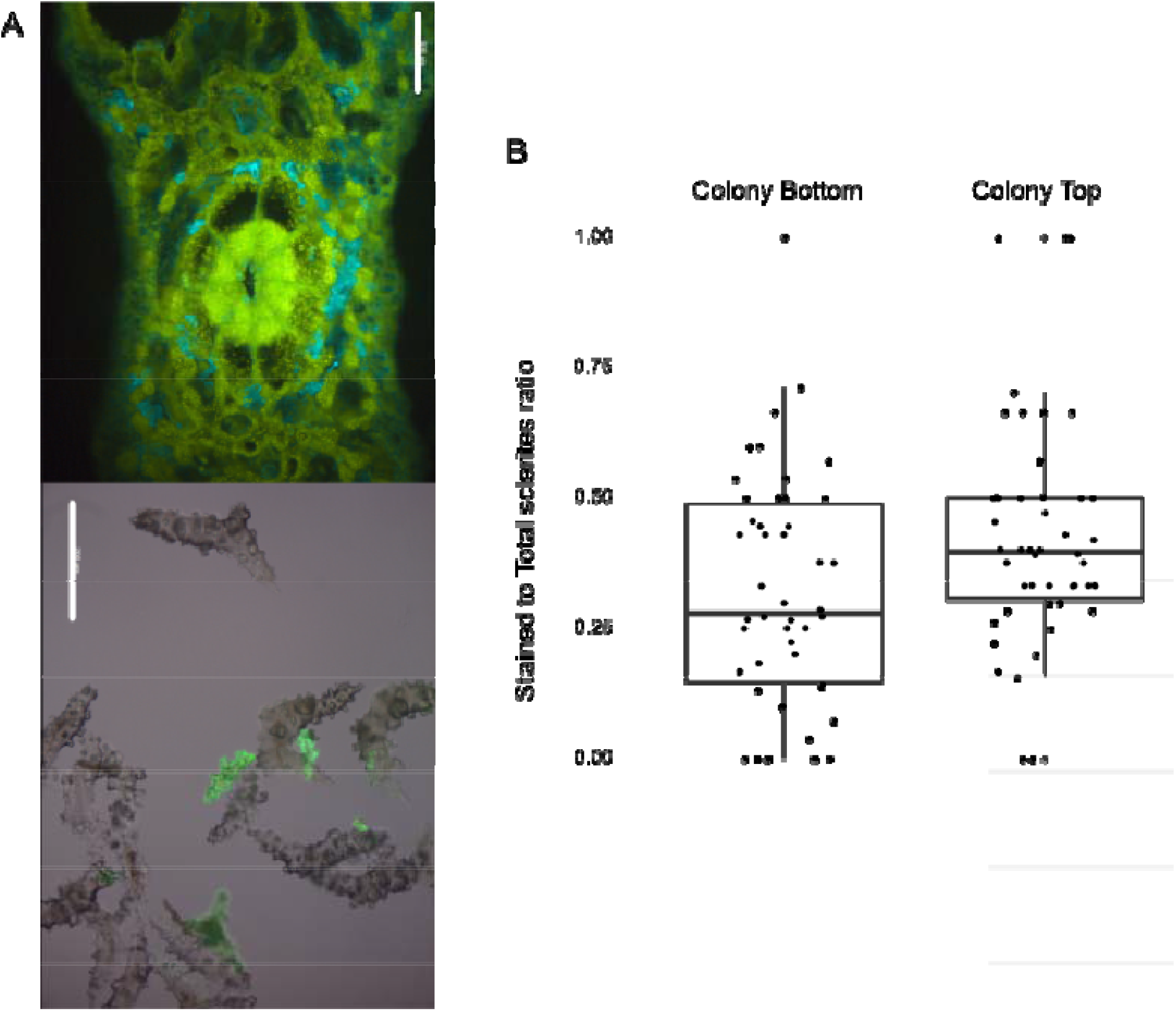
Calcification in *Pinnigorgia flava*. **A:** *In situ* (blue hue used to differentiate calcein staining from tissue autofluorescence) and isolated calcein-stained sclerites (green fluorescent), note the distribution of newly deposited CaCO_3_ (lighter green dots) around the polyp (figure center). Both scale bars = 200μm. **B:** ratio of calcein-stained to total number of sclerites in bottom vs. top colony parts. Welch two-sample t=-2.17, df=92.531, p=0.034.

### Taxonomically widespread and restricted proteins form the skeletal proteome of *P. flava*

After filtering potential known contaminants (e.g., keratins, trypsin), we identified a total of 27 transcripts as the skeletal organic matrix (SOM) proteome of *P. flava*. Label-free quantification (LFQ) revealed 14 proteins with a higher abundance in the acetic-insoluble matrix (AIM), seven proteins exclusive to the acetic-soluble matrix (ASM), and five proteins shared by both fractions (Fig. 2). Biochemically, seven of the detected SOM proteins are membrane-bound, containing either transmembrane domains (n = 3), GPI anchors (n = 3), or both (n = 1). About 46% (n = 13) of the SOM proteins detected contain signal peptide motifs, indicating that these proteins are secretion targets (Fig. 2). We found the SOM proteome to contain several components of the extracellular matrix. Among them are collagens and laminin, glycoproteins with calcium-binding domains (i.e., osteonidogen and agrin), and proteins likely involved in cell adhesion (i.e., otoancorin, hemicentin, and von Willebrand factor type A (VWA) domain bearing proteins). Finally, we found a protein similar to galaxin, two proteins enriched in aspartic-acid residues (acidic, predicted pI ∼ 3.4) with no significant UniProt blast hits, and several other proteins with diverse functionalities (e.g., disulfide isomerase or protease activity). According to their iBAQ, six proteins had abundances >5% in the skeletal proteome of *P. flava*. The two acidic proteins ranked first and third, osteonidogen and collagen were the second and fourth most abundant proteins, and galaxin and agrin ranked fifth and sixth in abundance. These six proteins account for 90% of the total iBAQ-derived skeletal protein abundance (Fig. 2). On average, the remaining proteins accounted for only 0.5% (±0.6%) of the total iBAQ-derived abundance.

**Figure 2.**
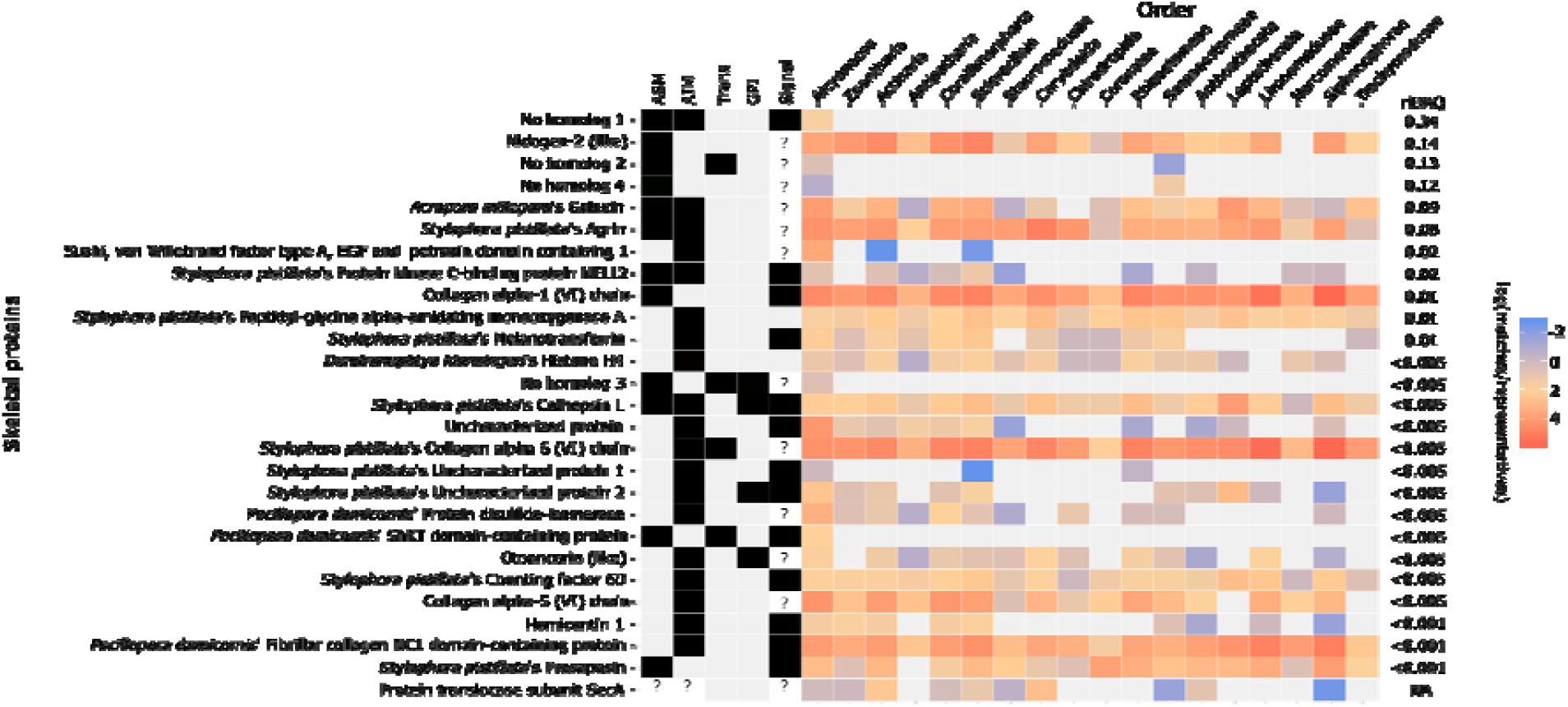
Presence/absence (black/gray rectangles, respectively) in the acid-soluble (ASM) and insoluble (AIM) matrix, biochemical properties, distribution among cnidarians, and abundance in the skeletal proteome of 27 proteins present in the skeleton of *P. flava*. Trans: Presence of transmembrane domains. GPI: Presence of GPI anchors. Signal: Presence of signal peptide.A question mark (?) indicates that the particular property could not be determined, e.g., due to the availability of only partial sequences.

In terms of their taxonomic distribution within Cnidaria (Fig. 2), similarity searches revealed that the skeletal proteome of *P. flava* includes proteins with a widespread distribution in this group, such as all detected collagens, agrin, osteonidogen, galaxin, and several enzymes, among others. Other proteins, like hemicentin or the protein kinase Nell 1, displayed a patchier occupancy and were found mostly in other anthozoans. Finally, only a few SOM components, namely the two acidic proteins found, a serine protein-kinase receptor and a protein similar to laminin, had a restricted taxon occupancy with significantly similar proteins found almost exclusively in other octocorals (i.e., Order Alcyonacea).

### Climate change-driven stress does not modulate the calcification toolkit of *P. flava*

Exposure to sublethal, high-temperature (∼29-30°C, ∼5°C above long-term average in the aquaria) seawater resulted in the modulation of 751 host transcripts (Benjamini-Hochberg corrected *p* < 0.05), 108 transcripts upregulated (log2 fold change ≥ 1) and with 221 transcripts downregulated (log2 fold change ≤ -1) in heat-treated colonies (Fig. 3A, Suppl. Fig. 2, and Suppl. Table 5). We found two representatives of the heat-shock protein 70 family among the transcripts upregulated by heat stress (Fig. 3B). Among calcification-related proteins, heat stress caused the downregulation of one carbonic anhydrase and one galaxin, not affecting the remaining 25 calcification-related proteins previously identified in octocorals (Conci et al., 2019). Also, heat stress did not significantly change the expression of any of the 27 transcripts encoding SOM proteins in *P. flava* (Fig. 3B).

**Figure 3.**
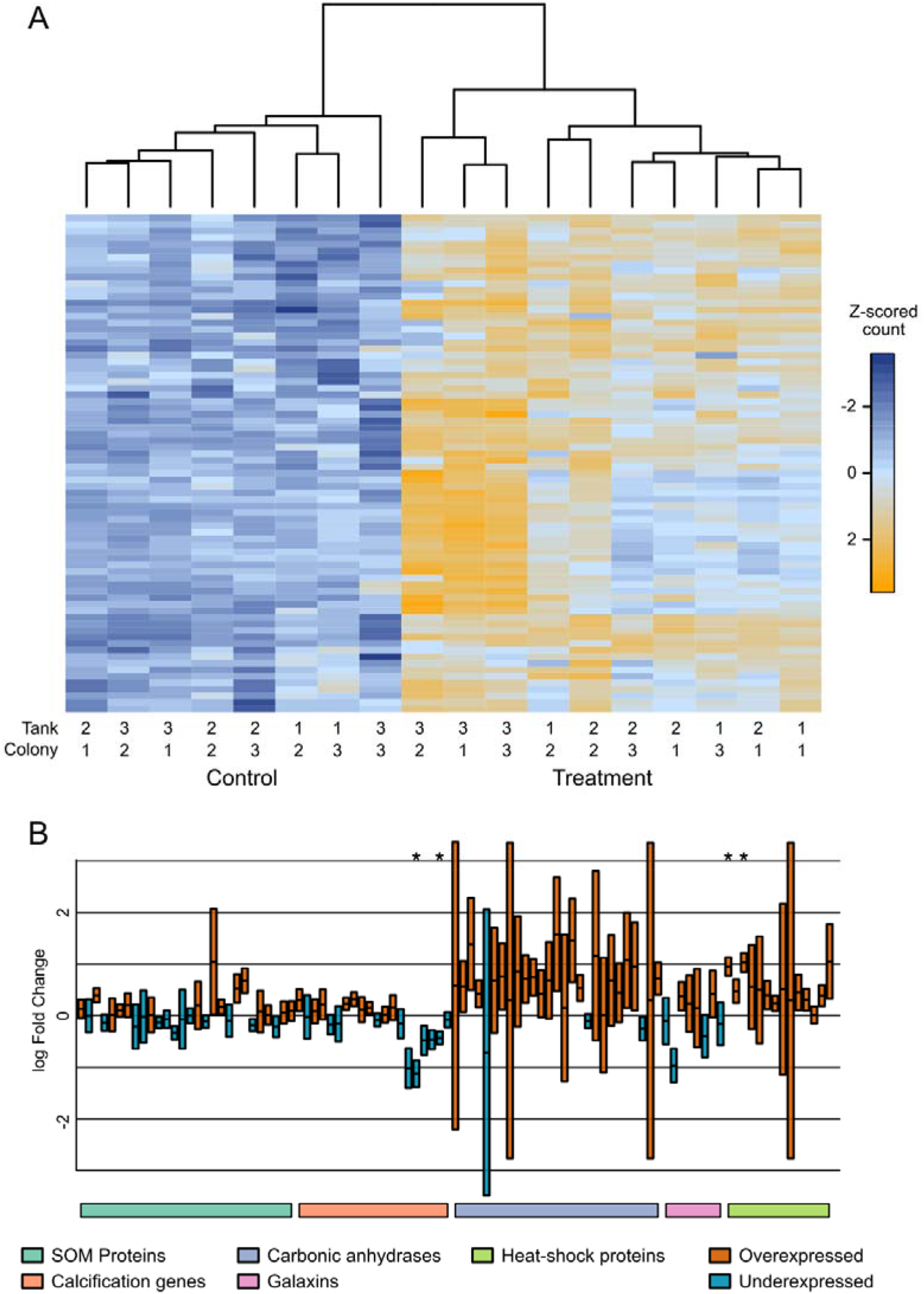
**A:** Expression (variance stabilized, z-scored transformed counts) of 76 transcripts modulated in *P*.*flava* colonies exposed to control and heat stress. For visualization purposes only transcripts with log2 fold change ≥ |1| and p < 0.01 were included in the heat plot. **B:** Change in gene expression of 27 SOM proteins, calcification related proteins *sensu* Conci et al. (2019), carbonic anhydrases, galaxins, and heat-shock proteins in *P. flava* colonies exposed to heat stress. The asterisks (*) indicate transcripts significantly differentially expressed in control vs. treatment samples.

The GO-term enrichment analysis of the set of upregulated transcripts during heat stress resulted in seven clusters of semantically similar, significantly enriched Biological Process (BP) GO-terms (Fig. 4 and Suppl. Fig. 3-16). The representative GO-terms of these clusters were related to processes of “programmed cell death” (GO:0012501) and the “regulation of apoptotic processes” (GO:0042981), “response to stress” (GO:0006950) and “adaptive immune responses” (GO:0002250), protein folding (several GO-terms, see S. Materials), redox and metabolic processes, peptide secretion and epidermis development. Detailed information about the GO-terms included in each cluster, the values of the Fisher-test, and the information content of each GO-term in the clusters is provided in the Suppl. Tables 6-12. We found seven clusters of semantically similar, significantly enriched BP GO-terms among transcripts downregulated by heat stress (Fig. 5 and Suppl. Fig. 17-30). These clusters were related to the “cellular response to (increased) oxygen levels” (GO:0071453), the “regulation of cellular response to oxidative stress” (GO:1900407), “cartilage development” (GO:0051216), including terms related to bone morphogenesis, “immune response” (GO: 0006955), the “regulation of necrotic death” (GO:0010939), and more miscellaneous GO-terms such as fatty acid elongation (GO:0019367 and GO:0019368) and secondary metabolic processes. Detailed information about the GO-terms included in each cluster, the values of the Fisher-test, and the information content of each GO-term in the clusters is provided in the Suppl. Tables 13-19.

**Figure 4.**
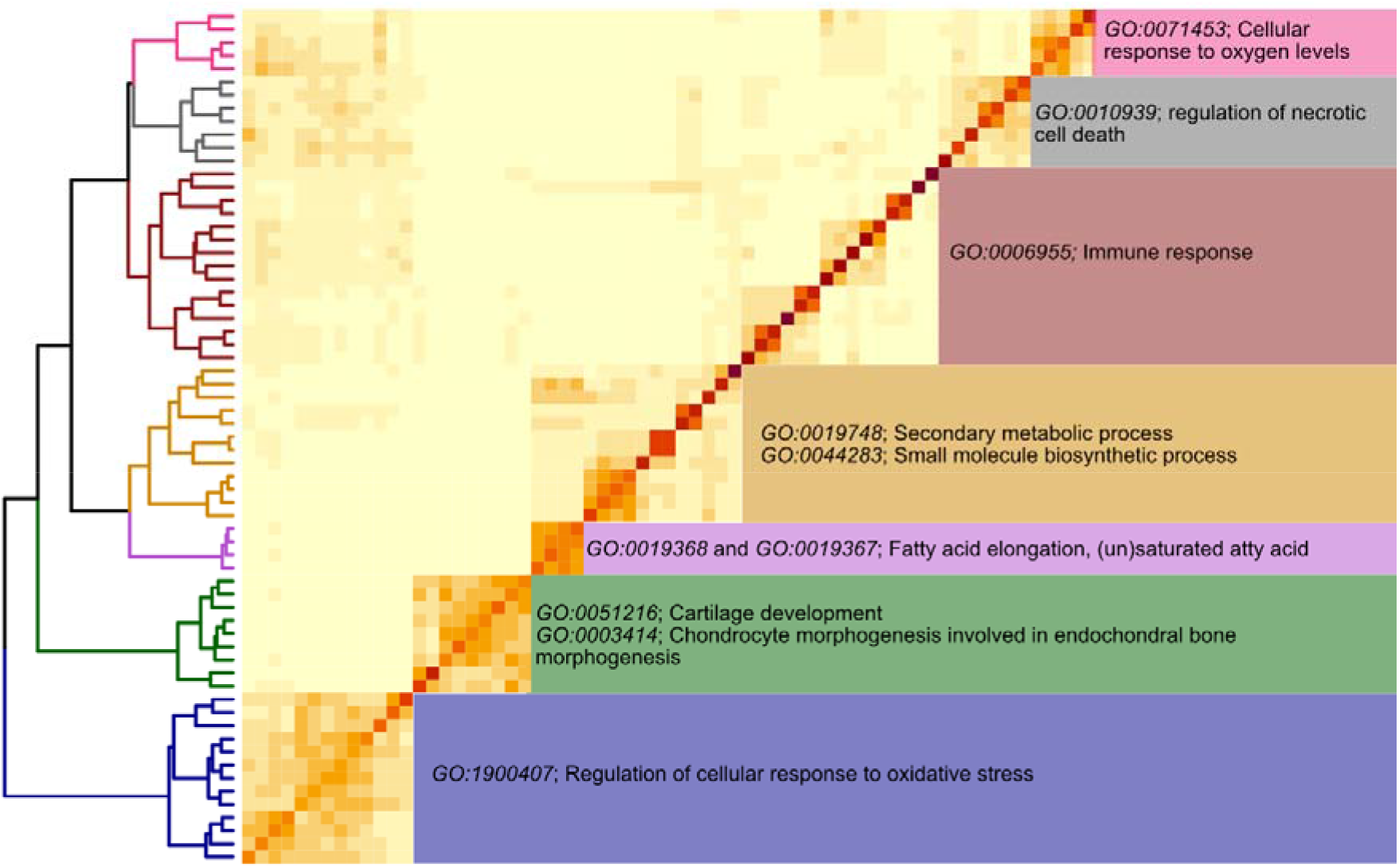
Clusters of semantically similar, significantly enriched Biological Process GO-terms found in the set of overexpressed genes under heat stress. Characteristic GO-terms are annotated to the right of each cluster.

**Figure 5.**
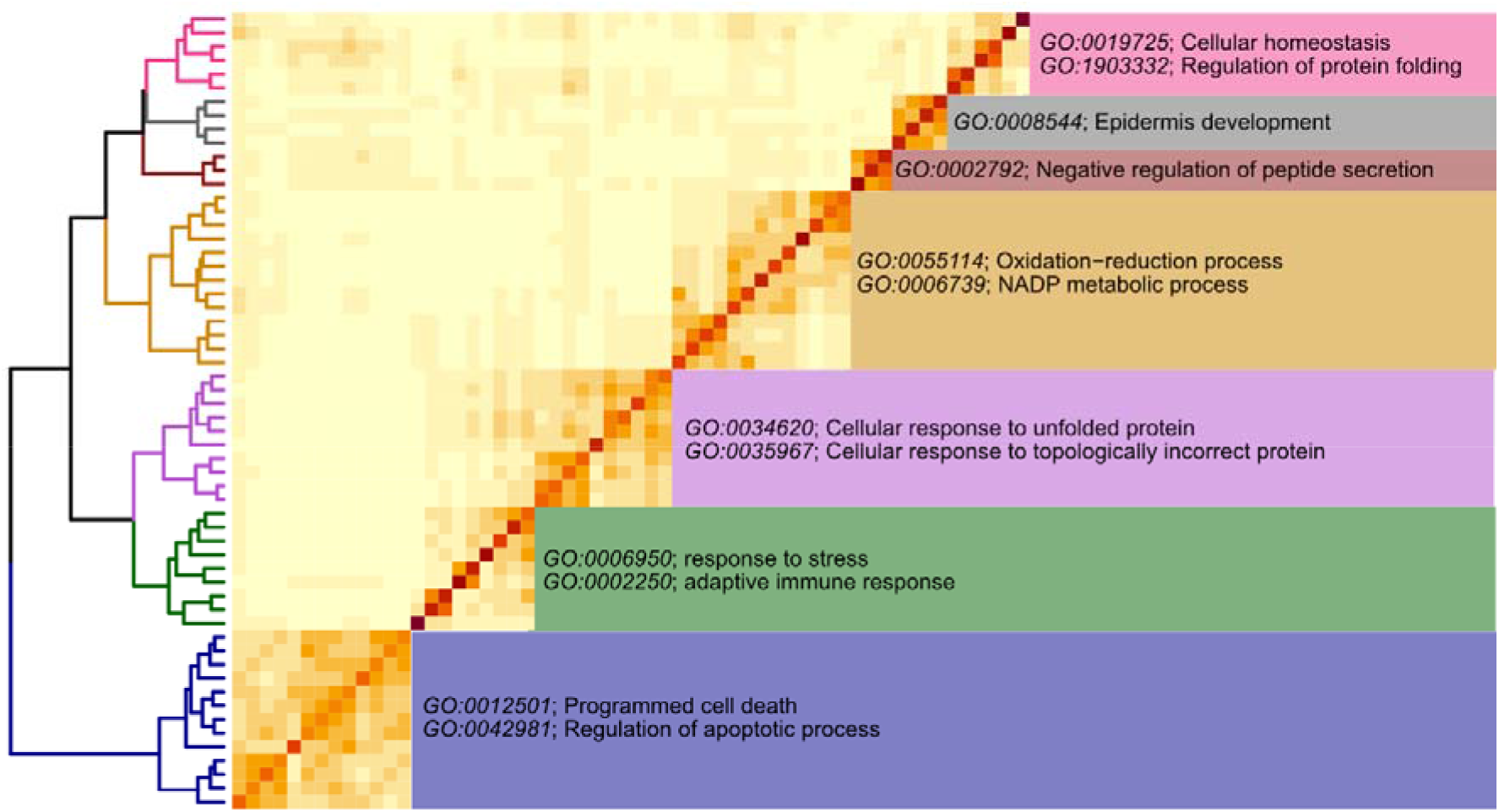
Clusters of semantically similar, significantly enriched Biological Process GO-terms found in the set of underexpressed genes under heat stress. Characteristic GO-terms are annotated to the right of each cluster.

In contrast to the response of the coral, heat stress resulted in the modulation of only 19 transcripts in *P. flava*’s symbionts; nine of these transcripts had a log2 fold change ≥ 1 and only one transcript had log2 fold change ≤ -1 (Suppl. Table 20).

Contrary to heat stress, pH stress did not trigger a stark stress response in *P. flava*, only causing the significant (*p* < 0.05) downregulation of a set of 70 host transcripts, 61 of which had a log2 fold change ≤ -1 (Fig. 6A, Suppl. Fig. 31, and Suppl. Table 21). We found one calcification-related, uncharacterized skeletal matrix protein and two SOM-encoding transcripts, namely osteonidogen -- the second more abundant SOM protein-- and a prosaposin-homolog downregulated in pH stressed corals (Fig. 6B). We did not detect changes in the expression of stress-related proteins from the Heat-Shock Protein family (Fig. 6B). Our GO-term enrichment analysis resulted in nine clusters of semantically similar, significantly enriched BP GO-terms (Fig. 7 and Suppl. Fig. 32-49). These clusters were related to the “regulation of metalloendopeptidase activity” (GO:1904683), “protein localization to synapse” (GO:0035418), “regulation of bone mineralization” (GO:0030500) and “regulation of biomineral tissue development” (GO:0070167), “terpenoid metabolic process” (GO:0006721), “interaction with host” (GO:0051701), “response to BMP” (GO:0071772) and “transmembrane receptor protein serine/threonine kinase signaling pathway” (GO:0007178), “regulation of leucocyte proliferation” (GO:0070663) and “adaptive immune response” (GO:0002250). We found one cluster grouping largely unrelated GO terms difficult to summarize with a single representative GO term. Detailed information about the GO-terms included in each cluster, the values of the Fisher-test, and the information content of each GO-term in the clusters is provided in the Suppl. Tables 22-30.

**Figure 6.**
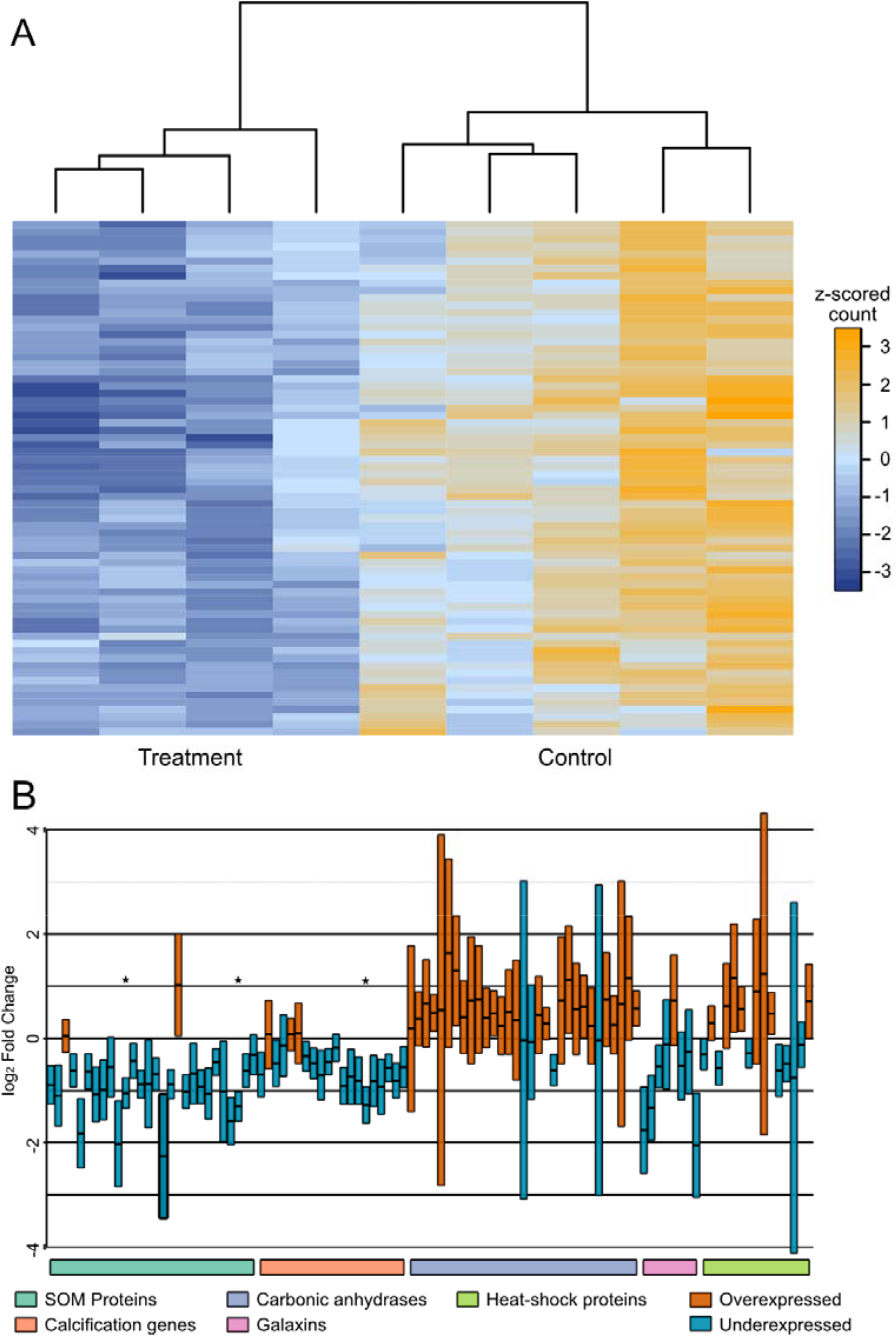
**A:** Expression (variance stabilized, z-scored transformed counts) of 70 transcripts modulated (p < 0.05) in *P*.*flava* colonies exposed to control and low pH stress. **B:** Change in gene expression of 27 SOM proteins, calcification related proteins *sensu* Conci et al. (2019), carbonic anhydrases, galaxins, and heat-shock proteins in *P. flava* colonies exposed to low pH stress. The asterisks (*) indicate transcripts significantly differentially expressed in control vs. treatment samples.

**Figure 7.**
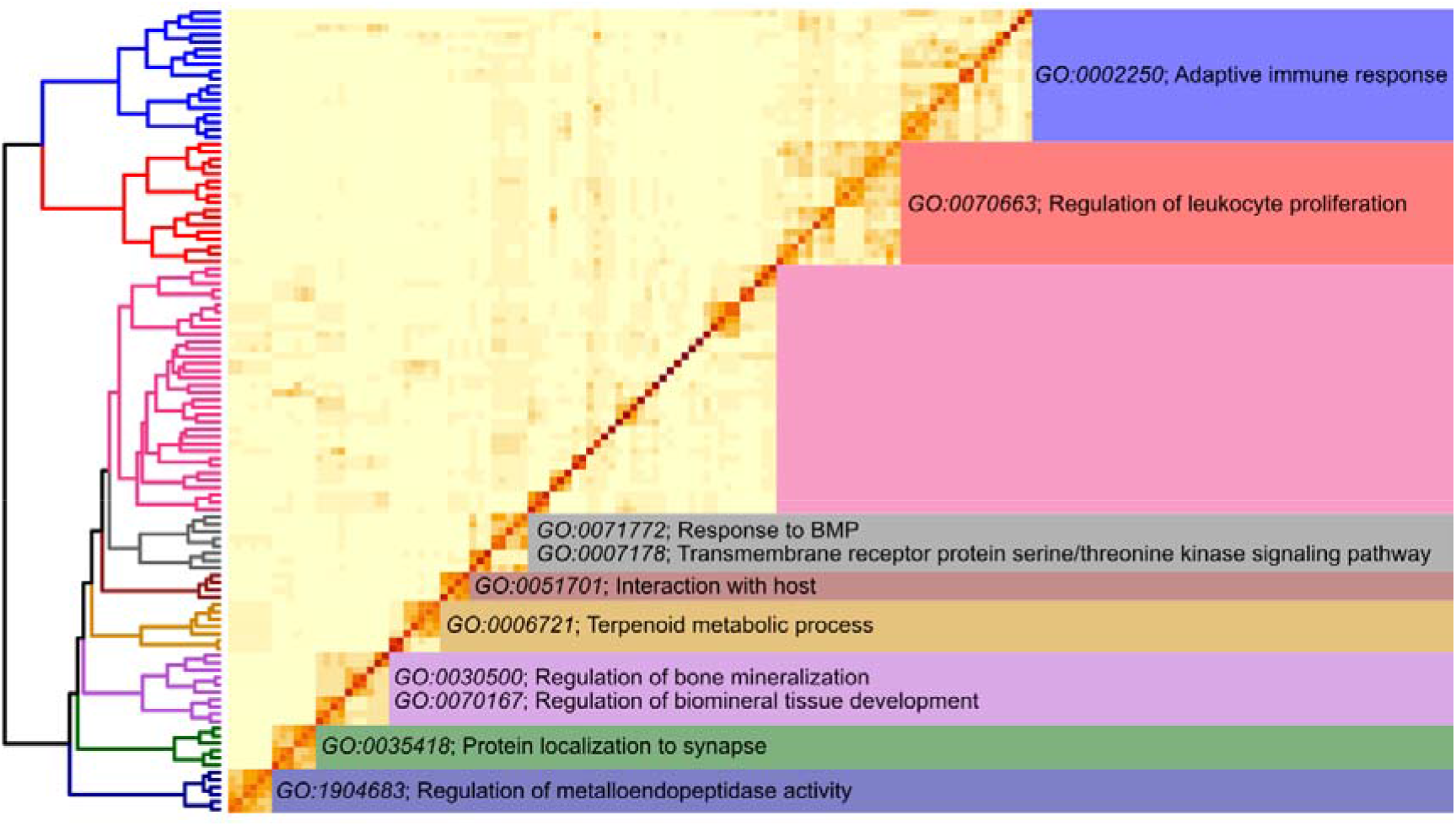
Clusters of semantically similar, significantly enriched Biological Process GO-terms found in the set of underexpressed genes under pH stress. Characteristic GO-terms are annotated to the right of each cluster.

pH stress did not result in the upregulation of any transcript in *P. flava* or the modulation of any transcripts in *P. flava*’s symbionts.

## Discussion

Sclerite growth in octocorals involves the synthesis by sclerocytes of skeletal organic matrix (SOM) proteins and their transport to a sclerite-forming vacuole where primordia, mostly made of irregularly shaped CaCO_3_ crystals, form (Kingsley, 1984). Sclerite primordia continue growing by the deposition of more regular crystals in an extracellular space created by multiple sclerocytes (Goldberg & Benayahu, 1987). In *P. flava*, our results indicate that the deposition of new sclerites occurs throughout the colony axis, increasing toward the colony tips. This pattern of calcification is similar to that of *Leptogorgia virgulata*, one of the few other octocoral species where data on calcification dynamics exist (Kingsley & Watabe, 1989). It indicates that active sclerocytes intersperse along the colony axis and that under normal conditions, we should not expect spatial differences in the expression of SOM-encoding transcripts in this species. Hence, the exposure of colonies of *P. flava* to stress factors affecting the expression of SOM encoding transcripts should impair the deposition of new sclerites at the colony level. The collapse of the calcification machinery, in turn, would result in a reduced ability of octocorals to sustain growth and eventually outcompete other reef organisms, like stony corals, in stressful environments.

The skeletal proteome of *P. flava* is similar to previously characterized proteomes in octo- and scleractinian corals (Conci, Lehmann, Vargas, & Wörheide, 2020; Drake et al., 2013; Ramos-Silva et al., 2013) and contains a mixture of taxonomically restricted and widespread elements as reported in other octocorals (Conci et al., 2020). This trend is particularly evident among the most abundant proteins in the SOM. Among those, galaxin and the two acidic proteins detected had a very narrow taxonomic distribution. Galaxins mainly occur within Cnidaria (Conci et al., 2019), and acidic proteins appear to evolve rapidly, species-specific proteins difficult to assign to cnidarian orthology groups using similarity searches (Conci et al., 2020). Our results are consistent with that observation, as the two acidic proteins found do not show significant similarity to other proteins deposited in public databases such as SwissProt, and even within Cnidaria, only matched proteins found in other octocorals. Among the group of taxonomically widespread proteins, we found typical components of animal basement membranes (Erickson & Couchman, 2000) with a broad distribution within Cnidaria and generally within Metazoa. Indeed, three of the six most abundant proteins, namely agrin, collagen, and osteonidogen, are glycoproteins involved in modulating cell-extracellular matrix interactions (Erickson & Couchman, 2000). Among vertebrates, agrin participates in synaptogenesis (Kröger & Schröder, 2002). Specifically, this protein coordinates the development of neuromuscular junctions, stabilizing and aligning the pre- and postsynaptic apparatuses of neurons and muscle fibers, respectively. It also triggers the differentiation of neuron growth cones into presynaptic terminals capable of calcium-dependent neurotransmission (Kröger & Schröder, 2002; Ruegg & Bixby, 1998). Its presence in the SOM of sclerites may imply a similar role in calcification, aligning the sclerocytes around sclerite primordia and triggering membrane differentiation to allow for calcium secretion into the calcifying space. In line with this reasoning, in addition to biomineralization-related GO-terms, colonies exposed to low pH stress modulated transcripts enriched in GO-terms, pointing to the active regulation of processes involving the localization of proteins to synapses. We did not observe any apparent effect of the pH treatment on colony behavior that suggested colonies could not react to mechanical stimuli. Thus, we propose interpreting the localization of proteins to “synapses” in a biomineralization context as the localization of proteins to calcification sites, which, given the presence of presynaptic proteins, appear to be related to some extent synapses. Although speculative at this point, the analysis of single-cell transcriptomes of octocorals exposed to low pH seawater can help disentangle this stress’ effect on different octocorals cell types (e.g., neurons vs. sclerocytes). In the case of osteonidogen, mammalian osteocytes and osteoclasts overexpress this protein (Barshis et al., 2013; Ramos-Silva et al., 2013), suggesting a direct role in calcification in that group and a possible involvement in calcification in octocorals.

To respond more rapidly to and survive episodes of environmental stress, resilient stony corals constitutively upregulate components of the coral cell death and immune pathways and genes involved in response to stress, like heat-shock proteins (Barshis et al., 2013). The concomitant downregulation of genes involved in calcification observed during environmental stress in these organisms (Barshis et al., 2013; Ramos-Silva et al., 2013) suggests that the transcriptional frontloading of the stress response toolkit comes at the expense of the coral calcification machinery and could lead to its collapse. Accordingly, calcification in colonies of *Siderastrea siderea* exposed to ocean acidification and warming shows a parabolic response, driven mainly by the abrupt drop in calcification rates under more extreme environmental regimes (Inoue et al., 2013; Ruzicka et al., 2013). Our results revealed a similar response of *P. flava* to heat stress, indicating a degree of commonality in the transcriptomic response of stony and soft corals to heat stress. Whether the octocorals’ reaction to heat stress is generally similar to that of stony corals remains to be investigated in a larger sample of octocoral species to assess the universality of the pathways modulated during environmental stress.

Despite the observed similarities in the transcriptional response of the stress toolkit of stony corals and octocorals exposed to heat stress, the calcification machinery of these two groups appear to react differently to adverse environmental conditions. Contrary to stony corals, our results indicate that *P. flava* can sustain the production of all the molecules necessary for the formation of new sclerites during stress events. In this regard, we detected only a mild effect of environmental stress on the calcification toolkit of *P. flava*, with heat stress affecting the expression of a galaxin and a carbonic anhydrase and seawater pH stress modulating the expression of osteonidogen and a prosaposin homolog. Galaxins, carbonic anhydrases, and osteonidogen are calcification-related genes in corals and other organisms, and mutations in human prosaposin cause Gaucher’s disease, a disorder characterized by skeleton deterioration (Vaccaro et al., 2010). The reported linear decrease in octocoral calcification rates under reduced seawater pH (Gómez et al., 2015) is consistent with the hypothesis of a decoupling between the calcification and stress-response toolkits of octocorals, as it indicates that the observed decrease in calcification is mainly driven by the environment, not by a transcriptional response of the octocoral calcification machinery to the adverse conditions. Under long-term adverse conditions, decoupling the physiological response to stress from calcification may give octocorals a competitive advantage over other species, like stony corals, adapted to respond better to episodic stress and lead to the community shifts observed in many reef locations (Inoue et al., 2013a; Ruzicka et al., 2013). Characterizing the transcriptional response to environmental stress of a broader taxonomic sample of octocorals will determine whether the calcification toolkit of these organisms is transcriptionally resilient.

In summary, our results provide new insights into the octocoral stress response to environmental stress. They suggest that the calcification and the stress-response toolkits of soft and stony corals react differently to stress episodes. This difference likely determines the somewhat contrasting response to stress observed in these groups. Anthropogenic-induced global climate change will undoubtedly impact future marine communities in unprecedented ways (Hughes et al., 2019). Processes such as acclimatization and adaptation (Palumbi, Barshis, Traylor-Knowles, & Bay, 2014), acting at organismal and population levels, and phenomena affecting the community, like ecological memory (Hughes et al., 2019), shape the response of coral reefs to these environmental pressures. Our results suggest that, compared to stony corals, octocorals use different gene regulation strategies to face climate change. Thus, understanding the diversity of molecular mechanisms involved in resilience and their regulation in different reef organisms is pivotal to predicting the future of the world’s coral reefs.

## Acknowledgments

We thank Gabriele Büttner for support during laboratory work and Dr. Peter Naumann for supporting the aquarium facilities of the Section of Geobiology and Paleobiology of the Dept. of Earth & Environmental Sciences. We would also like to thank René Neumaier for our High-Performance Computing system’s design, administration, and support; our work would be impossible without his careful and detailed work. The Deutsche Forschungs Gesellschaft funded this study through the grants Va1146-2/1 and Wo896-18/1. GW acknowledges funding through the LMU Munich’s Institutional Strategy LMUexcellent within the framework of the German Excellence Initiative. SV thanks N. Villalobos T., M. Vargas V., S. Vargas V. and S. Vargas V. for their support.

## Data availability statement

- Raw sequence reads: European Nucleotide Archive project PRJEB38757, accession numbers ERS4652944-ERS4652972.
- Raw MS/MS data files: https://data.ub.uni-muenchen.de/189/
- Transcriptome assembly, proteome annotations, count matrix, scripts, and other data files used in this study:https://gitlab.lrz.de/palmuc/pinnigorgia_resilience

## Author Contributions

**SV** Conceptualization, Methodology, Software, Validation, Formal Analysis, Investigation, Resources, Data Curation, Writing - Original Draft, Visualization, Supervision, Funding Acquisition. **TZ** Investigation, Formal Analysis. **NC** Investigation, Formal Analysis, Writing - Review & Editing. **ML** Investigation. **GW** Resources, Writing - Review & Editing, Funding Acquisition.

