## Supplementary Tables for "Transcriptional response of the calcification and stress response toolkits in an octocoral under heat and pH stress": Supplementary_Tables_1-3.pdf

Supplementary Table 1: Water temperature (°C) measured during the heat stress experiment.

| | Morning/Evening temperature (°C) measured on experimental day | | | | | | |
|---|---|---|---|---|---|---|---|
| Tank | 1 | 2 | 3 | 4 | 5 | 7 | 8 |
| Control 1 | 25.3/25.3 | 25.4/25.3 | 25.3/25.3 | 25.4/25.3 | 25.4/25.3 | 25.4/25.4 | 25.4/25.4 |
| Control 2 | 25.4/25.3 | 25.4/25.3 | 25.3/25.3 | 25.4/25.3 | 25.4/25.4 | 25.4/25.4 | 25.4/25.4 |
| Control 3 | 25.3/25.3 | 25.4/25.3 | 25.3/25.3 | 25.4/25.3 | 25.4/25.3 | 25.4/25.4 | 25.4/25.4 |
| Treatment 1 | 26.1/27.3 | 27.4/27.4 | 27.7/27.4 | 27.4/27.6 | 28.7/28.5 | 28.7/28.6 | 28.6/29.2 |
| Treatment 2 | 26.3/27.5 | 27.7/27.7 | 27.7/27.7 | 27.8/27.9 | 28.4/28.0 | 28.4/28.5 | 28.3/29.7 |
| Treatment 3 | 26.8/27.4 | 27.5/27.3 | 27.5/27.3 | 27.4/27.5 | 28.8/28.9 | 29.1/29.1 | 28.9/29.4 |

Supplementary Table 2: Water density (g/cm$^3$) measured during the heat stress experiment.

| | Water density (g/cm$^3$) measured on experimental day | | | | | | |
|---|---|---|---|---|---|---|---|
| Tank | 1 | 2 | 3 | 4 | 5 | 7 | 8 |
| Control 1 | 1.0217 | 1.0219 | 1.0216 | 1.0213 | 1.0216 | 1.0213 | 1.0217 |
| Control 2 | 1.0220 | 1.0212 | 1.0209 | 1.0207 | 1.0209 | 1.0209 | 1.0212 |
| Control 3 | 1.0223 | 1.0209 | 1.0208 | 1.0205 | 1.0206 | 1.0205 | 1.0207 |
| Treatment 1 | 1.0190 | 1.0219 | 1.0216 | 1.0219 | 1.0215 | 1.0215 | 1.0223 |
| Treatment 2 | 1.0217 | 1.0219 | 1.0221 | 1.0216 | 1.0218 | 1.0216 | 1.0226 |
| Treatment 3 | 1.0219 | 1.0217 | 1.0215 | 1.0218 | 1.0213 | 1.0215 | 1.0230 |

Supplementary Table 3: Water pH measured during the heat stress experiment.

| | Water pH measured on experimental day | | | | | | |
|---|---|---|---|---|---|---|---|
| Tank | 1 | 2 | 3 | 4 | 5 | 7 | 8 |
| Control 1 | - | 8.36 | 8.36 | 8.36 | 8.37 | 8.41 | 8.18 |
| Control 2 | - | 8.44 | 8.39 | 8.38 | 8.37 | 8.39 | 8.15 |
| Control 3 | - | 8.41 | 8.37 | 8.39 | 8.37 | 8.39 | 8.13 |
| Treatment 1 | - | 8.36 | 8.33 | 8.31 | 8.30 | 8.30 | 8.06 |
| Treatment 2 | - | 8.31 | 8.28 | 8.27 | 8.28 | 8.30 | 8.06 |
| Treatment 3 | - | 8.44 | 8.41 | 8.41 | 8.36 | 8.32 | 8.05 |
