## Supplementary Tables for "Transcriptional response of the calcification and stress response toolkits in an octocoral under heat and pH stress": Supplementary_Tables_4.pdf

Supplementary Table 4: Details of the RNASeq libraries sequenced for the heat stress and low pH experiments

| Sample code | Treatment | Pairs sequenced |
|---|---|---|
| C1.1.1 | Heat stress control | 20,330,948 |
| C1.2.1 | Heat stress control | 28,086,704 |
| C1.3.2 | Heat stress control | 21,401,348 |
| C2.1.2 | Heat stress control | 24,002,067 |
| C2.2.1 | Heat stress control | 19,455,882 |
| C2.3.1 | Heat stress control | 18,832,145 |
| C3.1.1 | Heat stress control | 24,054,859 |
| C3.2.1 | Heat stress control | 21,405,568 |
| C3.3.1 | Heat stress control | 21,433,801 |
| T1.1.2 | Heat stress treatment | 26,501,908 |
| T1.2.1 | Heat stress treatment | 23,504,198 |
| T1.3.2 | Heat stress treatment | 22,630,392 |
| T2.1.1 | Heat stress treatment | 33,928,544 |
| T2.1.2 | Heat stress treatment | 25,402,530 |
| T2.2.1 | Heat stress treatment | 25,632,897 |
| T2.3.1 | Heat stress treatment | 19,044,893 |
| T3.1.1 | Heat stress treatment | 22,612,456 |
| T3.2.1 | Heat stress treatment | 23,801,199 |
| T3.3.1 | Heat stress treatment | 18,448,797 |
| C1 | pH control | 20,491,967 |
| C2 | pH control | 15,000,640 |
| C3 | pH control | 23,266,075 |
| C4 | pH control | 20,168,689 |
| C6 | pH control | 15,053,888 |
| T1 | pH treatment | 18,244,934 |
| T2 | pH treatment | 15,045,703 |
| T4 | pH treatment | 19,359,013 |
| T5 | pH treatment | 20,546,854 |
| T6 | pH treatment | 25,320,180 |
