## Supplementary Tables for "Transcriptional response of the calcification and stress response toolkits in an octocoral under heat and pH stress": Supplementary_Tables_6-19.pdf

Supplementary Tables 6-12: GO-term enrichment analysis of the set of significantly overexpressed genes during heat stress. TopGO results showing significantly enriched GO-terms clustered by semantic similarity. GO.IC refers to the GO-term *Information Content* as determined using GOSemSim; larger values are assigned to more specific GO-terms.

S. Table 6: “Darkblue” Cluster.

S. Table 7: “Darkgreen” Cluster.

S. Table 8: “Mediumorchid3” Cluster.

S. Table 9: “Orange3” Cluster.

S. Table 10: “Firebrick4” Cluster.

S. Table 11: “Gray40” Cluster.

S. Table 12: “Violetred2” Cluster.

NOTE: cluster names refer to the Rcolor used to plot them in the dendrogram.

### Temp. OverDEGs Cluster darkblue

| | GO.Terms.TERM | Annotated | Significant | Expected | classicFisher | GO.IC |
|---|---|---|---|---|---|---|
| *GO:0010656* | negative regulation of muscle cell apoptotic process | 22 | 2 | 0.13 | 0.00741 | 8.991838 |
| *GO:0010657* | muscle cell apoptotic process | 37 | 3 | 0.22 | 0.00135 | 8.463771 |
| *GO:0010658* | striated muscle cell apoptotic process | 25 | 3 | 0.15 | 0.00042 | 8.869236 |
| *GO:0010659* | cardiac muscle cell apoptotic process | 22 | 3 | 0.13 | 0.00029 | 8.991838 |
| *GO:0010660* | regulation of muscle cell apoptotic process | 35 | 3 | 0.21 | 0.00115 | 8.543813 |
| *GO:0010662* | regulation of striated muscle cell apoptotic process | 23 | 3 | 0.14 | 0.00033 | 8.991838 |
| *GO:0010664* | negative regulation of striated muscle cell apoptotic process | 16 | 2 | 0.09 | 0.00394 | 9.419282 |
| *GO:0010665* | regulation of cardiac muscle cell apoptotic process | 20 | 3 | 0.12 | 0.00021 | 9.131600 |
| *GO:0010667* | negative regulation of cardiac muscle cell apoptotic process | 13 | 2 | 0.08 | 0.00259 | 9.642426 |
| *GO:0012501* | programmed cell death | 1359 | 16 | 8.03 | 0.00663 | 4.010617 |
| *GO:0042981* | regulation of apoptotic process | 917 | 12 | 5.42 | 0.00833 | 4.605473 |
| *GO:0043069* | negative regulation of programmed cell death | 492 | 8 | 2.91 | 0.00902 | 5.366918 |
| *GO:1904037* | positive regulation of epithelial cell apoptotic process | 12 | 2 | 0.07 | 0.00220 | 9.729437 |

### Temp. OverDEGs Cluster darkgreen

| | GO.Terms.TERM | Annotated | Significant | Expected | classicFisher | GO.IC |
|---|---|---|---|---|---|---|
| *GO:0001562* | response to protozoan | 16 | 2 | 0.09 | 0.00394 | 9.036290 |
| *GO:0002250* | adaptive immune response | 141 | 4 | 0.83 | 0.00988 | 6.697987 |
| *GO:0002819* | regulation of adaptive immune response | 74 | 3 | 0.44 | 0.00964 | 7.408834 |
| *GO:0006950* | response to stress | 2907 | 29 | 17.17 | 0.00318 | 2.784912 |
| *GO:0006953* | acute−phase response | 21 | 2 | 0.12 | 0.00676 | 8.908457 |
| *GO:0006970* | response to osmotic stress | 86 | 5 | 0.51 | 0.00016 | 6.420222 |
| *GO:0014823* | response to activity | 32 | 3 | 0.19 | 0.00088 | 8.320670 |
| *GO:0042832* | defense response to protozoan | 16 | 2 | 0.09 | 0.00394 | 9.036290 |
| *GO:0051851* | modulation by host of symbiont process | 24 | 2 | 0.14 | 0.00878 | 8.693345 |

### Temp. OverDEGs Cluster mediumorchid3

| | GO.Terms.TERM | Annotated | Significant | Expected | classicFisher | GO.IC |
|---|---|---|---|---|---|---|
| *GO:0009410* | response to xenobiotic stimulus | 152 | 5 | 0.90 | 0.00212 | 6.481886 |
| *GO:0010033* | response to organic substance | 2190 | 27 | 12.94 | 0.00019 | 3.392451 |
| *GO:0033993* | response to lipid | 524 | 9 | 3.10 | 0.00404 | 5.047306 |
| *GO:0034620* | cellular response to unfolded protein | 120 | 4 | 0.71 | 0.00565 | 6.729170 |
| *GO:0035967* | cellular response to topologically incorrect protein | 139 | 4 | 0.82 | 0.00941 | 6.532621 |
| *GO:0042221* | response to chemical | 3139 | 34 | 18.54 | 0.00030 | 2.876234 |
| *GO:0043200* | response to amino acid | 73 | 3 | 0.43 | 0.00929 | 7.435985 |
| *GO:0070555* | response to interleukin−1 | 70 | 3 | 0.41 | 0.00828 | 7.584038 |
| *GO:0070887* | cellular response to chemical stimulus | 2171 | 25 | 12.82 | 0.00092 | 3.538563 |
| *GO:0071310* | cellular response to organic substance | 1669 | 19 | 9.86 | 0.00449 | 3.895158 |

### Temp. OverDEGs Cluster orange3

| | GO.Terms.TERM | Annotated | Significant | Expected | classicFisher | GO.IC |
|---|---|---|---|---|---|---|
| *GO:0000096* | sulfur amino acid metabolic process | 54 | 3 | 0.32 | 0.00402 | 6.702382 |
| *GO:0006082* | organic acid metabolic process | 1026 | 14 | 6.06 | 0.00310 | 3.741388 |
| *GO:0006413* | translational initiation | 135 | 4 | 0.80 | 0.00851 | 6.103885 |
| *GO:0006555* | methionine metabolic process | 24 | 2 | 0.14 | 0.00878 | 7.454504 |
| *GO:0006739* | NADP metabolic process | 49 | 3 | 0.29 | 0.00305 | 7.291051 |
| *GO:0009086* | methionine biosynthetic process | 23 | 2 | 0.14 | 0.00808 | 7.573456 |
| *GO:0009404* | toxin metabolic process | 20 | 2 | 0.12 | 0.00614 | 8.000198 |
| *GO:0017144* | drug metabolic process | 503 | 10 | 2.97 | 0.00084 | 4.917992 |
| *GO:0019748* | secondary metabolic process | 73 | 4 | 0.43 | 0.00093 | 6.824028 |
| *GO:0044272* | sulfur compound biosynthetic process | 134 | 5 | 0.79 | 0.00122 | 6.143396 |
| *GO:0046500* | S−adenosylmethionine metabolic process | 18 | 3 | 0.11 | 0.00015 | 8.726135 |
| *GO:0055114* | oxidation−reduction process | 1272 | 19 | 7.51 | 0.00018 | 3.978598 |
| *GO:0070995* | NADPH oxidation | 7 | 2 | 0.04 | 0.00071 | 9.729437 |

### Temp. OverDEGs Cluster firebrick4

| | GO.Terms.TERM | Annotated | Significant | Expected | classicFisher | GO.IC |
|---|---|---|---|---|---|---|
| *GO:0002792* | negative regulation of peptide secretion | 73 | 3 | 0.43 | 0.00929 | 7.522162 |
| *GO:0046676* | negative regulation of insulin secretion | 22 | 2 | 0.13 | 0.00741 | 8.726135 |
| *GO:0050709* | negative regulation of protein secretion | 70 | 3 | 0.41 | 0.00828 | 7.573456 |

#### Temp. OverDEGs Cluster gray40

| | GO.Terms.TERM | Annotated | Significant | Expected | classicFisher | GO.IC |
|---|---|---|---|---|---|---|
| *GO:0008544* | epidermis development | 209 | 5 | 1.23 | 0.00813 | 6.517861 |
| *GO:0009913* | epidermal cell differentiation | 124 | 4 | 0.73 | 0.00634 | 7.045928 |
| *GO:0045604* | regulation of epidermal cell differentiation | 30 | 3 | 0.18 | 0.00073 | 8.413760 |
| *GO:0045682* | regulation of epidermis development | 41 | 3 | 0.24 | 0.00183 | 8.277185 |

#### Temp. OverDEGs Cluster violetred2

| | GO.Terms.TERM | Annotated | Significant | Expected | classicFisher | GO.IC |
|---|---|---|---|---|---|---|
| *GO:0007249* | I−kappaB kinase/NF−kappaB signaling | 172 | 6 | 1.02 | 0.00057 | 6.289602 |
| *GO:0019725* | cellular homeostasis | 697 | 12 | 4.12 | 0.00091 | 4.716381 |
| *GO:0043122* | regulation of I−kappaB kinase/NF−kappaB signaling | 149 | 5 | 0.88 | 0.00194 | 6.551383 |
| *GO:0051085* | chaperone cofactor−dependent protein refolding | 25 | 2 | 0.15 | 0.00951 | 8.413760 |
| *GO:1903332* | regulation of protein folding | 9 | 2 | 0.05 | 0.00121 | 9.294119 |
| *GO:1903334* | positive regulation of protein folding | 6 | 2 | 0.04 | 0.00051 | 10.335573 |

Supplementary Tables 13-19: GO-term enrichment analysis of the set of significantly underexpressed genes during heat stress. TopGO results showing significantly enriched GO-terms clustered by semantic similarity. GO.IC refers to the GO-term *Information Content* as determined using GOSemSim; larger values are assigned to more specific GO-terms.

S. Table 13: “Darkblue” Cluster.

S. Table 14: “Darkgreen” Cluster.

S. Table 15: “Mediumorchid3” Cluster.

S. Table 16: “Orange3” Cluster.

S. Table 17: “Firebrick4” Cluster.

S. Table 18: “Gray40” Cluster.

S. Table 19: “Violetred2” Cluster.

NOTE: cluster names refer to the Rcolor used to plot them in the dendrogram.

#### Temp. UnderDEGs Cluster darkblue

| | GO.Terms.TERM | Annotated | Significant | Expected | classicFisher | GO.IC |
|---|---|---|---|---|---|---|
| *GO:0008631* | intrinsic apoptotic signaling pathway in response to oxidative stress | 31 | 3 | 0.35 | 0.00507 | 8.600972 |
| *GO:0036473* | cell death in response to oxidative stress | 53 | 5 | 0.60 | 0.00033 | 7.893226 |
| *GO:0036475* | neuron death in response to oxidative stress | 21 | 3 | 0.24 | 0.00163 | 8.795128 |
| *GO:0062197* | cellular response to chemical stress | 255 | 9 | 2.88 | 0.00257 | 5.983147 |
| *GO:1900038* | negative regulation of cellular response to hypoxia | 11 | 2 | 0.12 | 0.00653 | 9.488275 |
| *GO:1900407* | regulation of cellular response to oxidative stress | 56 | 4 | 0.63 | 0.00369 | 7.783527 |
| *GO:1900408* | negative regulation of cellular response to oxidative stress | 31 | 4 | 0.35 | 0.00039 | 8.389663 |
| *GO:1901031* | regulation of response to reactive oxygen species | 32 | 3 | 0.36 | 0.00555 | 8.438453 |
| *GO:1902883* | negative regulation of response to oxidative stress | 34 | 4 | 0.38 | 0.00057 | 8.298691 |
| *GO:1903201* | regulation of oxidative stress−induced cell death | 45 | 4 | 0.51 | 0.00165 | 8.066889 |
| *GO:1903202* | negative regulation of oxidative stress−induced cell death | 30 | 4 | 0.34 | 0.00035 | 8.438453 |
| *GO:1903203* | regulation of oxidative stress−induced neuron death | 20 | 3 | 0.23 | 0.00141 | 8.831496 |
| *GO:1903204* | negative regulation of oxidative stress−induced neuron death | 14 | 3 | 0.16 | 0.00047 | 9.236961 |

#### Temp. UnderDEGs Cluster darkgreen

| | GO.Terms.TERM | Annotated | Significant | Expected | classicFisher | GO.IC |
|---|---|---|---|---|---|---|
| *GO:0002062* | chondrocyte differentiation | 99 | 5 | 1.12 | 0.00535 | 7.482942 |
| *GO:0003414* | chondrocyte morphogenesis involved in endochondral bone morphogenesis | 39 | 3 | 0.44 | 0.00964 | 8.463771 |
| *GO:0003429* | growth plate cartilage chondrocyte morphogenesis | 39 | 3 | 0.44 | 0.00964 | 8.463771 |
| *GO:0003433* | chondrocyte development involved in endochondral bone morphogenesis | 39 | 3 | 0.44 | 0.00964 | 8.463771 |
| *GO:0035051* | cardiocyte differentiation | 100 | 5 | 1.13 | 0.00558 | 7.015345 |
| *GO:0048738* | cardiac muscle tissue development | 142 | 6 | 1.60 | 0.00560 | 6.593943 |
| *GO:0051216* | cartilage development | 157 | 6 | 1.77 | 0.00899 | 6.814126 |
| *GO:0060351* | cartilage development involved in endochondral bone morphogenesis | 62 | 4 | 0.70 | 0.00531 | 8.000198 |
| *GO:0090171* | chondrocyte morphogenesis | 39 | 3 | 0.44 | 0.00964 | 8.463771 |

#### Temp. UnderDEGs Cluster mediumorchid3

| | GO.Terms.TERM | Annotated | Significant | Expected | classicFisher | GO.IC |
|---|---|---|---|---|---|---|
| *GO:0019367* | fatty acid elongation, saturated fatty acid | 10 | 2 | 0.11 | 0.00539 | 9.354744 |
| *GO:0019368* | fatty acid elongation, unsaturated fatty acid | 10 | 2 | 0.11 | 0.00539 | 8.661597 |
| *GO:0034625* | fatty acid elongation, monounsaturated fatty acid | 10 | 2 | 0.11 | 0.00539 | 9.354744 |
| *GO:0034626* | fatty acid elongation, polyunsaturated fatty acid | 10 | 2 | 0.11 | 0.00539 | 9.354744 |

#### Temp. UnderDEGs Cluster orange3

| | GO.Terms.TERM | Annotated | Significant | Expected | classicFisher | GO.IC |
|---|---|---|---|---|---|---|
| *GO:0006111* | regulation of gluconeogenesis | 31 | 3 | 0.35 | 0.00507 | 8.366132 |
| *GO:0010906* | regulation of glucose metabolic process | 71 | 4 | 0.80 | 0.00855 | 7.426852 |
| *GO:0019748* | secondary metabolic process | 73 | 4 | 0.82 | 0.00941 | 6.824028 |
| *GO:0032776* | DNA methylation on cytosine | 10 | 2 | 0.11 | 0.00539 | 8.908457 |
| *GO:0033866* | nucleoside bisphosphate biosynthetic process | 51 | 4 | 0.58 | 0.00262 | 7.283145 |
| *GO:0034030* | ribonucleoside bisphosphate biosynthetic process | 51 | 4 | 0.58 | 0.00262 | 7.283145 |
| *GO:0034033* | purine nucleoside bisphosphate biosynthetic process | 51 | 4 | 0.58 | 0.00262 | 7.283145 |
| *GO:0035384* | thioester biosynthetic process | 37 | 3 | 0.42 | 0.00834 | 7.522162 |
| *GO:0044283* | small molecule biosynthetic process | 573 | 14 | 6.47 | 0.00590 | 4.416231 |
| *GO:0046128* | purine ribonucleoside metabolic process | 73 | 4 | 0.82 | 0.00941 | 6.997434 |
| *GO:0071616* | acyl−CoA biosynthetic process | 37 | 3 | 0.42 | 0.00834 | 7.522162 |
| *GO:0090116* | C−5 methylation of cytosine | 9 | 2 | 0.10 | 0.00434 | 9.131600 |

### Temp. UnderDEGs Cluster firebrick4

| | GO.Terms.TERM | Annotated | Significant | Expected | classicFisher | GO.IC |
|---|---|---|---|---|---|---|
| *GO:0002576* | platelet degranulation | 33 | 3 | 0.37 | 0.00605 | 8.277185 |
| *GO:0006955* | immune response | 778 | 17 | 8.78 | 0.00759 | 4.665692 |
| *GO:0008228* | opsonization | 9 | 2 | 0.10 | 0.00434 | 9.824747 |
| *GO:0010917* | negative regulation of mitochondrial membrane potential | 11 | 2 | 0.12 | 0.00653 | 9.642426 |
| *GO:0015727* | lactate transport | 7 | 2 | 0.08 | 0.00257 | 9.930108 |
| *GO:0031000* | response to caffeine | 13 | 2 | 0.15 | 0.00913 | 8.760037 |
| *GO:0032355* | response to estradiol | 65 | 4 | 0.73 | 0.00628 | 7.770624 |
| *GO:0032780* | negative regulation of ATPase activity | 12 | 2 | 0.14 | 0.00778 | 9.488275 |
| *GO:0033189* | response to vitamin A | 6 | 2 | 0.07 | 0.00185 | 9.930108 |
| *GO:0035873* | lactate transmembrane transport | 7 | 2 | 0.08 | 0.00257 | 10.047891 |
| *GO:0036270* | response to diuretic | 13 | 2 | 0.15 | 0.00913 | 8.760037 |
| *GO:0045055* | regulated exocytosis | 329 | 10 | 3.71 | 0.00442 | 5.626043 |
| *GO:0045837* | negative regulation of membrane potential | 11 | 2 | 0.12 | 0.00653 | 9.562383 |
| *GO:0051029* | rRNA transport | 7 | 2 | 0.08 | 0.00257 | 9.729437 |
| *GO:0051573* | negative regulation of histone H3−K9 methylation | 5 | 2 | 0.06 | 0.00124 | 10.517895 |

#### Temp. UnderDEGs Cluster gray40

| | GO.Terms.TERM | Annotated | Significant | Expected | classicFisher | GO.IC |
|---|---|---|---|---|---|---|
| *GO:0010737* | protein kinase A signaling | 64 | 4 | 0.72 | 0.00595 | 7.661424 |
| *GO:0010939* | regulation of necrotic cell death | 29 | 3 | 0.33 | 0.00419 | 8.101981 |
| *GO:0010940* | positive regulation of necrotic cell death | 7 | 2 | 0.08 | 0.00257 | 9.562383 |
| *GO:0042268* | regulation of cytolysis | 9 | 2 | 0.10 | 0.00434 | 9.729437 |
| *GO:0045919* | positive regulation of cytolysis | 9 | 2 | 0.10 | 0.00434 | 9.729437 |
| *GO:1900076* | regulation of cellular response to insulin stimulus | 51 | 4 | 0.58 | 0.00262 | 7.952945 |
| *GO:2000310* | regulation of NMDA receptor activity | 8 | 2 | 0.09 | 0.00340 | 9.562383 |

#### Temp. UnderDEGs Cluster violetred2

| | GO.Terms.TERM | Annotated | Significant | Expected | classicFisher | GO.IC |
|---|---|---|---|---|---|---|
| *GO:0036295* | cellular response to increased oxygen levels | 10 | 2 | 0.11 | 0.00539 | 9.562383 |
| *GO:0071453* | cellular response to oxygen levels | 129 | 6 | 1.46 | 0.00351 | 6.684915 |
| *GO:0071455* | cellular response to hyperoxia | 6 | 2 | 0.07 | 0.00185 | 9.930108 |
| *GO:0090649* | response to oxygen−glucose deprivation | 11 | 2 | 0.12 | 0.00653 | 9.419282 |
| *GO:0090650* | cellular response to oxygen−glucose deprivation | 9 | 2 | 0.10 | 0.00434 | 9.562383 |
