## Supplementary Tables for "Transcriptional response of the calcification and stress response toolkits in an octocoral under heat and pH stress": Supplementary_Tables_22-30.pdf

Supplementary Tables 22-30: GO-term enrichment analysis of the set of significantly underexpressed genes during low pH stress. TopGO results showing significantly enriched GO-terms clustered by semantic similarity. GO.IC refers to the GO-term *Information Content* as determined using GOSemSim; larger values are assigned to more specific GO-terms.

S. Table 22: “Darkblue” Cluster.

S. Table 23: “Darkgreen” Cluster.

S. Table 24: “Mediumorchid3” Cluster.

S. Table 25: “Orange3” Cluster.

S. Table 26: “Firebrick4” Cluster.

S. Table 27: “Gray40” Cluster.

S. Table 28: “Violetred2” Cluster.

S. Table 29: “Red” Cluster.

S. Table 30: “Blue” Cluster.

NOTE: cluster names refer to the Rcolor used to plot them in the dendrogram.

### pH UnderDEGs Cluster darkblue

| | GO.Terms.TERM | Annotated | Significant | Expected | classicFisher | GO.IC |
|---|---|---|---|---|---|---|
| *GO:1902959* | regulation of aspartic−type endopeptidase activity involved in amyloid precursor protein catabolic process | 7 | 1 | 0.04 | 0.0374 | 9.824747 |
| *GO:1902962* | regulation of metalloendopeptidase activity involved in amyloid precursor protein catabolic process | 5 | 1 | 0.03 | 0.0268 | 10.335573 |
| *GO:1902963* | negative regulation of metalloendopeptidase activity involved in amyloid precursor protein catabolic process | 5 | 1 | 0.03 | 0.0268 | 10.335573 |
| *GO:1904683* | regulation of metalloendopeptidase activity | 7 | 1 | 0.04 | 0.0374 | 10.047891 |
| *GO:1904684* | negative regulation of metalloendopeptidase activity | 7 | 1 | 0.04 | 0.0374 | 10.047891 |
| *GO:1905245* | regulation of aspartic−type peptidase activity | 9 | 1 | 0.05 | 0.0478 | 9.642426 |

### pH UnderDEGs Cluster darkgreen

| | GO.Terms.TERM | Annotated | Significant | Expected | classicFisher | GO.IC |
|---|---|---|---|---|---|---|
| *GO:0035418* | protein localization to synapse | 65 | 2 | 0.35 | 0.0487 | 7.584038 |
| *GO:0062237* | protein localization to postsynapse | 49 | 2 | 0.27 | 0.0291 | 7.937678 |
| *GO:0097120* | receptor localization to synapse | 54 | 2 | 0.29 | 0.0348 | 7.836873 |
| *GO:0099633* | protein localization to postsynaptic specialization membrane | 18 | 2 | 0.10 | 0.0042 | 9.036290 |
| *GO:0099645* | neurotransmitter receptor localization to postsynaptic specialization membrane | 18 | 2 | 0.10 | 0.0042 | 9.036290 |
| *GO:1903539* | protein localization to postsynaptic membrane | 41 | 2 | 0.22 | 0.0209 | 8.235512 |

### pH UnderDEGs Cluster mediumorchid3

| | GO.Terms.TERM | Annotated | Significant | Expected | classicFisher | GO.IC |
|---|---|---|---|---|---|---|
| *GO:0030500* | regulation of bone mineralization | 61 | 2 | 0.33 | 0.0434 | 7.952945 |
| *GO:0035886* | vascular smooth muscle cell differentiation | 9 | 1 | 0.05 | 0.0478 | 10.047891 |
| *GO:0048844* | artery morphogenesis | 61 | 2 | 0.33 | 0.0434 | 7.732883 |
| *GO:0060736* | prostate gland growth | 7 | 1 | 0.04 | 0.0374 | 10.047891 |
| *GO:0060742* | epithelial cell differentiation involved in prostate gland development | 5 | 1 | 0.03 | 0.0268 | 10.741038 |
| *GO:0070167* | regulation of biomineral tissue development | 65 | 2 | 0.35 | 0.0487 | 7.907825 |
| *GO:0072103* | glomerulus vasculature morphogenesis | 9 | 1 | 0.05 | 0.0478 | 9.824747 |
| *GO:0072104* | glomerular capillary formation | 9 | 1 | 0.05 | 0.0478 | 9.824747 |
| *GO:1904177* | regulation of adipose tissue development | 7 | 1 | 0.04 | 0.0374 | 10.181422 |
| *GO:1904179* | positive regulation of adipose tissue development | 5 | 1 | 0.03 | 0.0268 | 10.517895 |

### pH UnderDEGs Cluster orange3

| | GO.Terms.TERM | Annotated | Significant | Expected | classicFisher | GO.IC |
|---|---|---|---|---|---|---|
| *GO:0001523* | retinoid metabolic process | 51 | 2 | 0.28 | 0.0313 | 7.473372 |
| *GO:0006530* | asparagine catabolic process | 8 | 1 | 0.04 | 0.0426 | 9.729437 |
| *GO:0006721* | terpenoid metabolic process | 59 | 2 | 0.32 | 0.0409 | 6.844129 |
| *GO:0016101* | diterpenoid metabolic process | 54 | 2 | 0.29 | 0.0348 | 7.382400 |
| *GO:0016114* | terpenoid biosynthetic process | 8 | 1 | 0.04 | 0.0426 | 8.000198 |
| *GO:0033345* | asparagine catabolic process via L−aspartate | 8 | 1 | 0.04 | 0.0426 | 10.047891 |
| *GO:0042573* | retinoic acid metabolic process | 5 | 1 | 0.03 | 0.0268 | 9.562383 |

#### pH UnderDEGs Cluster firebrick4

| | GO.Terms.TERM | Annotated | Significant | Expected | classicFisher | GO.IC |
|---|---|---|---|---|---|---|
| *GO:0044409* | entry into host | 71 | 3 | 0.39 | 0.0068 | 7.584038 |
| *GO:0046718* | viral entry into host cell | 64 | 3 | 0.35 | 0.0051 | 7.684681 |
| *GO:0051701* | interaction with host | 118 | 3 | 0.64 | 0.0265 | 6.923326 |
| *GO:0052126* | movement in host environment | 83 | 3 | 0.45 | 0.0105 | 7.391134 |

#### pH UnderDEGs Cluster gray40

| | GO.Terms.TERM | Annotated | Significant | Expected | classicFisher | GO.IC |
|---|---|---|---|---|---|---|
| *GO:0007178* | transmembrane receptor protein serine/threonine kinase signaling pathway | 234 | 5 | 1.27 | 0.0091 | 6.133371 |
| *GO:0010469* | regulation of signaling receptor activity | 53 | 2 | 0.29 | 0.0336 | 7.809844 |
| *GO:0030509* | BMP signaling pathway | 111 | 3 | 0.60 | 0.0226 | 7.222058 |
| *GO:0034138* | toll−like receptor 3 signaling pathway | 6 | 1 | 0.03 | 0.0321 | 10.181422 |
| *GO:0071772* | response to BMP | 116 | 3 | 0.63 | 0.0254 | 7.136900 |
| *GO:0071773* | cellular response to BMP stimulus | 116 | 3 | 0.63 | 0.0254 | 7.136900 |
| *GO:0090092* | regulation of transmembrane receptor protein serine/threonine kinase signaling pathway | 179 | 4 | 0.97 | 0.0165 | 6.601879 |
| *GO:0099601* | regulation of neurotransmitter receptor activity | 26 | 2 | 0.14 | 0.0087 | 8.661597 |

### pH UnderDEGs Cluster violetred2

| | GO.Terms.TERM | Annotated | Significant | Expected | classicFisher | GO.IC |
|---|---|---|---|---|---|---|
| GO:0001976 | nervous system process involved in regulation of systemic arterial blood pressure | 7 | 1 | 0.04 | 0.0374 | 9.824747 |
| GO:0006616 | SRP−dependent cotranslational protein targeting to membrane, translocation | 6 | 1 | 0.03 | 0.0321 | 9.562383 |
| GO:0007157 | heterophilic cell−cell adhesion via plasma membrane cell adhesion molecules | 104 | 3 | 0.56 | 0.0191 | 7.864653 |
| GO:0007565 | female pregnancy | 59 | 3 | 0.32 | 0.0041 | 7.435985 |
| GO:0010897 | negative regulation of triglyceride catabolic process | 7 | 1 | 0.04 | 0.0374 | 10.047891 |
| GO:0010940 | positive regulation of necrotic cell death | 7 | 1 | 0.04 | 0.0374 | 9.562383 |
| GO:0015827 | tryptophan transport | 7 | 1 | 0.04 | 0.0374 | 9.930108 |
| GO:0031204 | posttranslational protein targeting to membrane, translocation | 7 | 1 | 0.04 | 0.0374 | 9.419282 |
| GO:0031342 | negative regulation of cell killing | 6 | 1 | 0.03 | 0.0321 | 9.930108 |
| GO:0031644 | regulation of nervous system process | 84 | 3 | 0.46 | 0.0108 | 7.283145 |
| GO:0032460 | negative regulation of protein oligomerization | 9 | 1 | 0.05 | 0.0478 | 9.824747 |
| GO:0032733 | positive regulation of interleukin−10 production | 8 | 1 | 0.04 | 0.0426 | 9.824747 |
| GO:0032800 | receptor biosynthetic process | 9 | 1 | 0.05 | 0.0478 | 9.642426 |
| GO:0044706 | multi−multicellular organism process | 79 | 3 | 0.43 | 0.0091 | 6.794614 |
| GO:0045185 | maintenance of protein location | 124 | 3 | 0.67 | 0.0301 | 6.854333 |
| GO:0045989 | positive regulation of striated muscle contraction | 8 | 1 | 0.04 | 0.0426 | 9.236961 |
| GO:0050715 | positive regulation of cytokine secretion | 51 | 2 | 0.28 | 0.0313 | 7.562984 |
| GO:0050905 | neuromuscular process | 63 | 2 | 0.34 | 0.0460 | 7.532213 |
| GO:0060135 | maternal process involved in female pregnancy | 19 | 2 | 0.10 | 0.0047 | 9.036290 |
| GO:0061502 | early endosome to recycling endosome transport | 8 | 1 | 0.04 | 0.0426 | 9.930108 |
| GO:0072643 | interferon−gamma secretion | 7 | 1 | 0.04 | 0.0374 | 10.047891 |
| GO:0099072 | regulation of postsynaptic membrane neurotransmitter receptor levels | 56 | 2 | 0.30 | 0.0372 | 7.809844 |
| GO:0110149 | regulation of biomineralization | 65 | 2 | 0.35 | 0.0487 | 7.907825 |
| GO:0150078 | positive regulation of neuroinflammatory response | 7 | 1 | 0.04 | 0.0374 | 10.181422 |
| GO:1900426 | positive regulation of defense response to bacterium | 9 | 1 | 0.05 | 0.0478 | 9.182893 |
| GO:1901739 | regulation of myoblast fusion | 9 | 1 | 0.05 | 0.0478 | 9.182893 |
| GO:1901741 | positive regulation of myoblast fusion | 9 | 1 | 0.05 | 0.0478 | 9.236961 |
| GO:1902713 | regulation of interferon−gamma secretion | 6 | 1 | 0.03 | 0.0321 | 10.335573 |
| GO:1902769 | regulation of choline O−acetyltransferase activity | 5 | 1 | 0.03 | 0.0268 | 10.181422 |
| GO:1902954 | regulation of early endosome to recycling endosome transport | 6 | 1 | 0.03 | 0.0321 | 10.181422 |
| GO:1903980 | positive regulation of microglial cell activation | 6 | 1 | 0.03 | 0.0321 | 10.335573 |
| GO:2000679 | positive regulation of transcription regulatory region DNA binding | 9 | 1 | 0.05 | 0.0478 | 10.181422 |
| GO:2000778 | positive regulation of interleukin−6 secretion | 8 | 1 | 0.04 | 0.0426 | 9.729437 |
| GO:2001135 | regulation of endocytic recycling | 9 | 1 | 0.05 | 0.0478 | 9.236961 |

### pH UnderDEGs Cluster red

| | GO.Terms.TERM | Annotated | Significant | Expected | classicFisher | GO.IC |
|---|---|---|---|---|---|---|
| *GO:0032814* | regulation of natural killer cell activation | 5 | 1 | 0.03 | 0.0268 | 9.729437 |
| *GO:0032944* | regulation of mononuclear cell proliferation | 64 | 2 | 0.35 | 0.0473 | 7.638696 |
| *GO:0032945* | negative regulation of mononuclear cell proliferation | 29 | 2 | 0.16 | 0.0108 | 8.463771 |
| *GO:0042116* | macrophage activation | 38 | 2 | 0.21 | 0.0181 | 8.176089 |
| *GO:0042129* | regulation of T cell proliferation | 44 | 2 | 0.24 | 0.0238 | 8.016459 |
| *GO:0042130* | negative regulation of T cell proliferation | 20 | 2 | 0.11 | 0.0052 | 8.795128 |
| *GO:0043030* | regulation of macrophage activation | 30 | 2 | 0.16 | 0.0115 | 8.600972 |
| *GO:0043032* | positive regulation of macrophage activation | 10 | 2 | 0.05 | 0.0013 | 9.562383 |
| *GO:0046640* | regulation of alpha−beta T cell proliferation | 8 | 1 | 0.04 | 0.0426 | 10.047891 |
| *GO:0050670* | regulation of lymphocyte proliferation | 64 | 2 | 0.35 | 0.0473 | 7.638696 |
| *GO:0050672* | negative regulation of lymphocyte proliferation | 29 | 2 | 0.16 | 0.0108 | 8.463771 |
| *GO:0050868* | negative regulation of T cell activation | 42 | 2 | 0.23 | 0.0218 | 8.084281 |
| *GO:0051250* | negative regulation of lymphocyte activation | 62 | 2 | 0.34 | 0.0447 | 7.638696 |
| *GO:0070663* | regulation of leukocyte proliferation | 64 | 2 | 0.35 | 0.0473 | 7.627523 |
| *GO:0070664* | negative regulation of leukocyte proliferation | 29 | 2 | 0.16 | 0.0108 | 8.463771 |
| *GO:1903038* | negative regulation of leukocyte cell−cell adhesion | 44 | 2 | 0.24 | 0.0238 | 8.049795 |
| *GO:2000515* | negative regulation of CD4−positive, alpha−beta T cell activation | 7 | 1 | 0.04 | 0.0374 | 10.181422 |

Supplementary Tables 22-30: GO-term enrichment analysis of the set of significantly underexpressed genes during low pH stress. TopGO results showing significantly enriched GO-terms clustered by semantic similarity. GO.IC refers to the GO-term *Information Content* as determined using GOSemSim; larger values are assigned to more specific GO-terms.

S. Table 22: “Darkblue” Cluster.

S. Table 23: “Darkgreen” Cluster.

S. Table 24: “Mediumorchid3” Cluster.

S. Table 25: “Orange3” Cluster.

S. Table 26: “Firebrick4” Cluster.

S. Table 27: “Gray40” Cluster.

S. Table 28: “Violetred2” Cluster.

S. Table 29: “Red” Cluster.

S. Table 30: “Blue” Cluster.

NOTE: cluster names refer to the Rcolor used to plot them in the dendrogram.

### pH UnderDEGs Cluster darkblue

| | GO.Terms.TERM | Annotated | Significant | Expected | classicFisher | GO.IC |
|---|---|---|---|---|---|---|
| *GO:1902959* | regulation of aspartic−type endopeptidase activity involved in amyloid precursor protein catabolic process | 7 | 1 | 0.04 | 0.0374 | 9.824747 |
| *GO:1902962* | regulation of metalloendopeptidase activity involved in amyloid precursor protein catabolic process | 5 | 1 | 0.03 | 0.0268 | 10.335573 |
| *GO:1902963* | negative regulation of metalloendopeptidase activity involved in amyloid precursor protein catabolic process | 5 | 1 | 0.03 | 0.0268 | 10.335573 |
| *GO:1904683* | regulation of metalloendopeptidase activity | 7 | 1 | 0.04 | 0.0374 | 10.047891 |
| *GO:1904684* | negative regulation of metalloendopeptidase activity | 7 | 1 | 0.04 | 0.0374 | 10.047891 |
| *GO:1905245* | regulation of aspartic−type peptidase activity | 9 | 1 | 0.05 | 0.0478 | 9.642426 |

#### pH UnderDEGs Cluster darkgreen

| | GO.Terms.TERM | Annotated | Significant | Expected | classicFisher | GO.IC |
|---|---|---|---|---|---|---|
| *GO:0035418* | protein localization to synapse | 65 | 2 | 0.35 | 0.0487 | 7.584038 |
| *GO:0062237* | protein localization to postsynapse | 49 | 2 | 0.27 | 0.0291 | 7.937678 |
| *GO:0097120* | receptor localization to synapse | 54 | 2 | 0.29 | 0.0348 | 7.836873 |
| *GO:0099633* | protein localization to postsynaptic specialization membrane | 18 | 2 | 0.10 | 0.0042 | 9.036290 |
| *GO:0099645* | neurotransmitter receptor localization to postsynaptic specialization membrane | 18 | 2 | 0.10 | 0.0042 | 9.036290 |
| *GO:1903539* | protein localization to postsynaptic membrane | 41 | 2 | 0.22 | 0.0209 | 8.235512 |

### pH UnderDEGs Cluster mediumorchid3

| | GO.Terms.TERM | Annotated | Significant | Expected | classicFisher | GO.IC |
|---|---|---|---|---|---|---|
| *GO:0030500* | regulation of bone mineralization | 61 | 2 | 0.33 | 0.0434 | 7.952945 |
| *GO:0035886* | vascular smooth muscle cell differentiation | 9 | 1 | 0.05 | 0.0478 | 10.047891 |
| *GO:0048844* | artery morphogenesis | 61 | 2 | 0.33 | 0.0434 | 7.732883 |
| *GO:0060736* | prostate gland growth | 7 | 1 | 0.04 | 0.0374 | 10.047891 |
| *GO:0060742* | epithelial cell differentiation involved in prostate gland development | 5 | 1 | 0.03 | 0.0268 | 10.741038 |
| *GO:0070167* | regulation of biomineral tissue development | 65 | 2 | 0.35 | 0.0487 | 7.907825 |
| *GO:0072103* | glomerulus vasculature morphogenesis | 9 | 1 | 0.05 | 0.0478 | 9.824747 |
| *GO:0072104* | glomerular capillary formation | 9 | 1 | 0.05 | 0.0478 | 9.824747 |
| *GO:1904177* | regulation of adipose tissue development | 7 | 1 | 0.04 | 0.0374 | 10.181422 |
| *GO:1904179* | positive regulation of adipose tissue development | 5 | 1 | 0.03 | 0.0268 | 10.517895 |

### pH UnderDEGs Cluster orange3

| | GO.Terms.TERM | Annotated | Significant | Expected | classicFisher | GO.IC |
|---|---|---|---|---|---|---|
| *GO:0001523* | retinoid metabolic process | 51 | 2 | 0.28 | 0.0313 | 7.473372 |
| *GO:0006530* | asparagine catabolic process | 8 | 1 | 0.04 | 0.0426 | 9.729437 |
| *GO:0006721* | terpenoid metabolic process | 59 | 2 | 0.32 | 0.0409 | 6.844129 |
| *GO:0016101* | diterpenoid metabolic process | 54 | 2 | 0.29 | 0.0348 | 7.382400 |
| *GO:0016114* | terpenoid biosynthetic process | 8 | 1 | 0.04 | 0.0426 | 8.000198 |
| *GO:0033345* | asparagine catabolic process via L−aspartate | 8 | 1 | 0.04 | 0.0426 | 10.047891 |
| *GO:0042573* | retinoic acid metabolic process | 5 | 1 | 0.03 | 0.0268 | 9.562383 |

#### pH UnderDEGs Cluster firebrick4

| | GO.Terms.TERM | Annotated | Significant | Expected | classicFisher | GO.IC |
|---|---|---|---|---|---|---|
| *GO:0044409* | entry into host | 71 | 3 | 0.39 | 0.0068 | 7.584038 |
| *GO:0046718* | viral entry into host cell | 64 | 3 | 0.35 | 0.0051 | 7.684681 |
| *GO:0051701* | interaction with host | 118 | 3 | 0.64 | 0.0265 | 6.923326 |
| *GO:0052126* | movement in host environment | 83 | 3 | 0.45 | 0.0105 | 7.391134 |

#### pH UnderDEGs Cluster gray40

| | GO.Terms.TERM | Annotated | Significant | Expected | classicFisher | GO.IC |
|---|---|---|---|---|---|---|
| *GO:0007178* | transmembrane receptor protein serine/threonine kinase signaling pathway | 234 | 5 | 1.27 | 0.0091 | 6.133371 |
| *GO:0010469* | regulation of signaling receptor activity | 53 | 2 | 0.29 | 0.0336 | 7.809844 |
| *GO:0030509* | BMP signaling pathway | 111 | 3 | 0.60 | 0.0226 | 7.222058 |
| *GO:0034138* | toll−like receptor 3 signaling pathway | 6 | 1 | 0.03 | 0.0321 | 10.181422 |
| *GO:0071772* | response to BMP | 116 | 3 | 0.63 | 0.0254 | 7.136900 |
| *GO:0071773* | cellular response to BMP stimulus | 116 | 3 | 0.63 | 0.0254 | 7.136900 |
| *GO:0090092* | regulation of transmembrane receptor protein serine/threonine kinase signaling pathway | 179 | 4 | 0.97 | 0.0165 | 6.601879 |
| *GO:0099601* | regulation of neurotransmitter receptor activity | 26 | 2 | 0.14 | 0.0087 | 8.661597 |

### pH UnderDEGs Cluster violetred2

| | GO.Terms.TERM | Annotated | Significant | Expected | classicFisher | GO.IC |
|---|---|---|---|---|---|---|
| *GO:0001976* | nervous system process involved in regulation of systemic arterial blood pressure | 7 | 1 | 0.04 | 0.0374 | 9.824747 |
| *GO:0006616* | SRP−dependent cotranslational protein targeting to membrane, translocation | 6 | 1 | 0.03 | 0.0321 | 9.562383 |
| *GO:0007157* | heterophilic cell−cell adhesion via plasma membrane cell adhesion molecules | 104 | 3 | 0.56 | 0.0191 | 7.864653 |
| *GO:0007565* | female pregnancy | 59 | 3 | 0.32 | 0.0041 | 7.435985 |
| *GO:0010897* | negative regulation of triglyceride catabolic process | 7 | 1 | 0.04 | 0.0374 | 10.047891 |
| *GO:0010940* | positive regulation of necrotic cell death | 7 | 1 | 0.04 | 0.0374 | 9.562383 |
| *GO:0015827* | tryptophan transport | 7 | 1 | 0.04 | 0.0374 | 9.930108 |
| *GO:0031204* | posttranslational protein targeting to membrane, translocation | 7 | 1 | 0.04 | 0.0374 | 9.419282 |
| *GO:0031342* | negative regulation of cell killing | 6 | 1 | 0.03 | 0.0321 | 9.930108 |
| *GO:0031644* | regulation of nervous system process | 84 | 3 | 0.46 | 0.0108 | 7.283145 |
| *GO:0032460* | negative regulation of protein oligomerization | 9 | 1 | 0.05 | 0.0478 | 9.824747 |
| *GO:0032733* | positive regulation of interleukin−10 production | 8 | 1 | 0.04 | 0.0426 | 9.824747 |
| *GO:0032800* | receptor biosynthetic process | 9 | 1 | 0.05 | 0.0478 | 9.642426 |
| *GO:0044706* | multi−multicellular organism process | 79 | 3 | 0.43 | 0.0091 | 6.794614 |
| *GO:0045185* | maintenance of protein location | 124 | 3 | 0.67 | 0.0301 | 6.854333 |
| *GO:0045989* | positive regulation of striated muscle contraction | 8 | 1 | 0.04 | 0.0426 | 9.236961 |
| *GO:0050715* | positive regulation of cytokine secretion | 51 | 2 | 0.28 | 0.0313 | 7.562984 |
| *GO:0050905* | neuromuscular process | 63 | 2 | 0.34 | 0.0460 | 7.532213 |
| *GO:0060135* | maternal process involved in female pregnancy | 19 | 2 | 0.10 | 0.0047 | 9.036290 |
| *GO:0061502* | early endosome to recycling endosome transport | 8 | 1 | 0.04 | 0.0426 | 9.930108 |
| *GO:0072643* | interferon−gamma secretion | 7 | 1 | 0.04 | 0.0374 | 10.047891 |
| *GO:0099072* | regulation of postsynaptic membrane neurotransmitter receptor levels | 56 | 2 | 0.30 | 0.0372 | 7.809844 |
| *GO:0110149* | regulation of biomineralization | 65 | 2 | 0.35 | 0.0487 | 7.907825 |
| *GO:0150078* | positive regulation of neuroinflammatory response | 7 | 1 | 0.04 | 0.0374 | 10.181422 |
| *GO:1900426* | positive regulation of defense response to bacterium | 9 | 1 | 0.05 | 0.0478 | 9.182893 |
| *GO:1901739* | regulation of myoblast fusion | 9 | 1 | 0.05 | 0.0478 | 9.182893 |
| *GO:1901741* | positive regulation of myoblast fusion | 9 | 1 | 0.05 | 0.0478 | 9.236961 |
| *GO:1902713* | regulation of interferon−gamma secretion | 6 | 1 | 0.03 | 0.0321 | 10.335573 |
| *GO:1902769* | regulation of choline O−acetyltransferase activity | 5 | 1 | 0.03 | 0.0268 | 10.181422 |
| *GO:1902954* | regulation of early endosome to recycling endosome transport | 6 | 1 | 0.03 | 0.0321 | 10.181422 |
| *GO:1903980* | positive regulation of microglial cell activation | 6 | 1 | 0.03 | 0.0321 | 10.335573 |
| *GO:2000679* | positive regulation of transcription regulatory region DNA binding | 9 | 1 | 0.05 | 0.0478 | 10.181422 |
| *GO:2000778* | positive regulation of interleukin−6 secretion | 8 | 1 | 0.04 | 0.0426 | 9.729437 |
| *GO:2001135* | regulation of endocytic recycling | 9 | 1 | 0.05 | 0.0478 | 9.236961 |

### pH UnderDEGs Cluster red

| | GO.Terms.TERM | Annotated | Significant | Expected | classicFisher | GO.IC |
|---|---|---|---|---|---|---|
| *GO:0032814* | regulation of natural killer cell activation | 5 | 1 | 0.03 | 0.0268 | 9.729437 |
| *GO:0032944* | regulation of mononuclear cell proliferation | 64 | 2 | 0.35 | 0.0473 | 7.638696 |
| *GO:0032945* | negative regulation of mononuclear cell proliferation | 29 | 2 | 0.16 | 0.0108 | 8.463771 |
| *GO:0042116* | macrophage activation | 38 | 2 | 0.21 | 0.0181 | 8.176089 |
| *GO:0042129* | regulation of T cell proliferation | 44 | 2 | 0.24 | 0.0238 | 8.016459 |
| *GO:0042130* | negative regulation of T cell proliferation | 20 | 2 | 0.11 | 0.0052 | 8.795128 |
| *GO:0043030* | regulation of macrophage activation | 30 | 2 | 0.16 | 0.0115 | 8.600972 |
| *GO:0043032* | positive regulation of macrophage activation | 10 | 2 | 0.05 | 0.0013 | 9.562383 |
| *GO:0046640* | regulation of alpha−beta T cell proliferation | 8 | 1 | 0.04 | 0.0426 | 10.047891 |
| *GO:0050670* | regulation of lymphocyte proliferation | 64 | 2 | 0.35 | 0.0473 | 7.638696 |
| *GO:0050672* | negative regulation of lymphocyte proliferation | 29 | 2 | 0.16 | 0.0108 | 8.463771 |
| *GO:0050868* | negative regulation of T cell activation | 42 | 2 | 0.23 | 0.0218 | 8.084281 |
| *GO:0051250* | negative regulation of lymphocyte activation | 62 | 2 | 0.34 | 0.0447 | 7.638696 |
| *GO:0070663* | regulation of leukocyte proliferation | 64 | 2 | 0.35 | 0.0473 | 7.627523 |
| *GO:0070664* | negative regulation of leukocyte proliferation | 29 | 2 | 0.16 | 0.0108 | 8.463771 |
| *GO:1903038* | negative regulation of leukocyte cell−cell adhesion | 44 | 2 | 0.24 | 0.0238 | 8.049795 |
| *GO:2000515* | negative regulation of CD4−positive, alpha−beta T cell activation | 7 | 1 | 0.04 | 0.0374 | 10.181422 |
