## Supplementary Figures for "Transcriptional response of the calcification and stress response toolkits in an octocoral under heat and pH stress": Supplementary_Figures_1.pdf

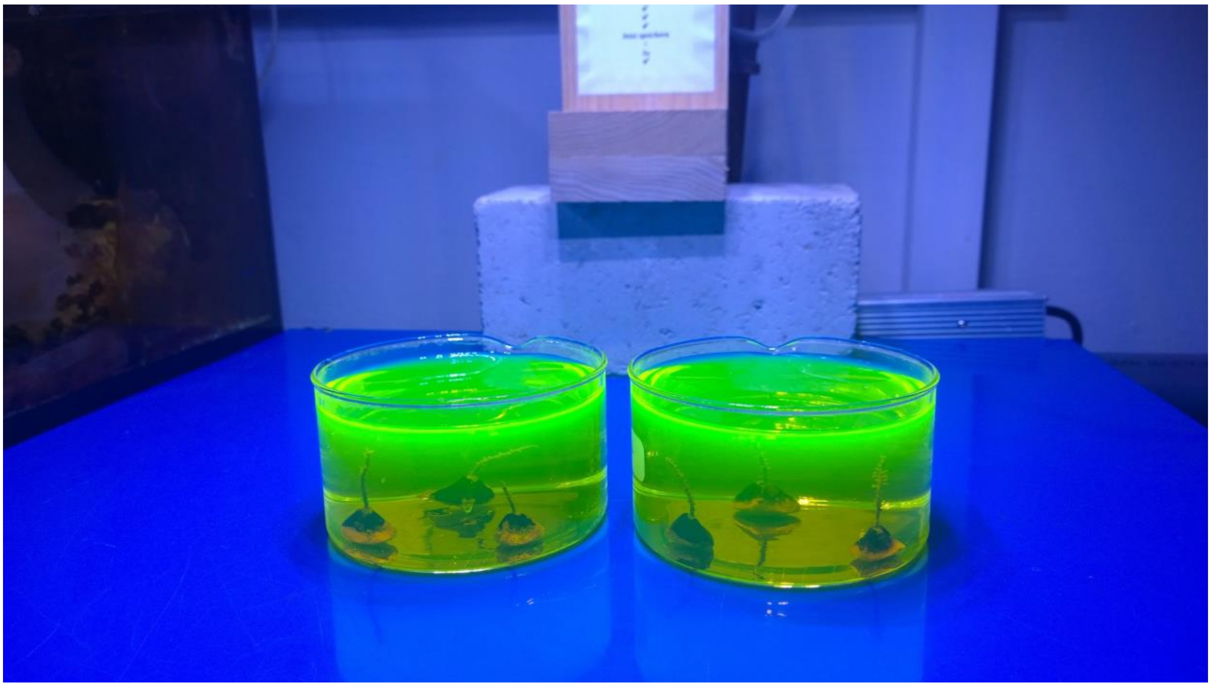

Supplementary Figure 1: Example of a calcein staining experiment. We kept corals for 72 hours in calcein under constant aeration (air pump removed for the photo) at room temperature.
