## Supplementary Figures for "Transcriptional response of the calcification and stress response toolkits in an octocoral under heat and pH stress": Supplementary_Figures_2.pdf

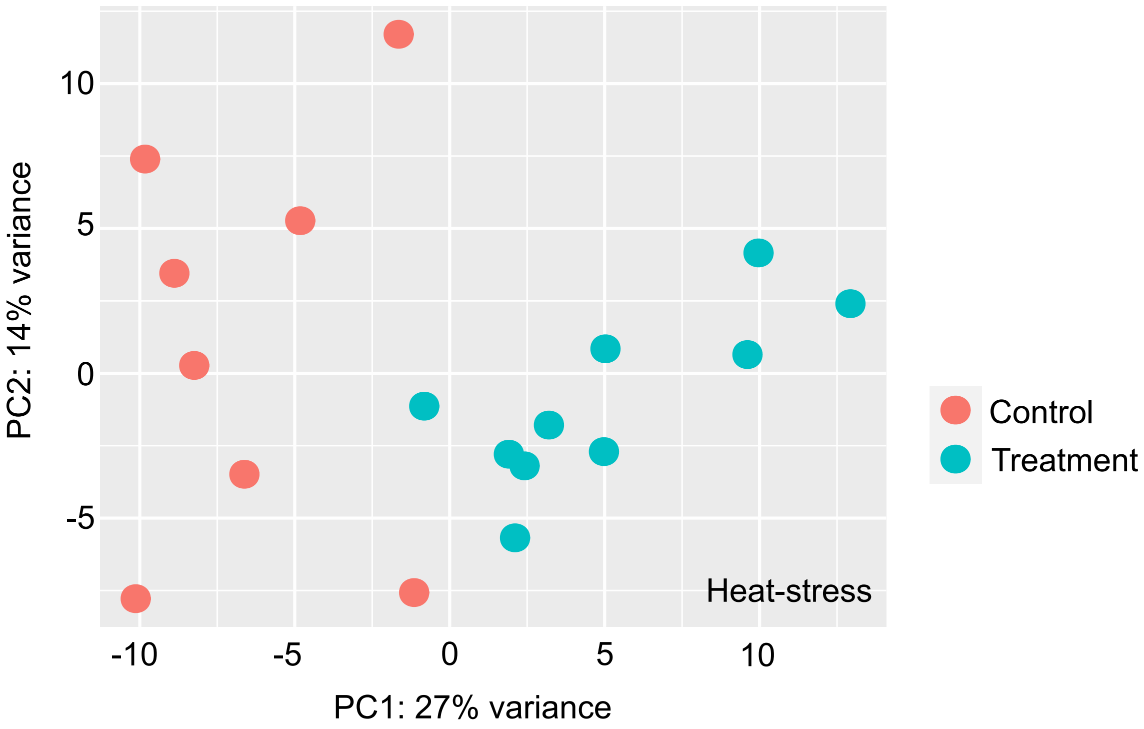


Supplementary Figure 2: Principal Component Analysis showing the global gene expression response of *Pinnigorgia flava* colonies exposed to heat stress.
