## Supplementary Figures for "Transcriptional response of the calcification and stress response toolkits in an octocoral under heat and pH stress": Supplementary_Figures_3-16.pdf

Chord plots (Supplementary Figures 3, 5, 7, 9, 11, 13 and 15) showing the semantic similarity within clusters of GO-terms enriched in the set of significantly overexpressed genes in *P. flava* colonies exposed to heat stress. For each cluster of semantically similar GO-terms, a directed acyclic subgraph (DAsG; Supplementary Figures 4, 6, 8, 10, 12, 14, 16) showing the relation between the GO-terms within a cluster in the general Biological Process gene ontology.

Supplementary Figure 3: Chord plot for cluster “darkblue”
Supplementary Figure 4: DAsG for cluster “darkblue”

Supplementary Figure 5: Chord plot for cluster “darkgreen”
Supplementary Figure 6: DAsG for cluster “darkgreen”

Supplementary Figure 7: Chord plot for cluster “mediumorchid3”
Supplementary Figure 8: DAsG for cluster “mediumorchid3”

Supplementary Figure 9: Chord plot for cluster “orange3”
Supplementary Figure 10: DAsG for cluster “orange3”

Supplementary Figure 11: Chord plot for cluster “firebrick4”
Supplementary Figure 12: DAsG for cluster “firebrick4”

Supplementary Figure 13: Chord plot for cluster “gray40”
Supplementary Figure 14: DAsG for cluster “gray40”

Supplementary Figure 15: Chord plot for cluster “violetred2”
Supplementary Figure 16: DAsG for cluster “violetred2”

NOTE: cluster names refer to the Rcolor used to plot them in the dendrogram.

**Temp. OverDEGs Cluster darkblue**

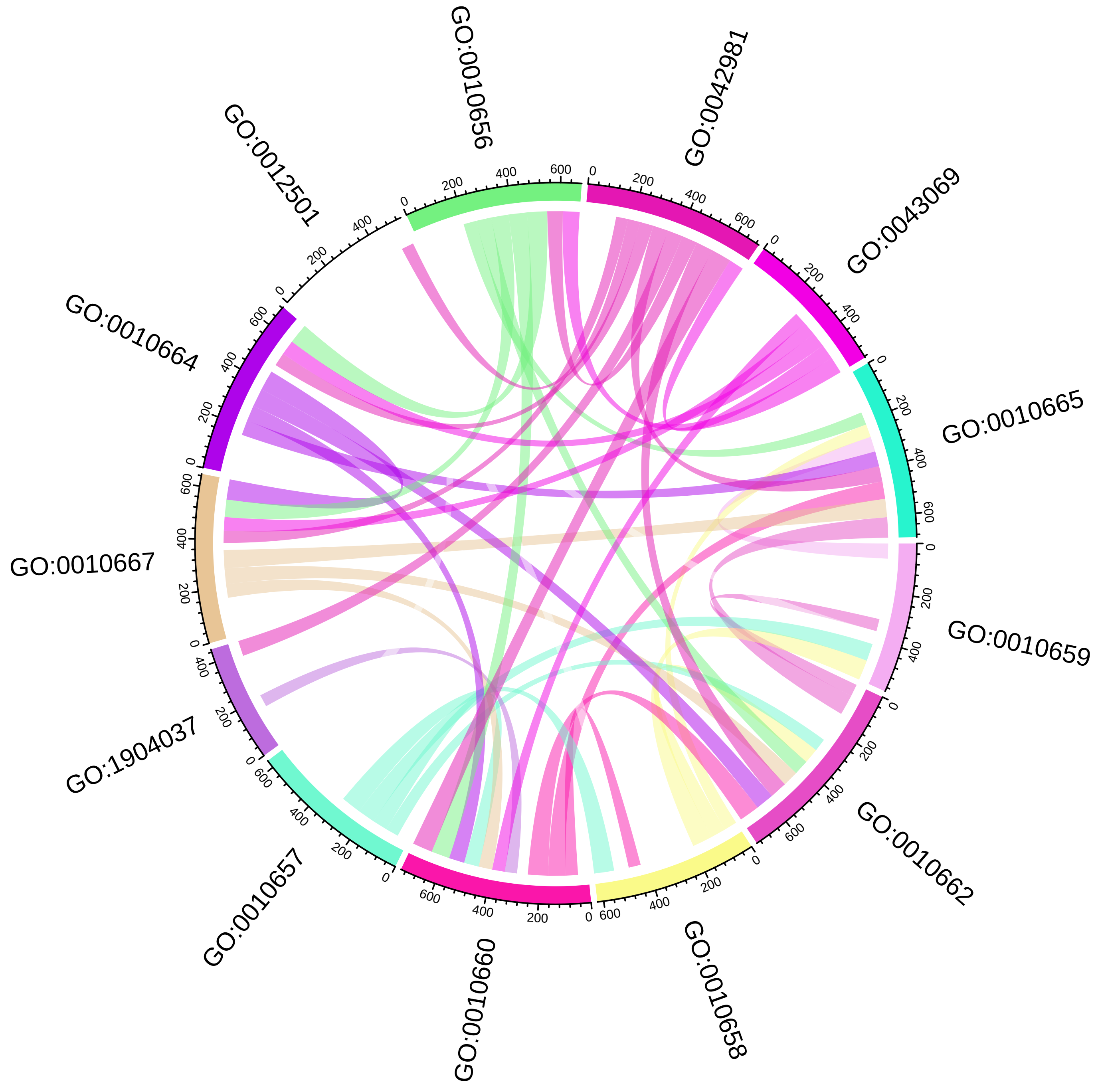

**Temp. OverDEGs Cluster darkblue**

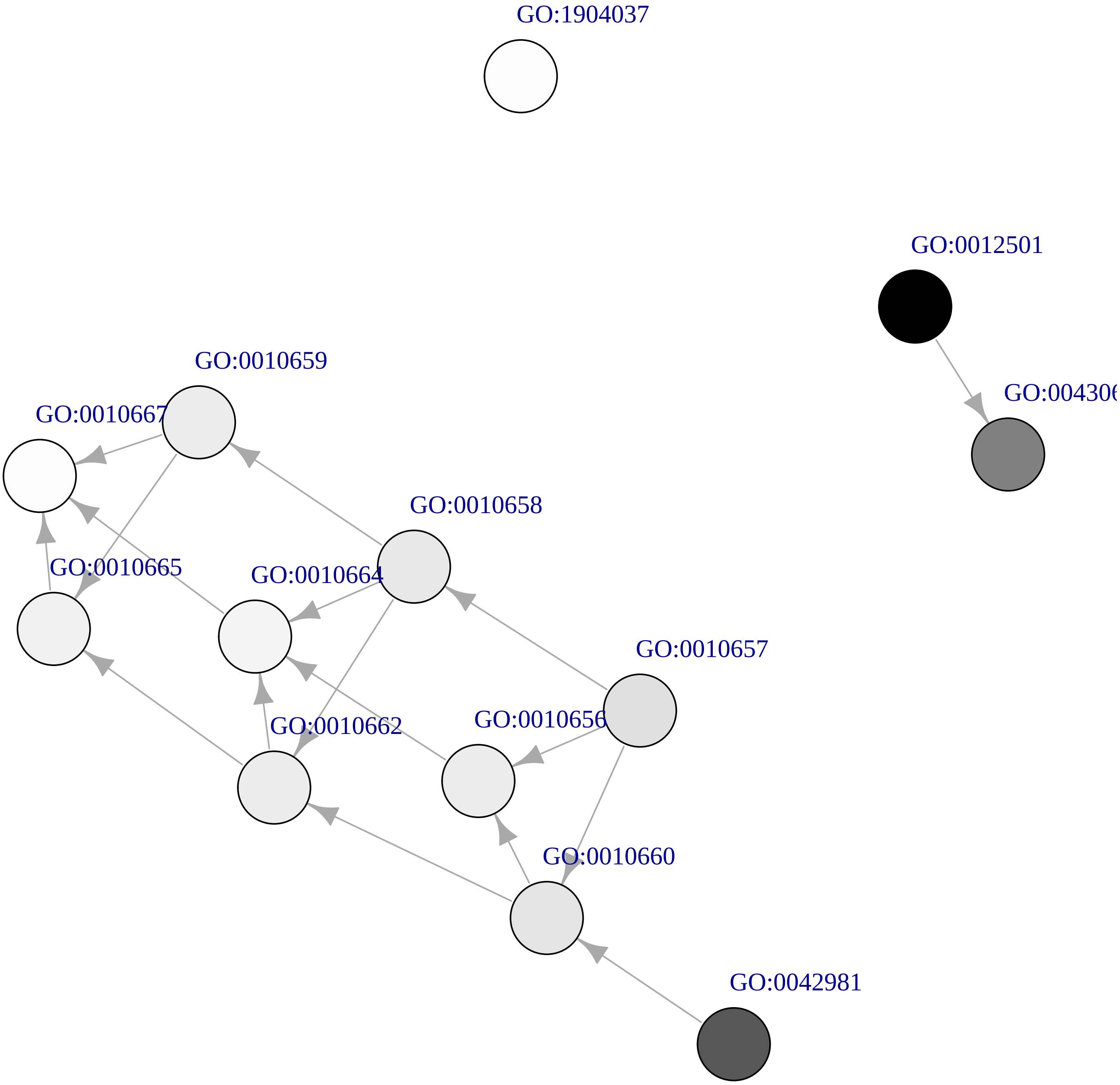

### Temp. OverDEGs Cluster darkgreen

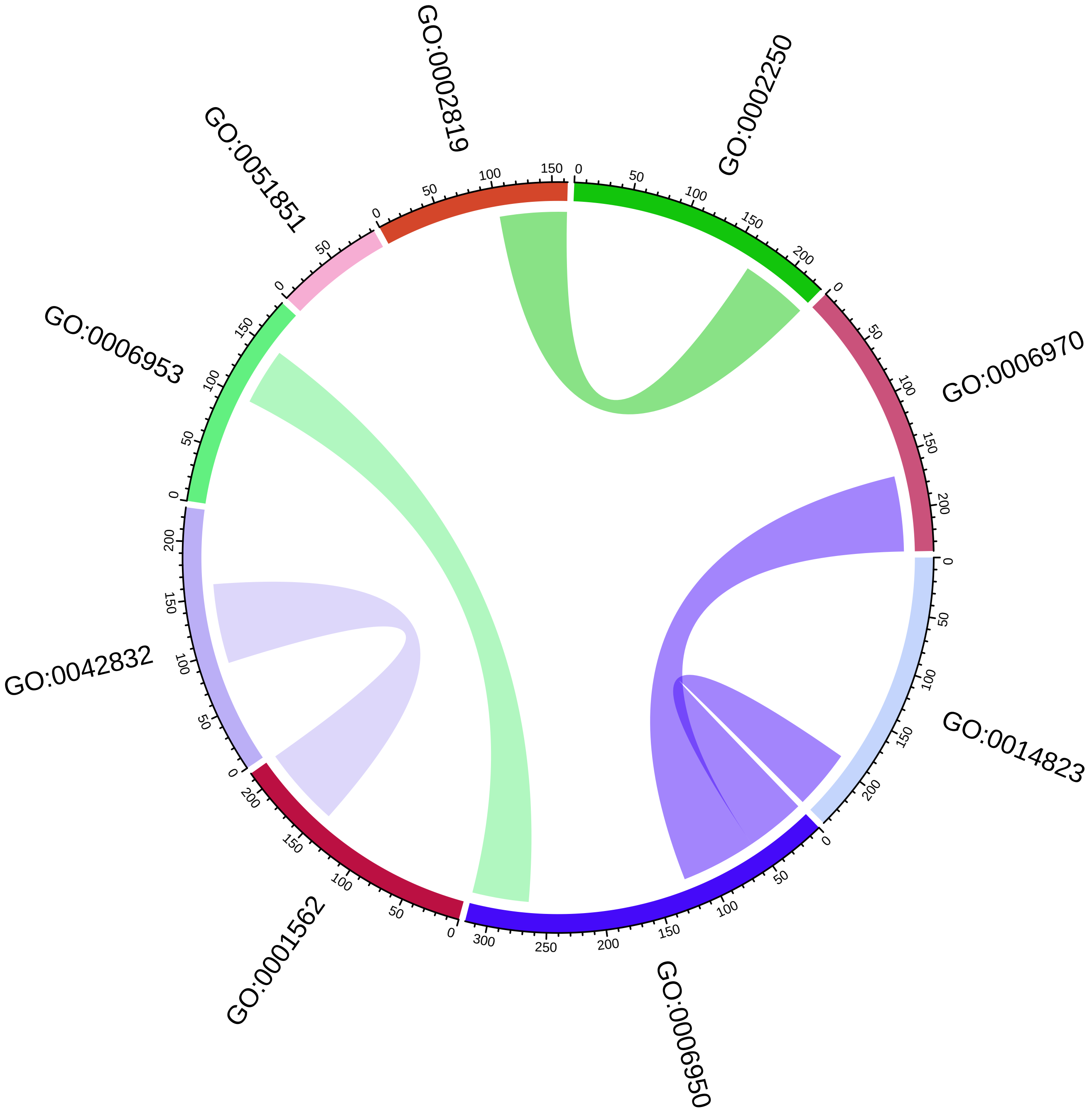

**Temp. OverDEGs Cluster darkgreen**

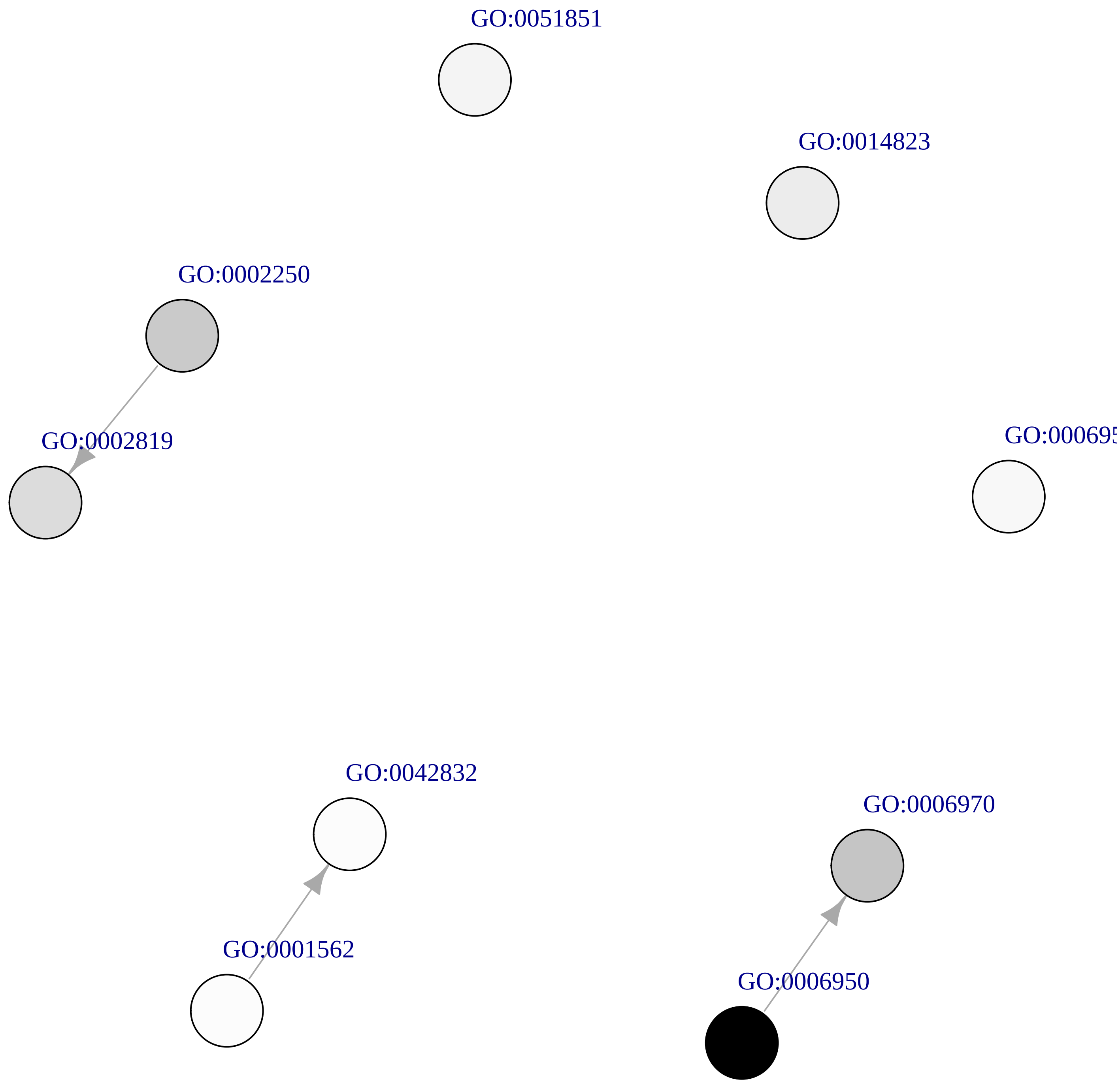

#### Temp. OverDEGs Cluster mediumorchid3

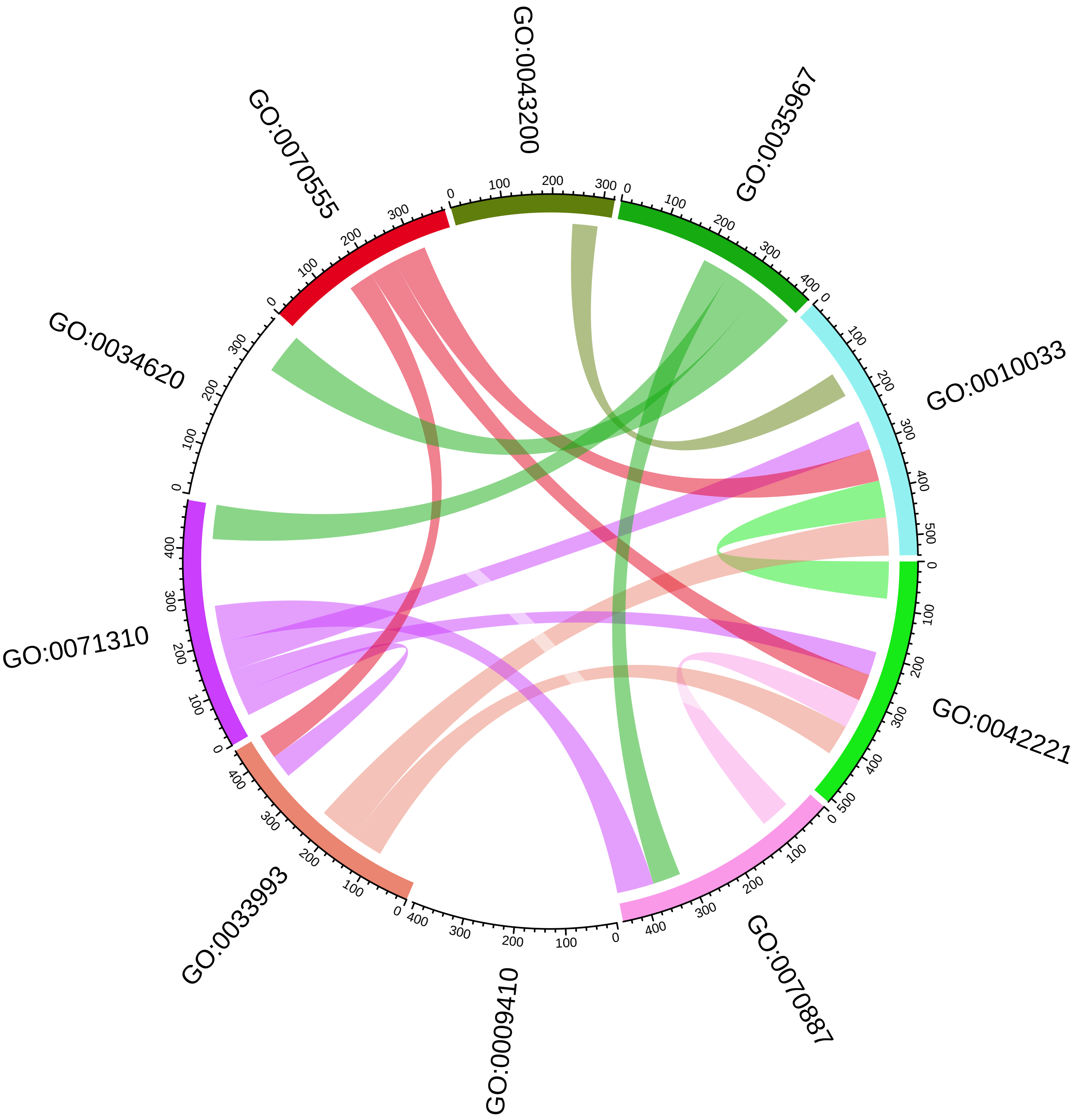

### Temp. OverDEGs Cluster mediumorchid3

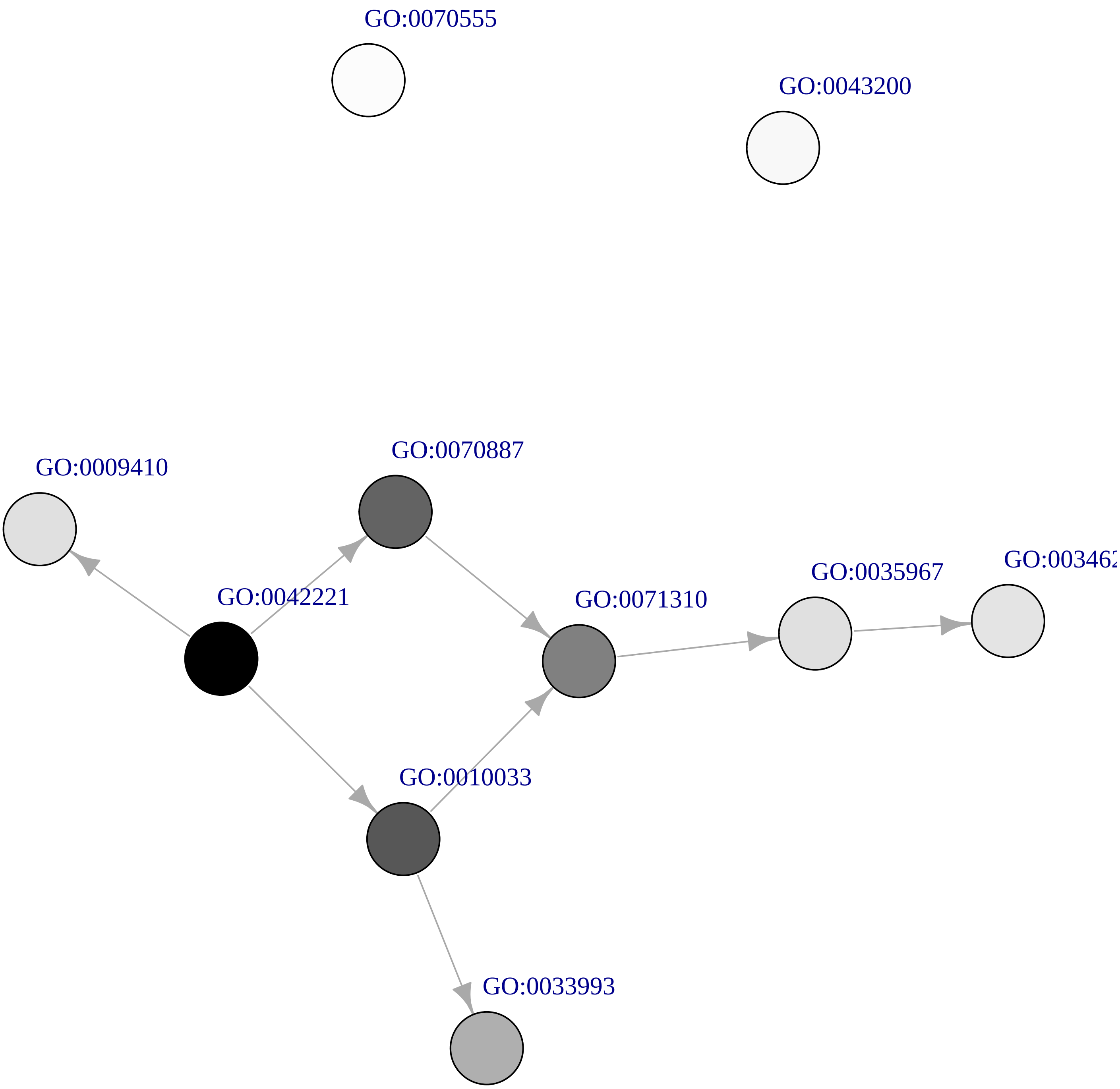

### Temp. OverDEGs Cluster orange3

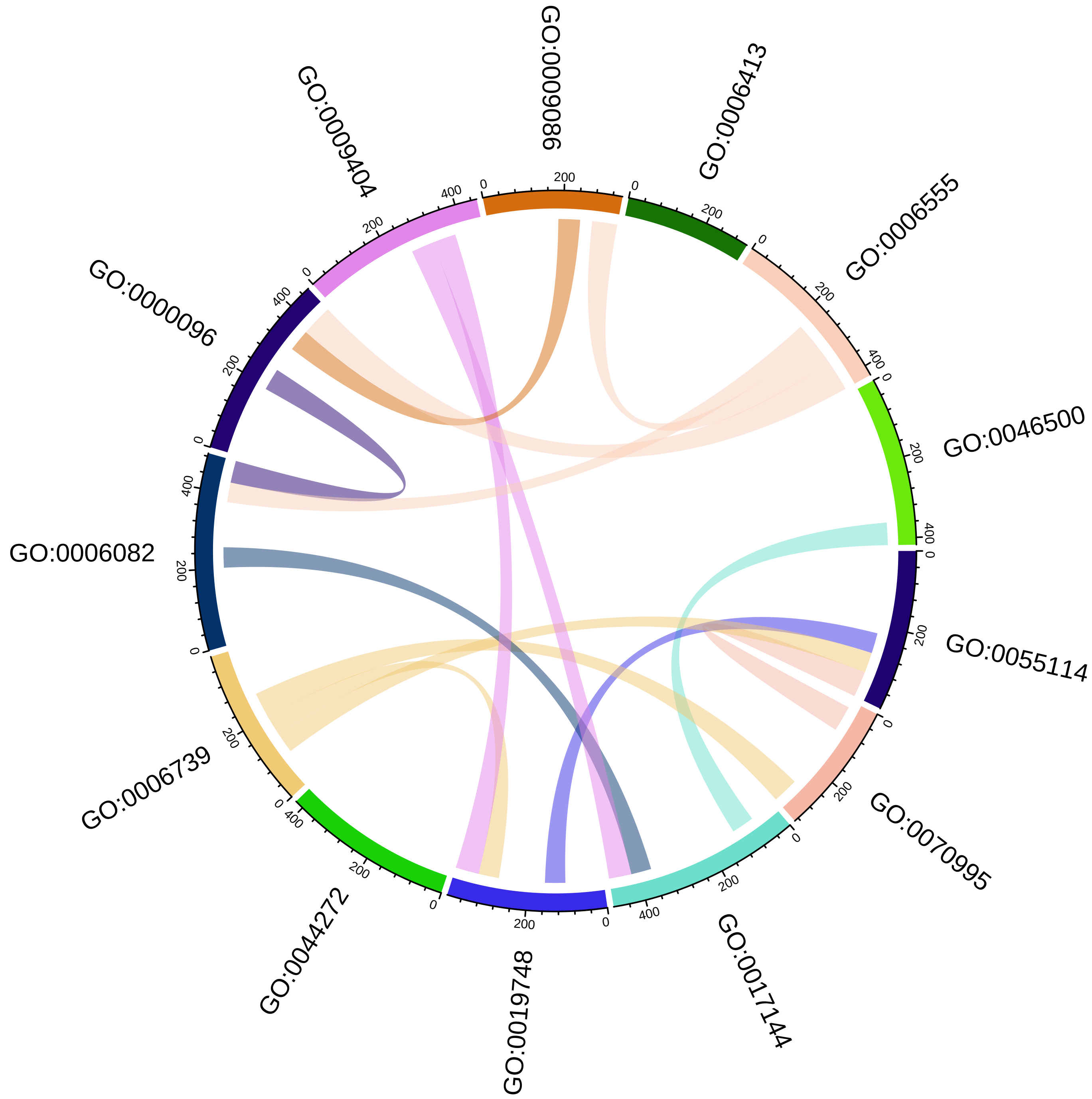

#### Temp. OverDEGs Cluster orange3

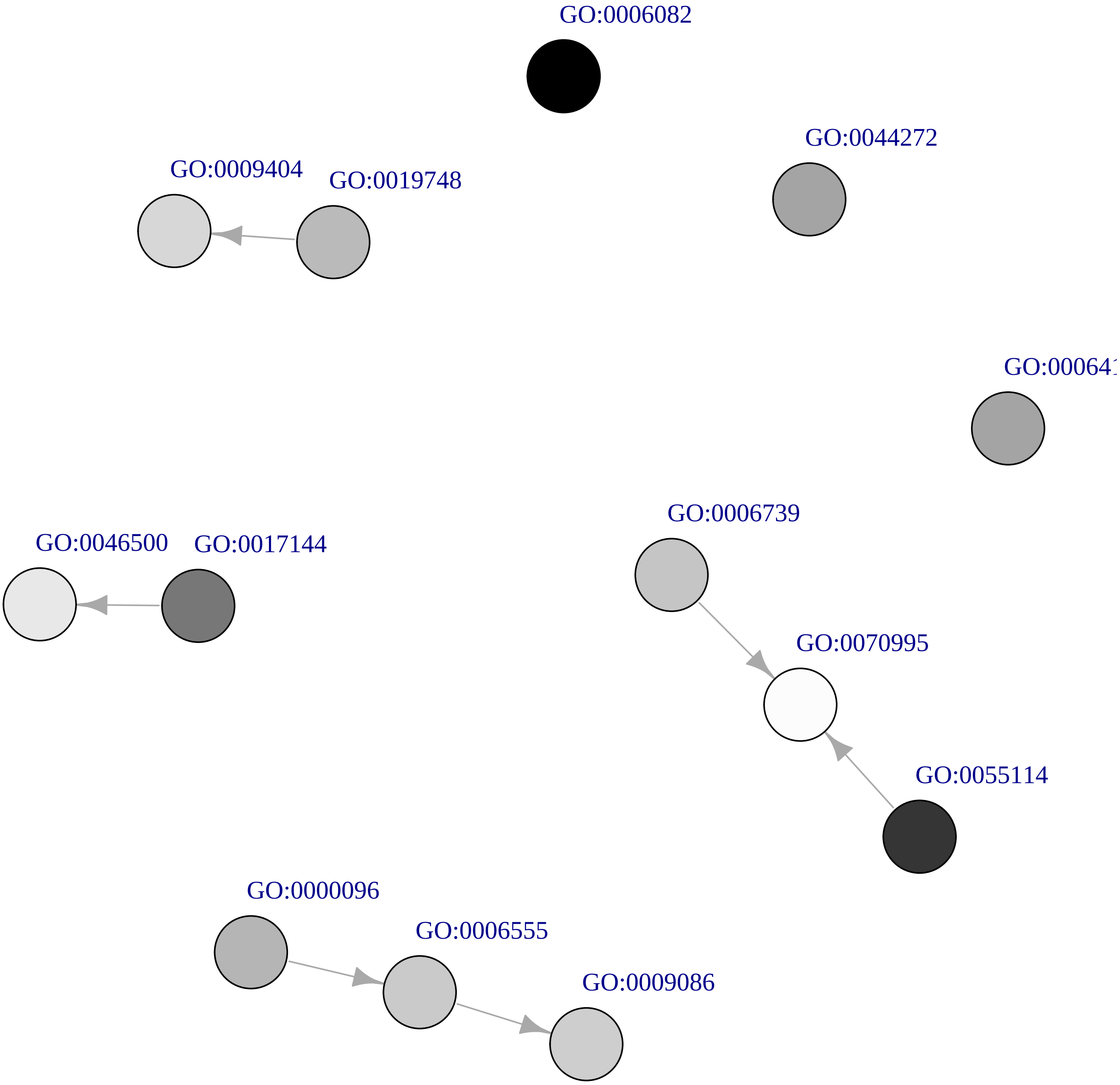

### Temp. OverDEGs Cluster firebrick4

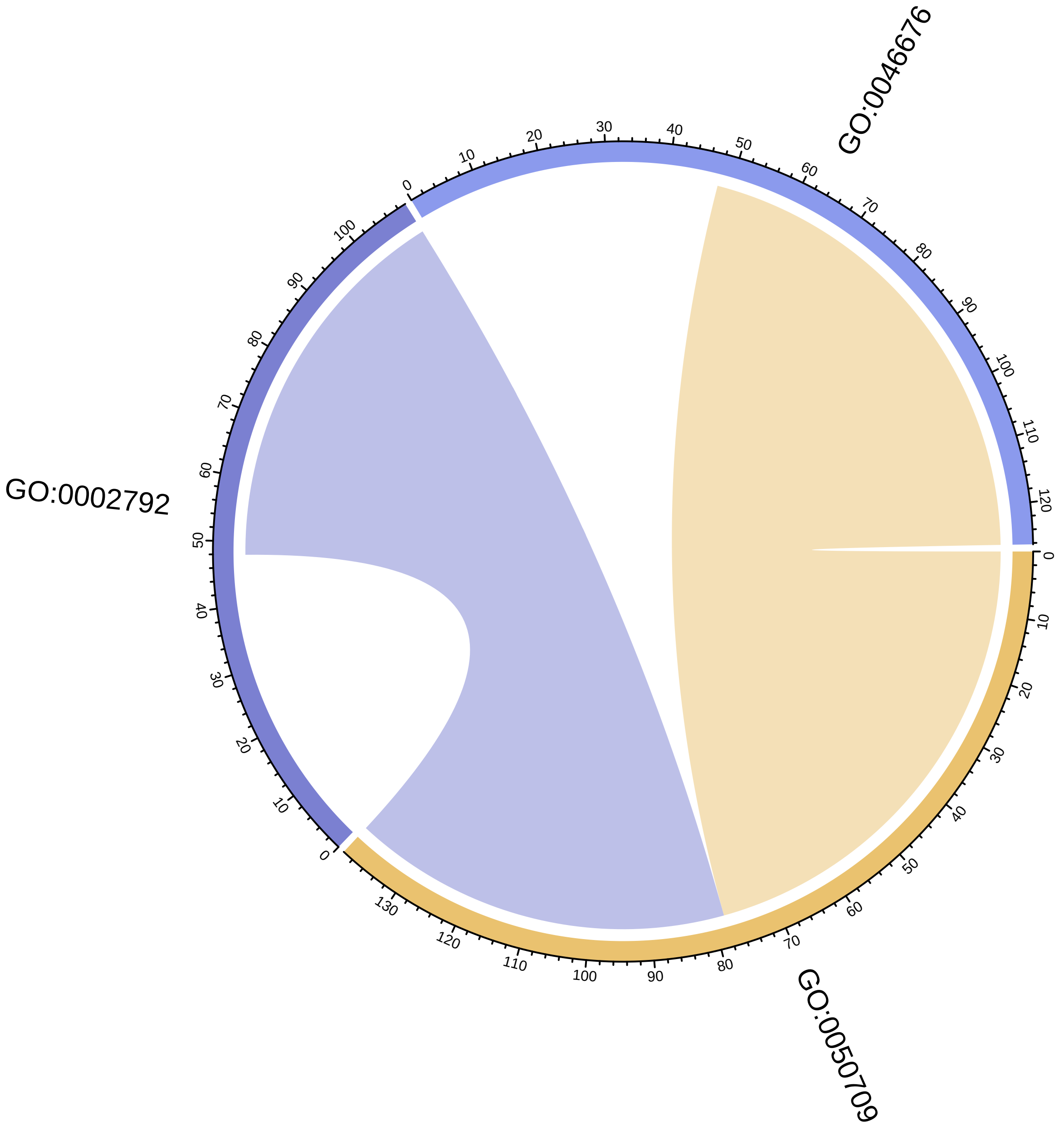

#### Temp. OverDEGs Cluster firebrick4

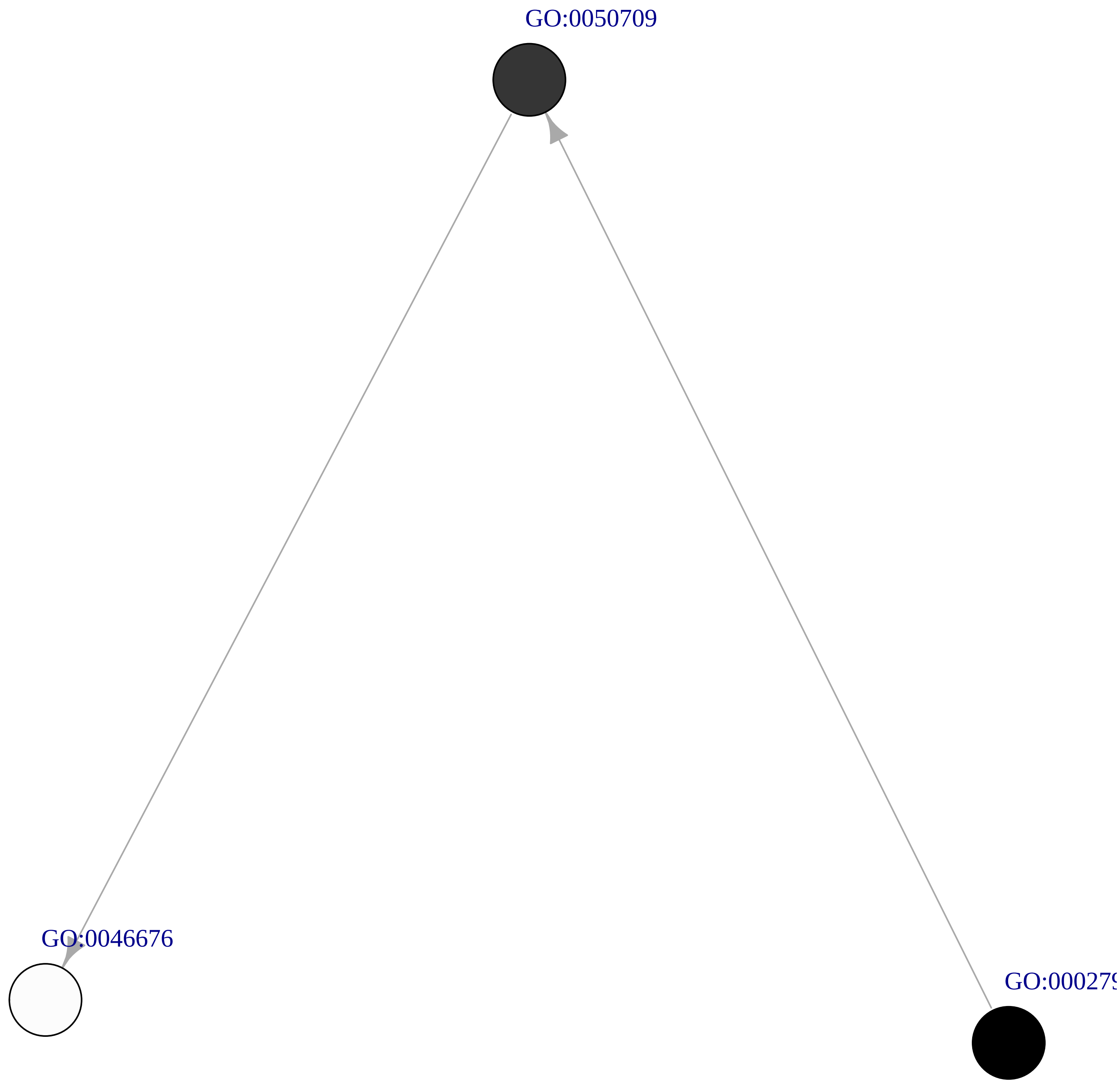

### Temp. OverDEGs Cluster gray40

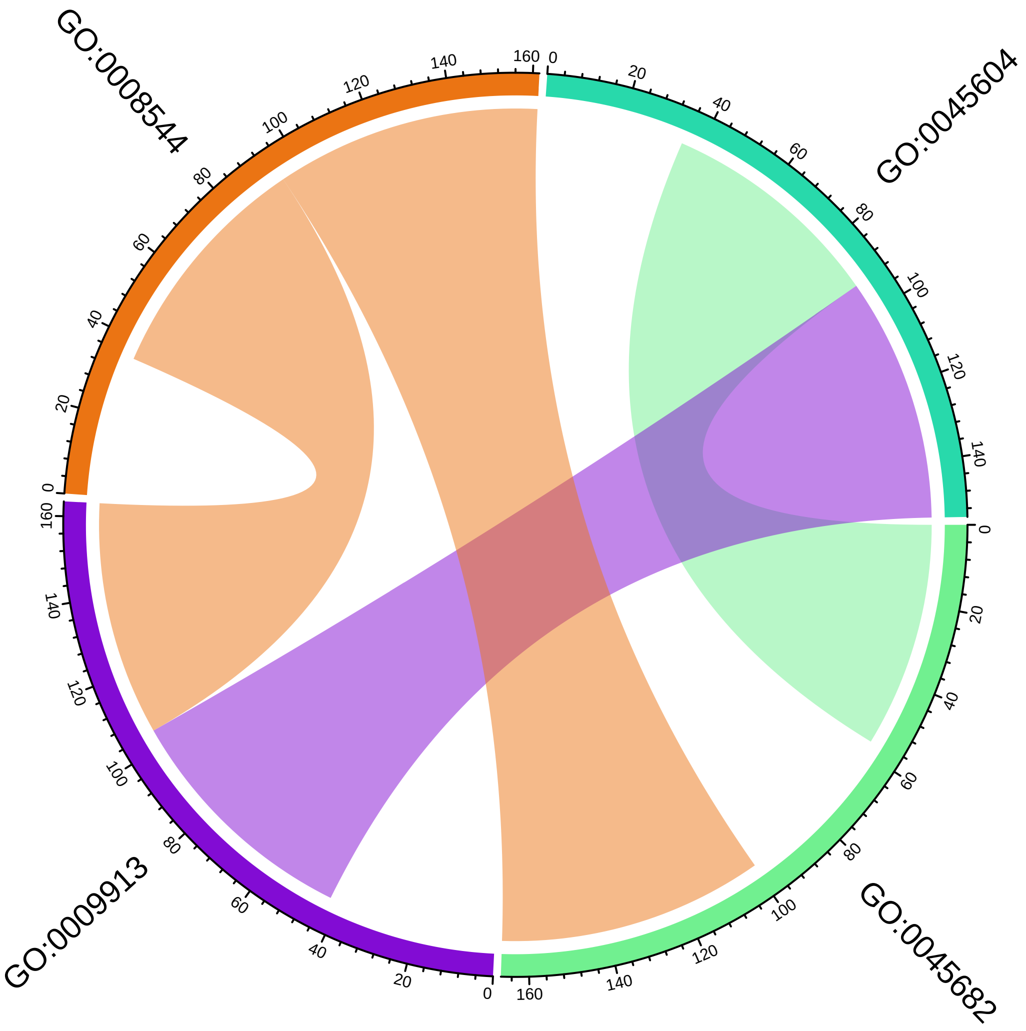

**Temp. OverDEGs Cluster gray40**

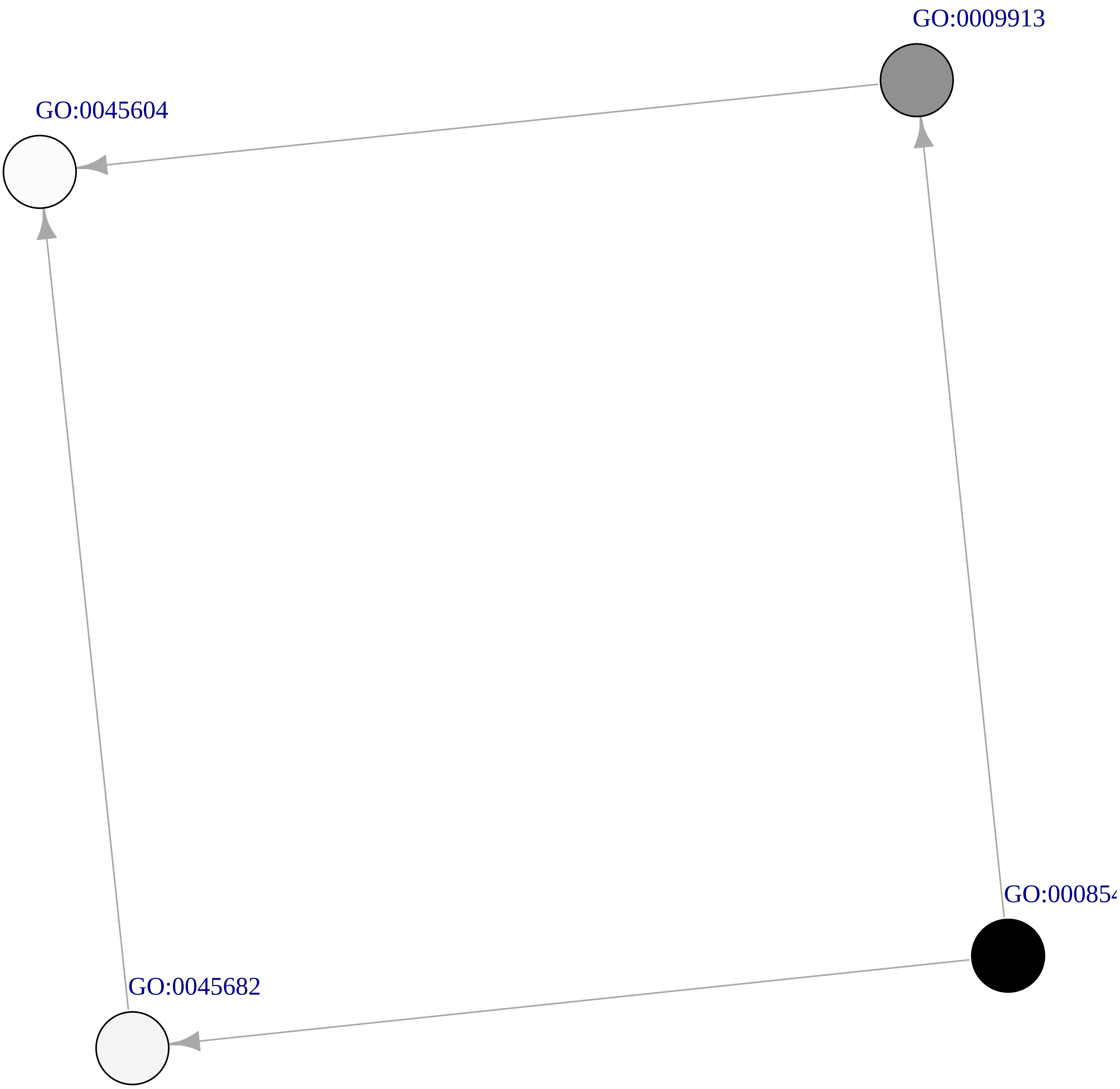

**Temp. OverDEGs Cluster violetred2**

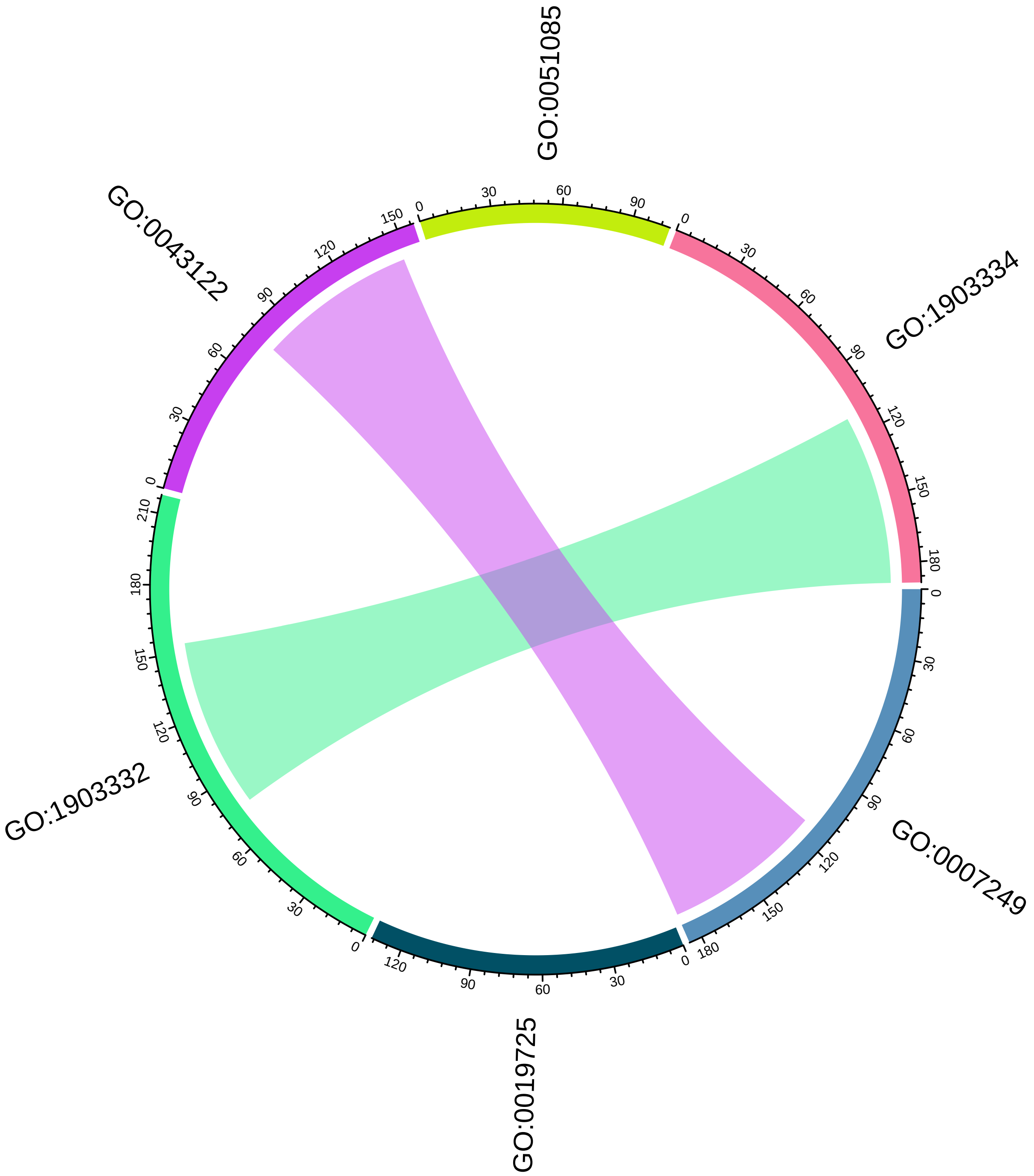

Temp. OverDEGs Cluster violetred2

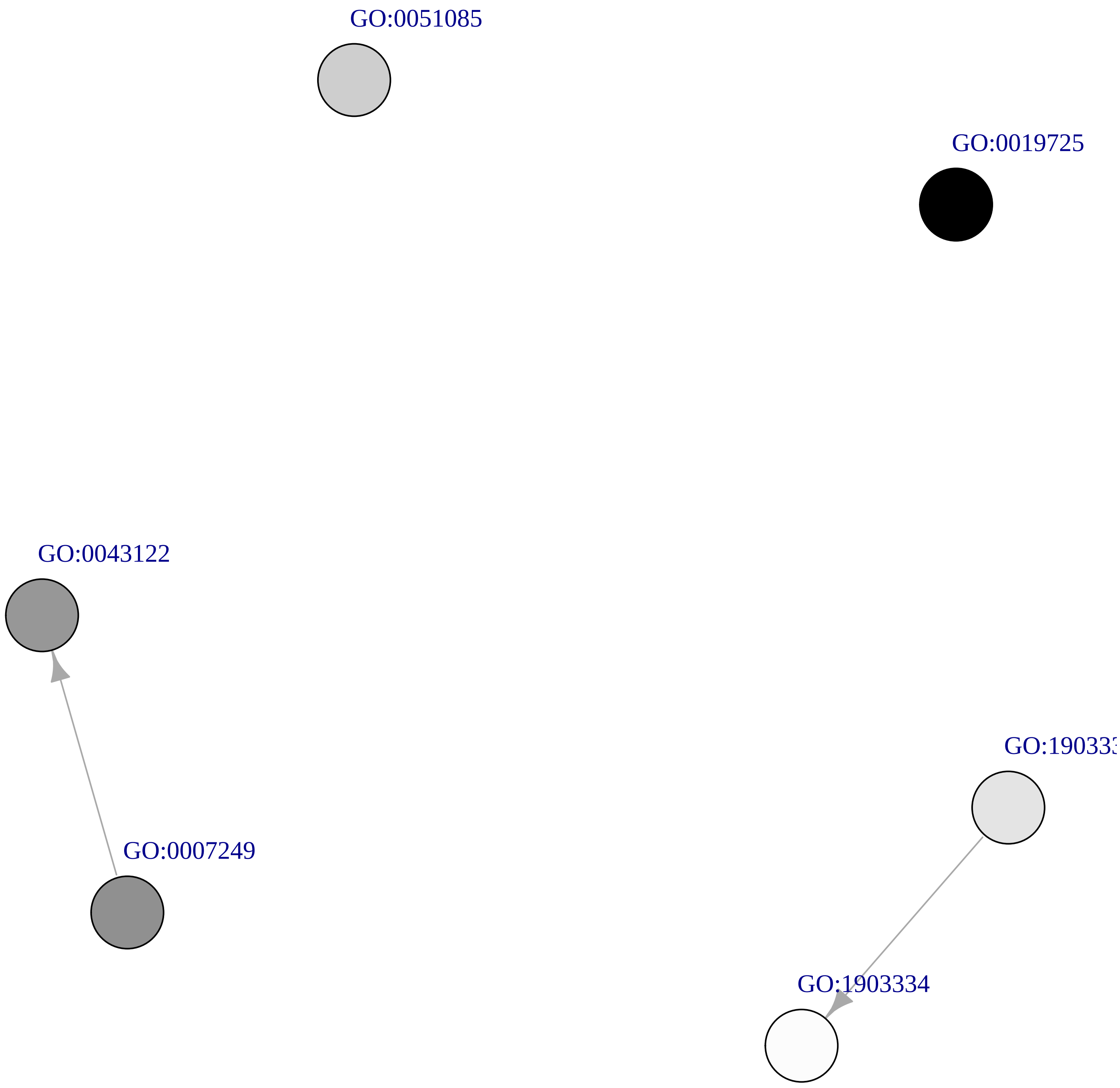
