## Supplementary Figures for "Transcriptional response of the calcification and stress response toolkits in an octocoral under heat and pH stress": Supplementary_Figures_17-30.pdf

Chord plots (Supplementary Figures 17, 19, 21, 23, 25, 27 and 29) showing the semantic similarity within clusters of GO-terms enriched in the set of significantly underexpressed genes in *P. flava* colonies exposed to heat stress. For each cluster of semantically similar GO-terms, a directed acyclic subgraph (DAsG; Supplementary Figures 18, 20, 22, 24, 26, 28, 30) showing the relation between the GO-terms within a cluster in the general Biological Process gene ontology.

Supplementary Figure 17: Chord plot for cluster “darkblue”
Supplementary Figure 18: DAsG for cluster “darkblue”

Supplementary Figure 19: Chord plot for cluster “darkgreen”
Supplementary Figure 20: DAsG for cluster “darkgreen”

Supplementary Figure 21: Chord plot for cluster “mediumorchid3”
Supplementary Figure 22: DAsG for cluster “mediumorchid3”

Supplementary Figure 23: Chord plot for cluster “orange3”
Supplementary Figure 24: DAsG for cluster “orange3”

Supplementary Figure 25: Chord plot for cluster “firebrick4”
Supplementary Figure 26: DAsG for cluster “firebrick4”

Supplementary Figure 27: Chord plot for cluster “gray40”
Supplementary Figure 28: DAsG for cluster “gray40”

Supplementary Figure 29: Chord plot for cluster “violetred2”
Supplementary Figure 30: DAsG for cluster “violetred2”

NOTE: cluster names refer to the Rcolor used to plot them in the dendrogram.

#### Temp. UnderDEGs Cluster darkblue

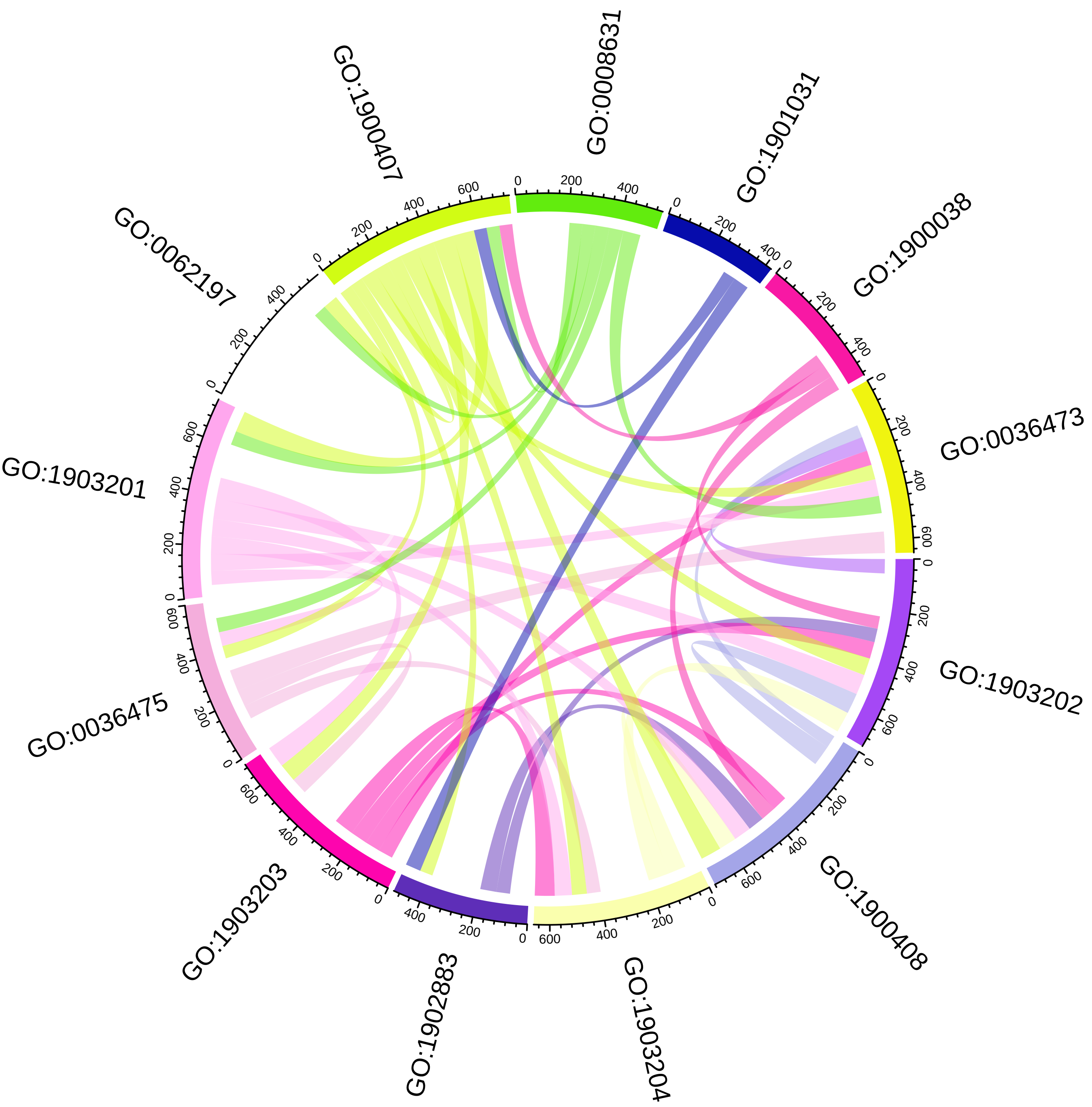

### Temp. UnderDEGs Cluster darkblue

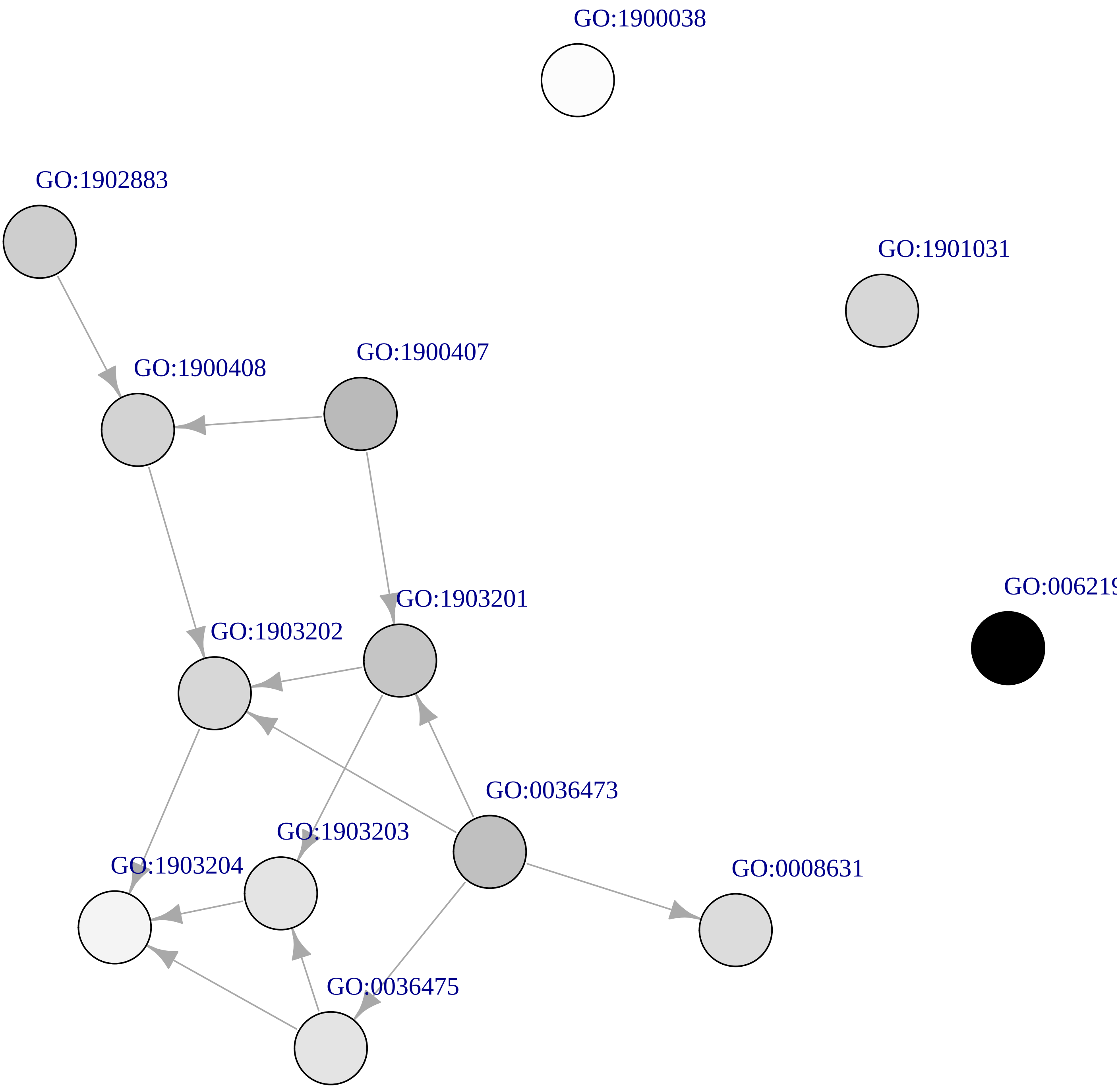

**Temp. UnderDEGs Cluster darkgreen**

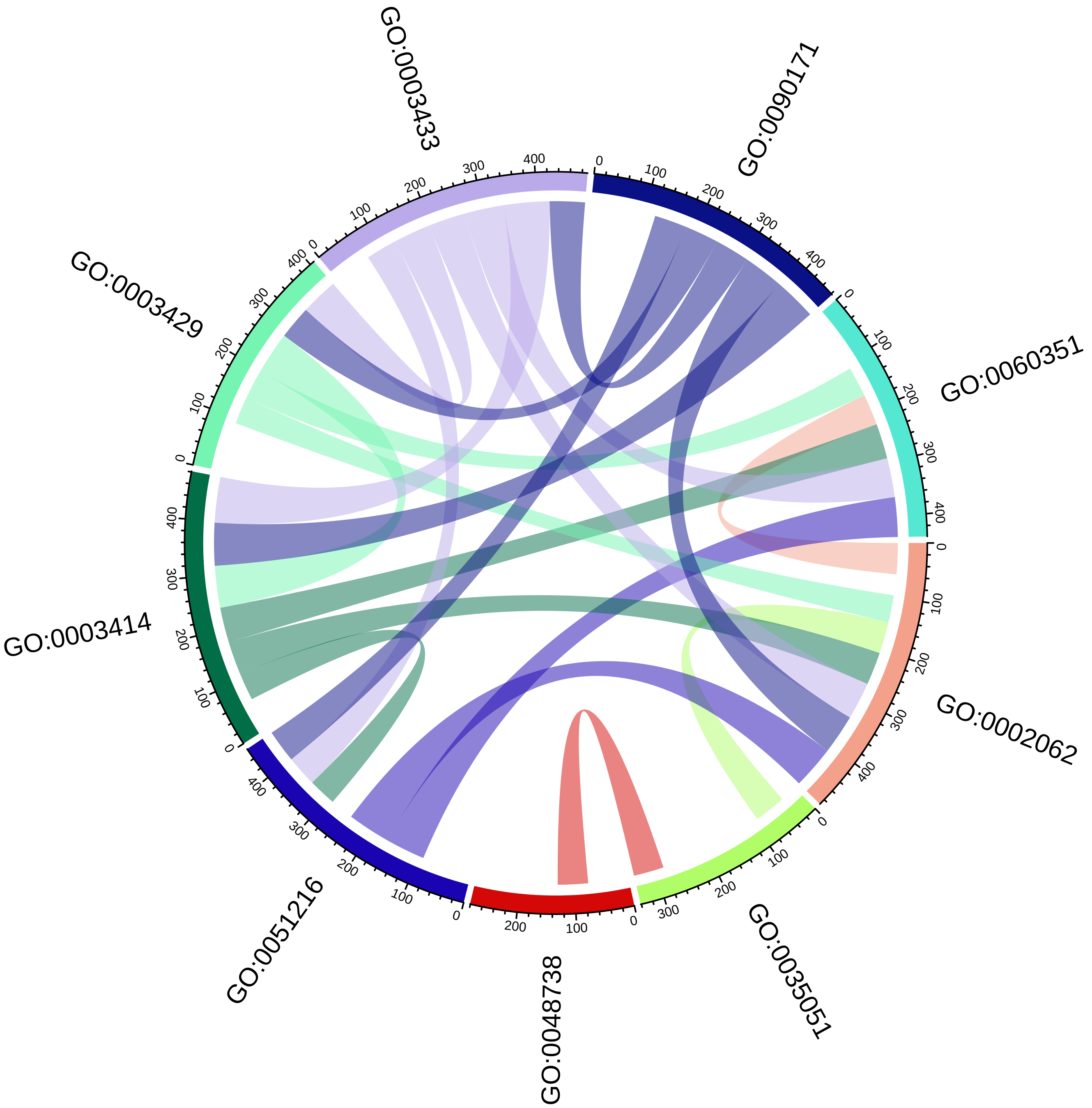

### Temp. UnderDEGs Cluster darkgreen

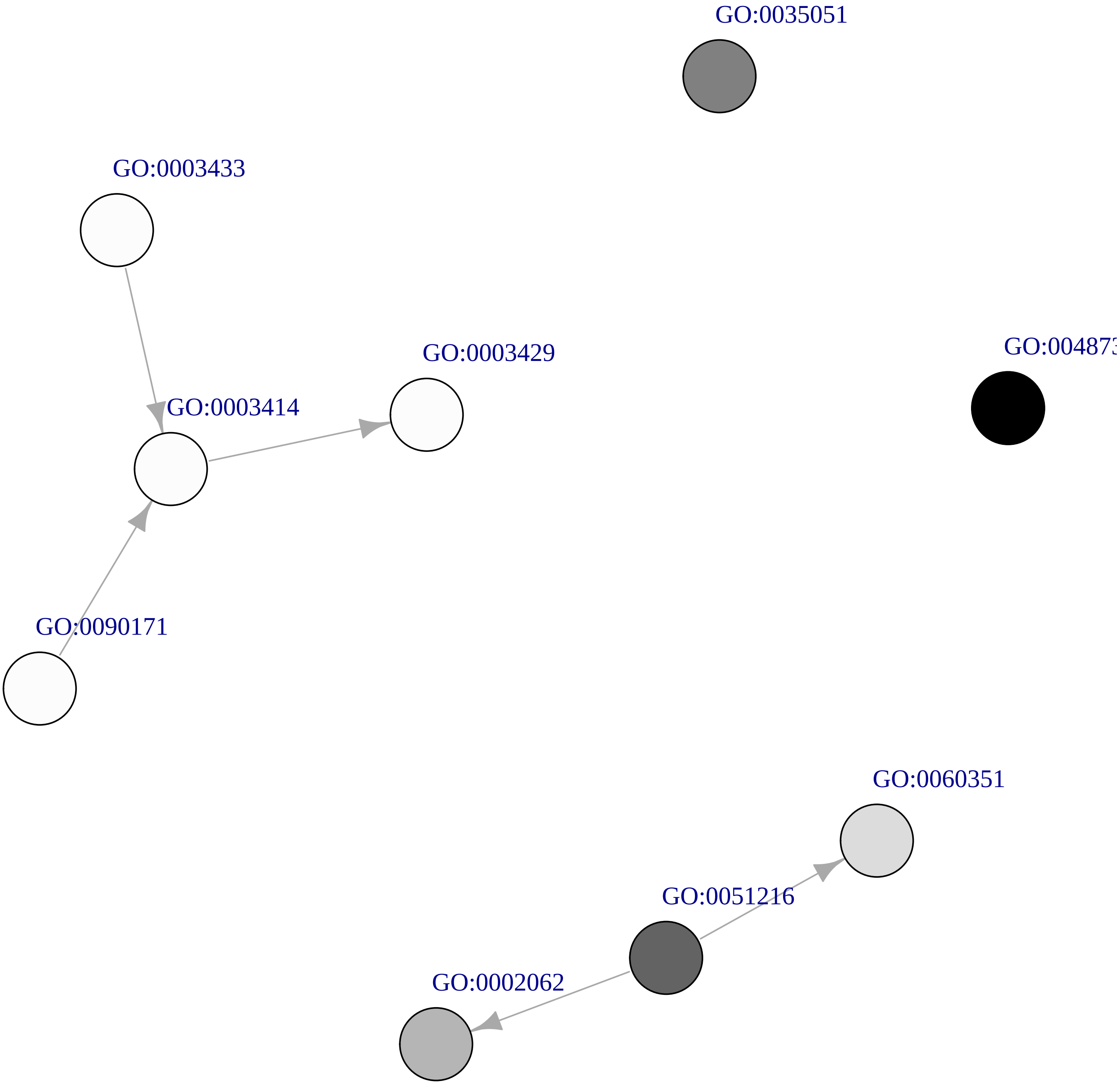

### Temp. UnderDEGs Cluster mediumorchid3

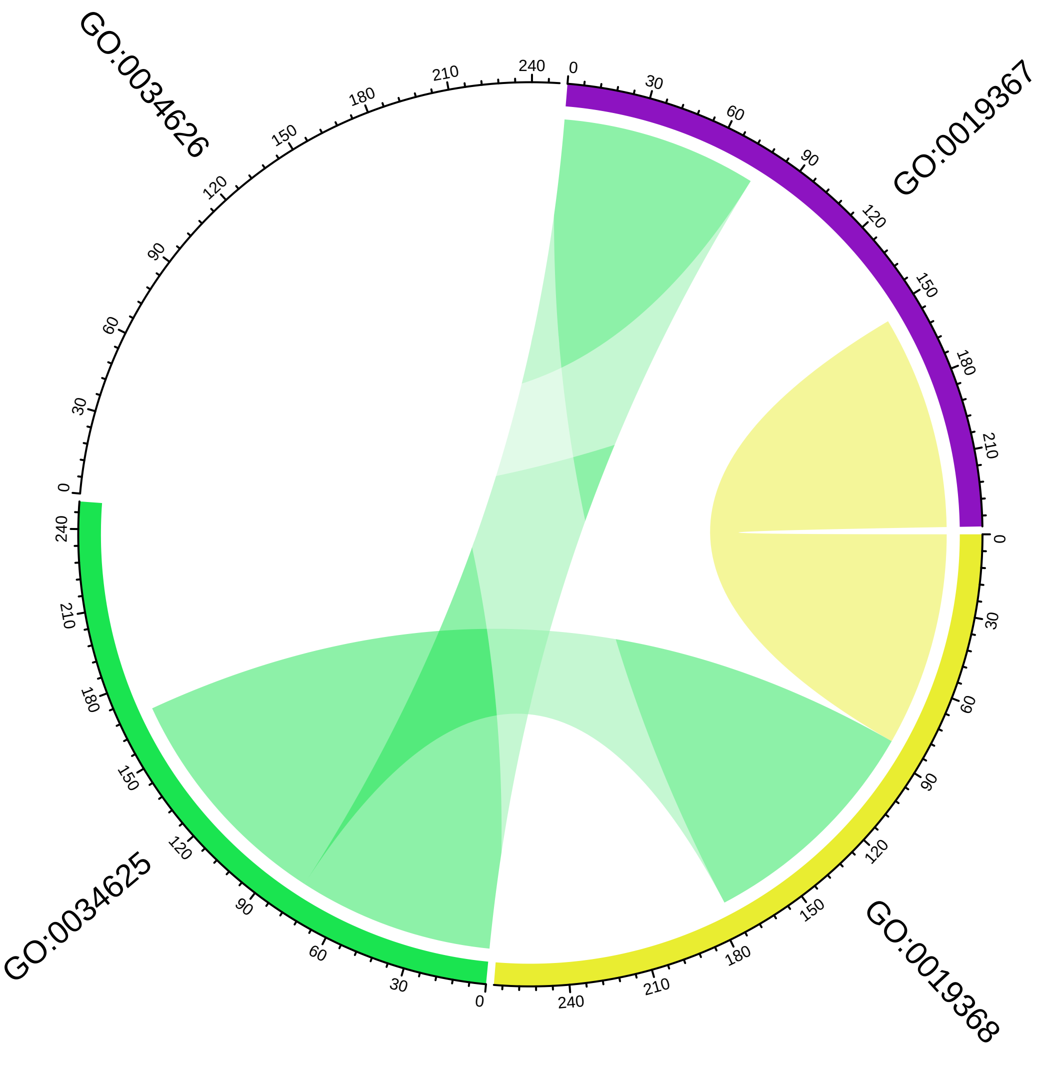

#### Temp. UnderDEGs Cluster mediumorchid3

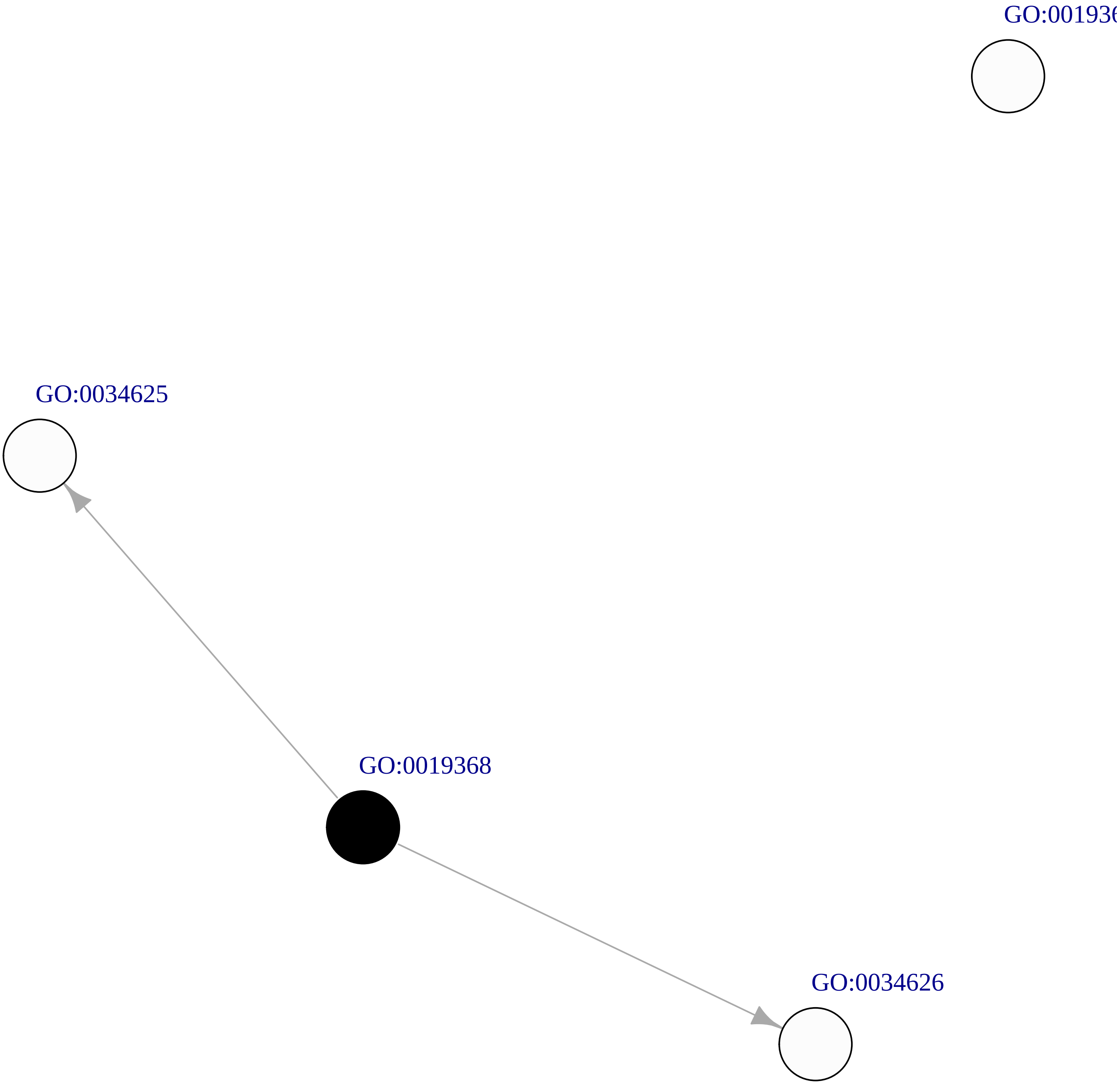

#### Temp. UnderDEGs Cluster orange3

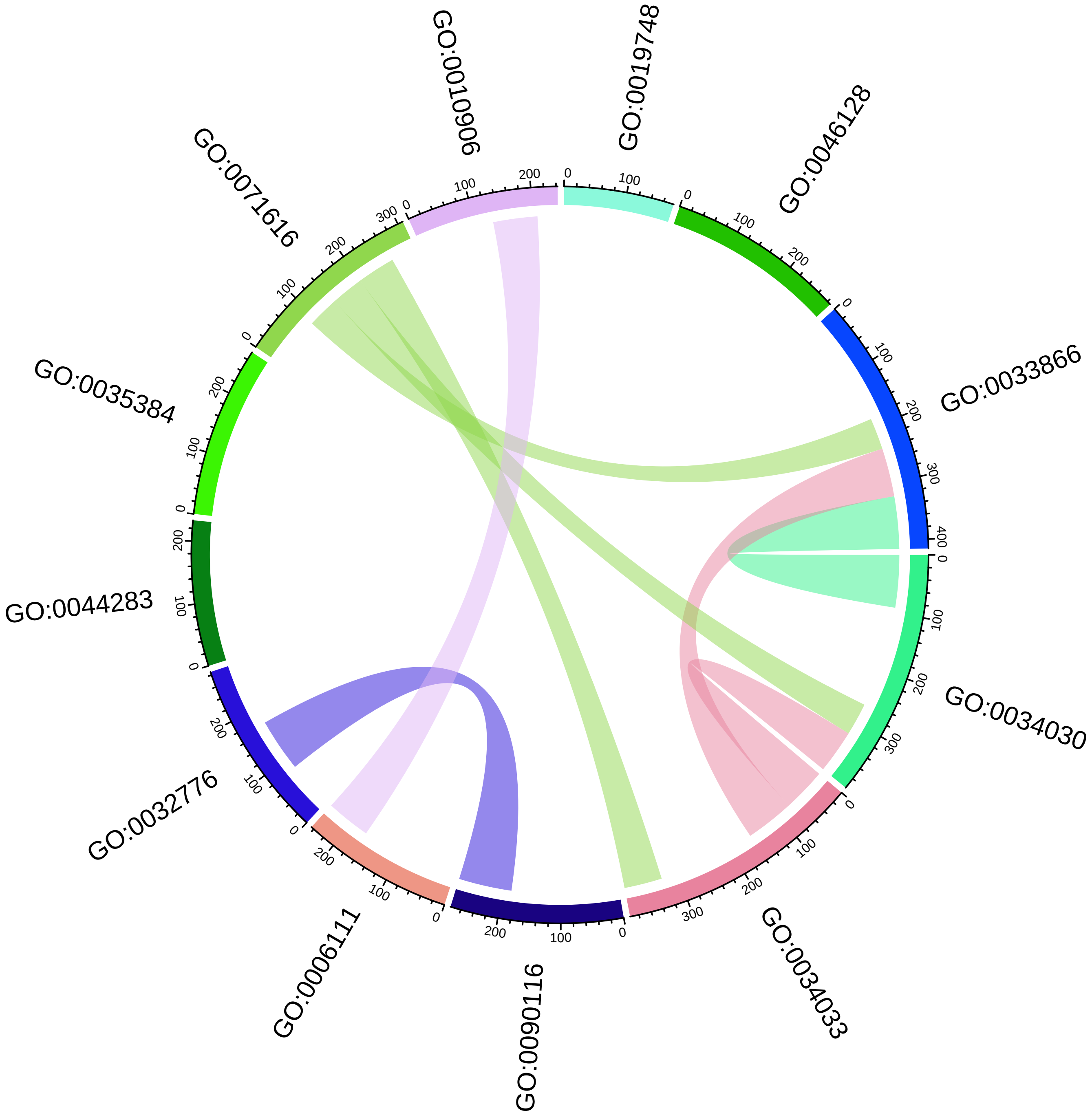

#### Temp. UnderDEGs Cluster orange3

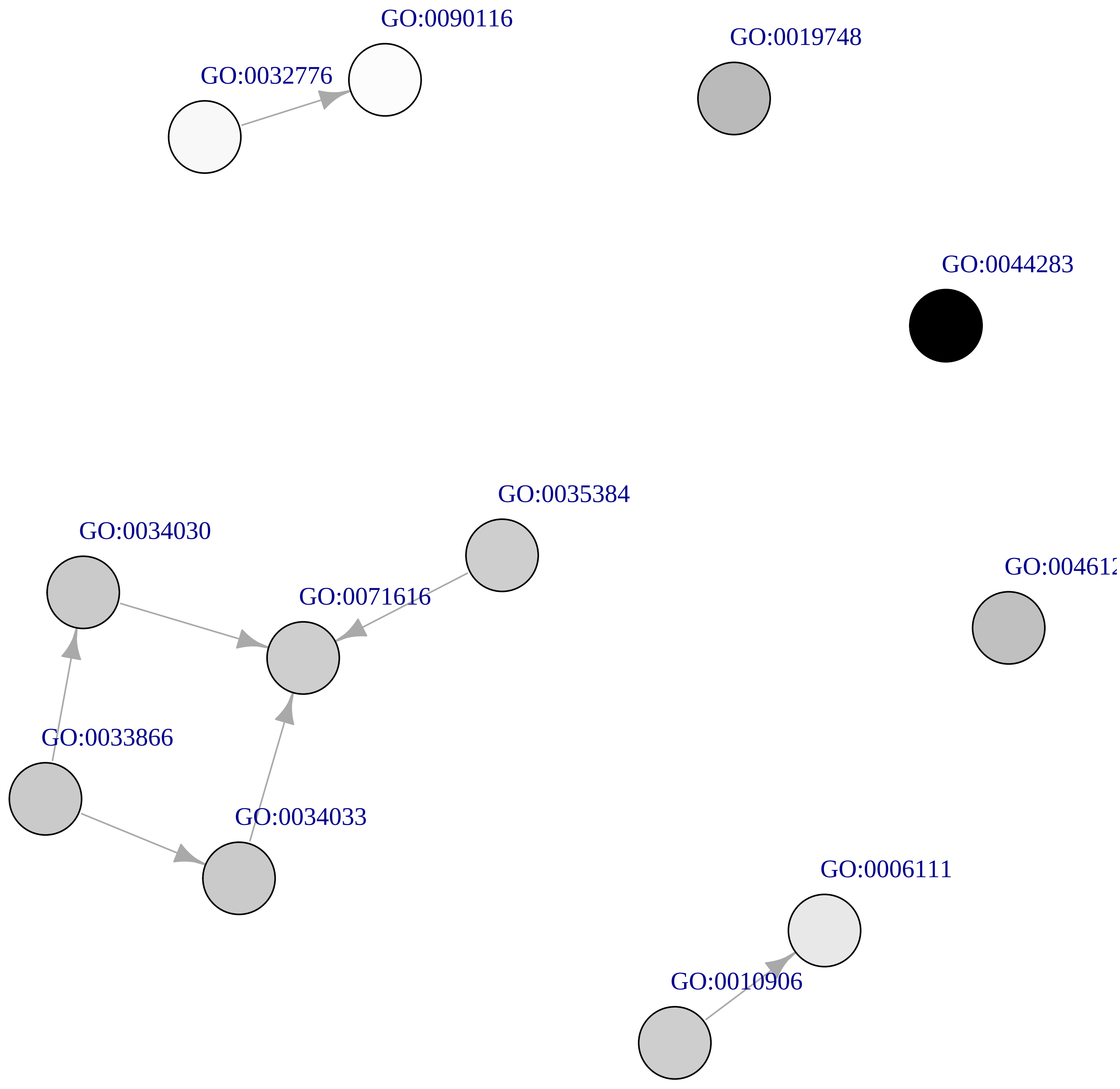

**Temp. UnderDEGs Cluster firebrick4**

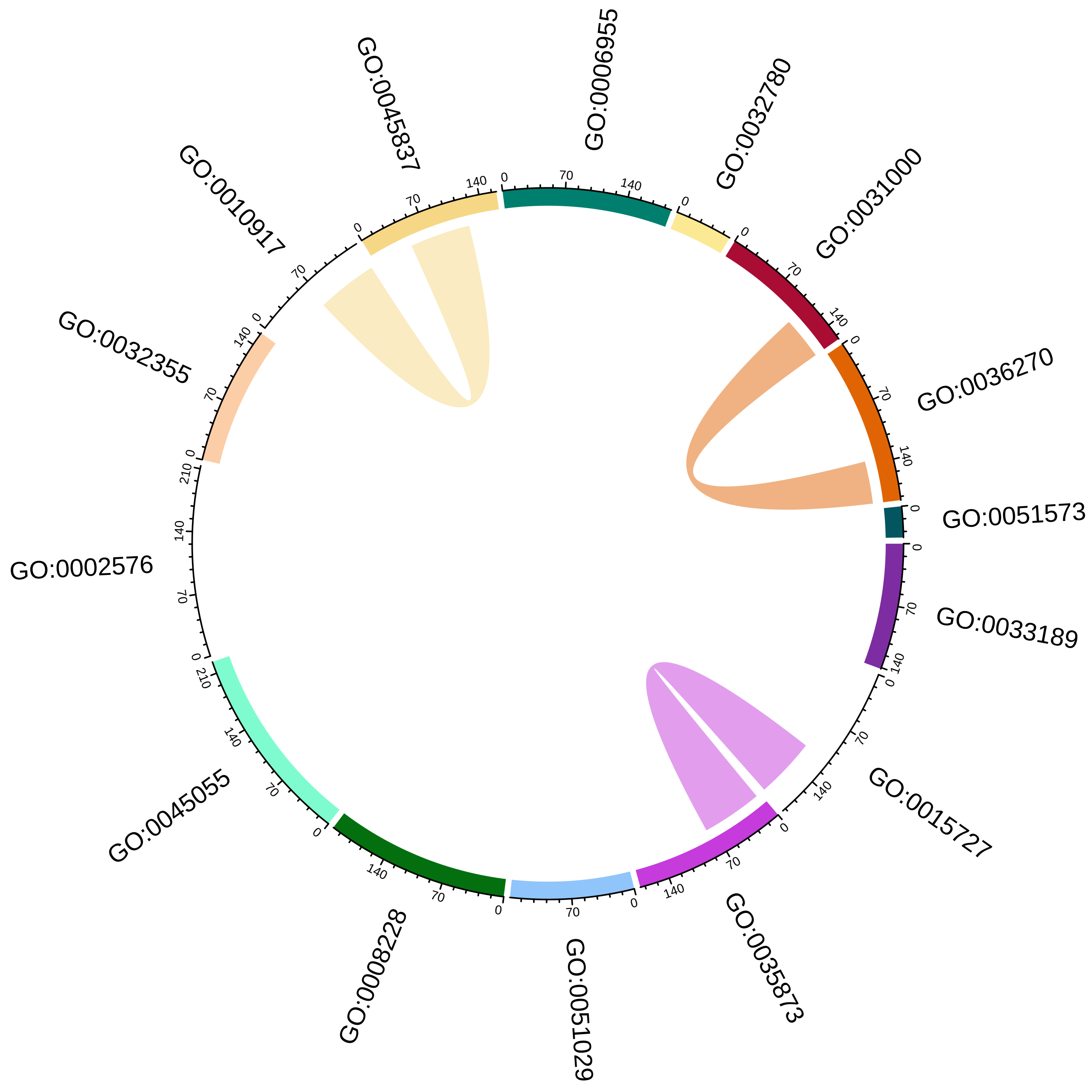

### Temp. UnderDEGs Cluster firebrick4

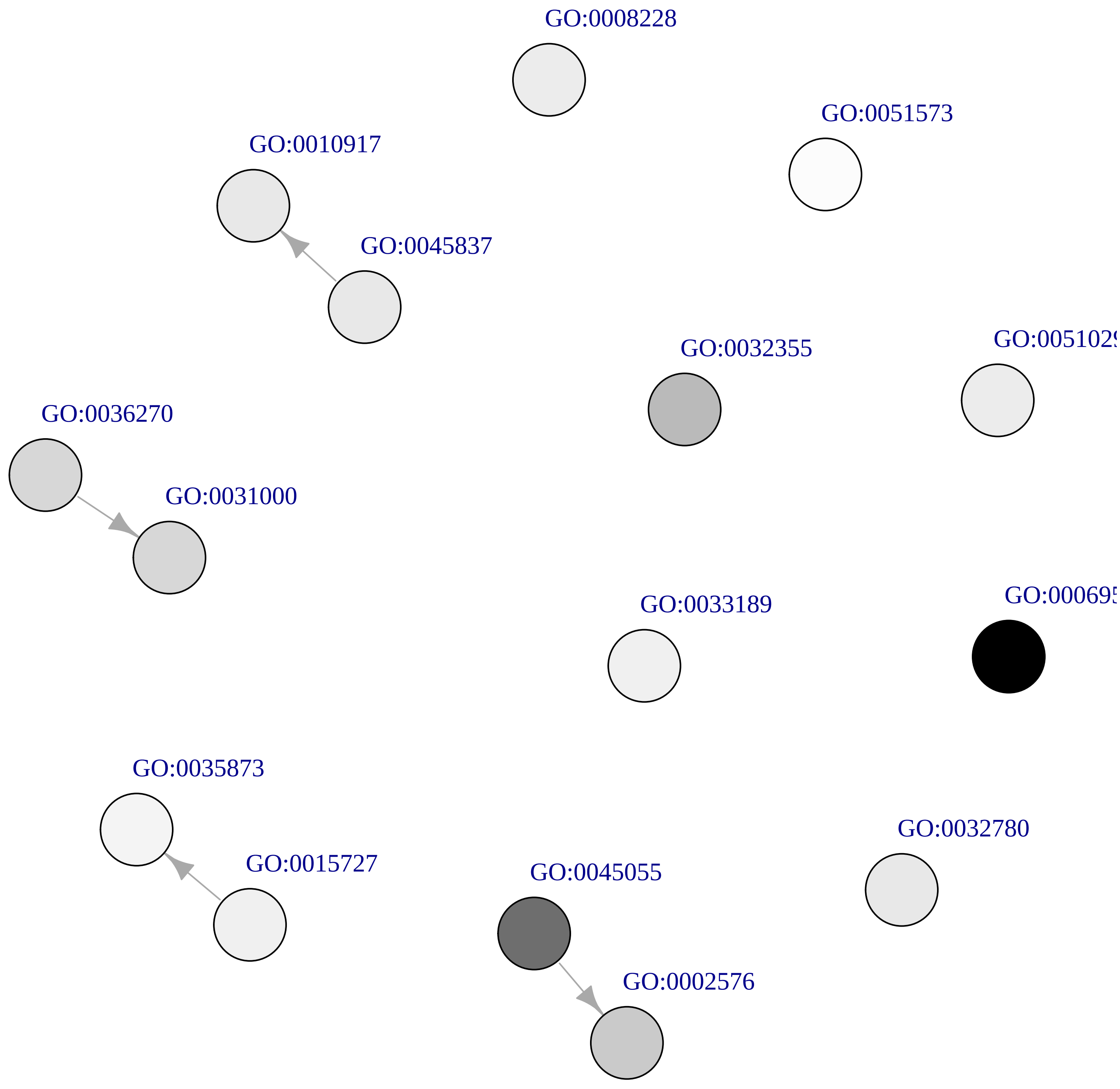

**Temp. UnderDEGs Cluster gray40**

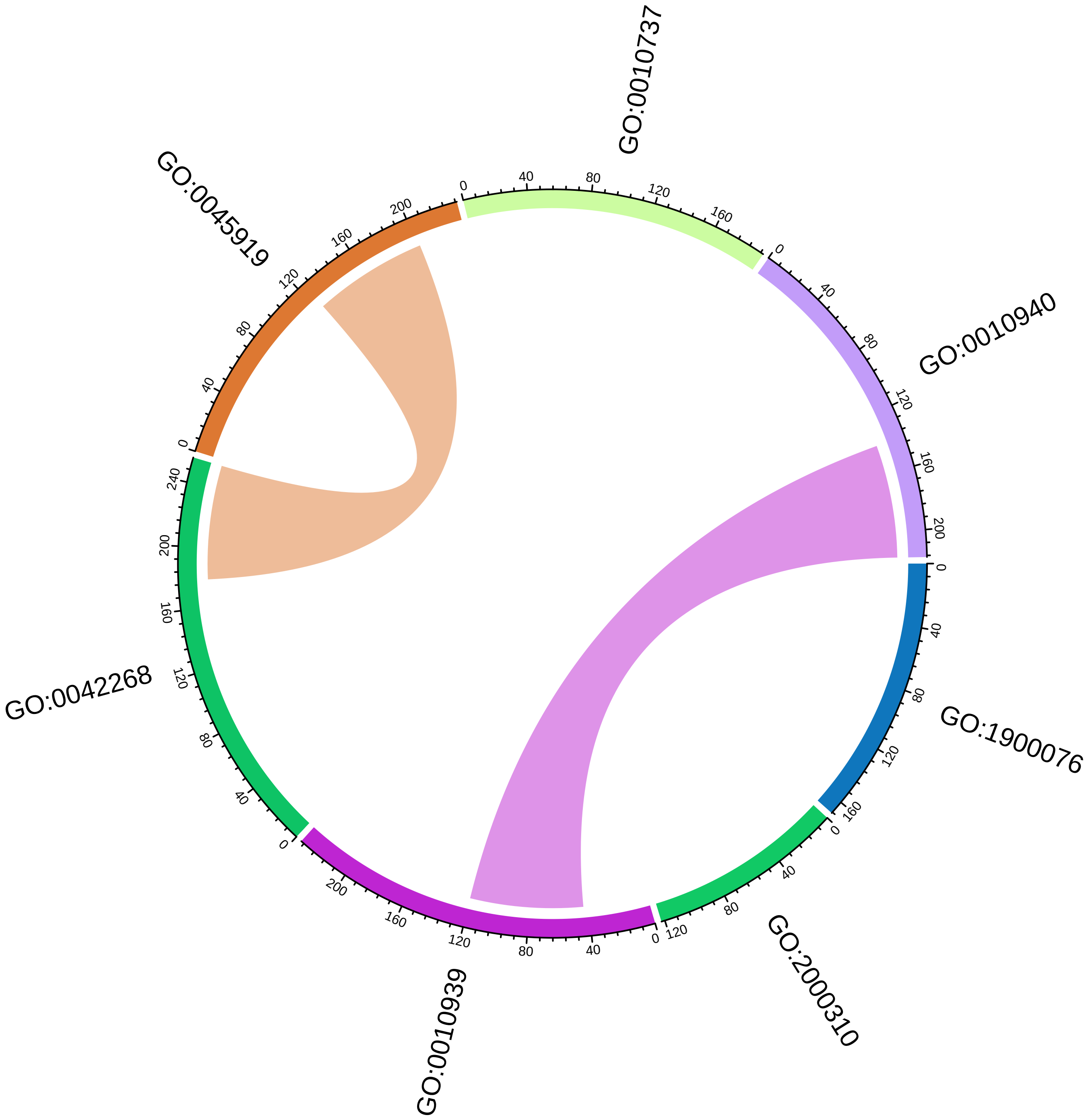

Temp. UnderDEGs Cluster gray40

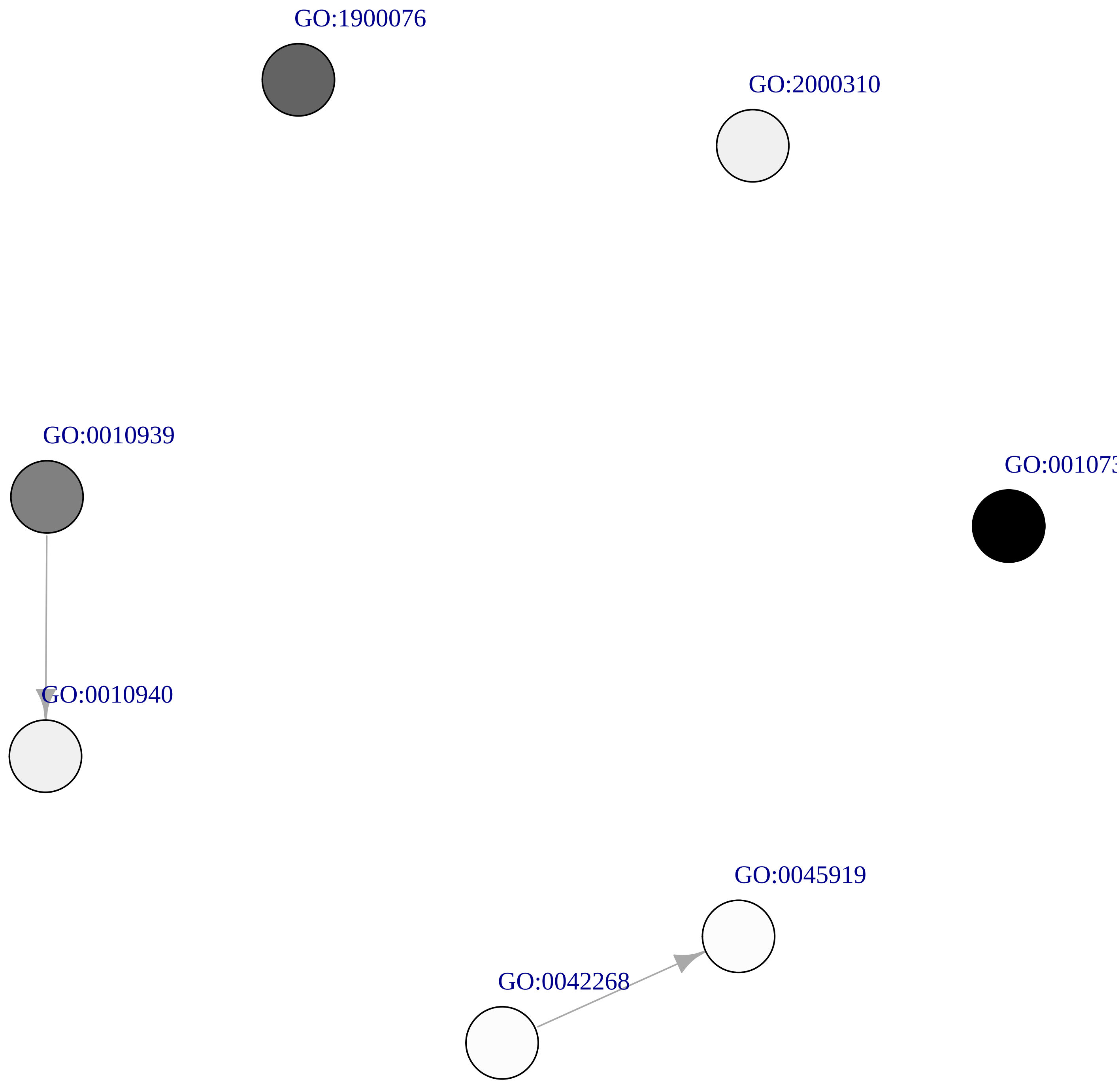

**Temp. UnderDEGs Cluster violetred2**

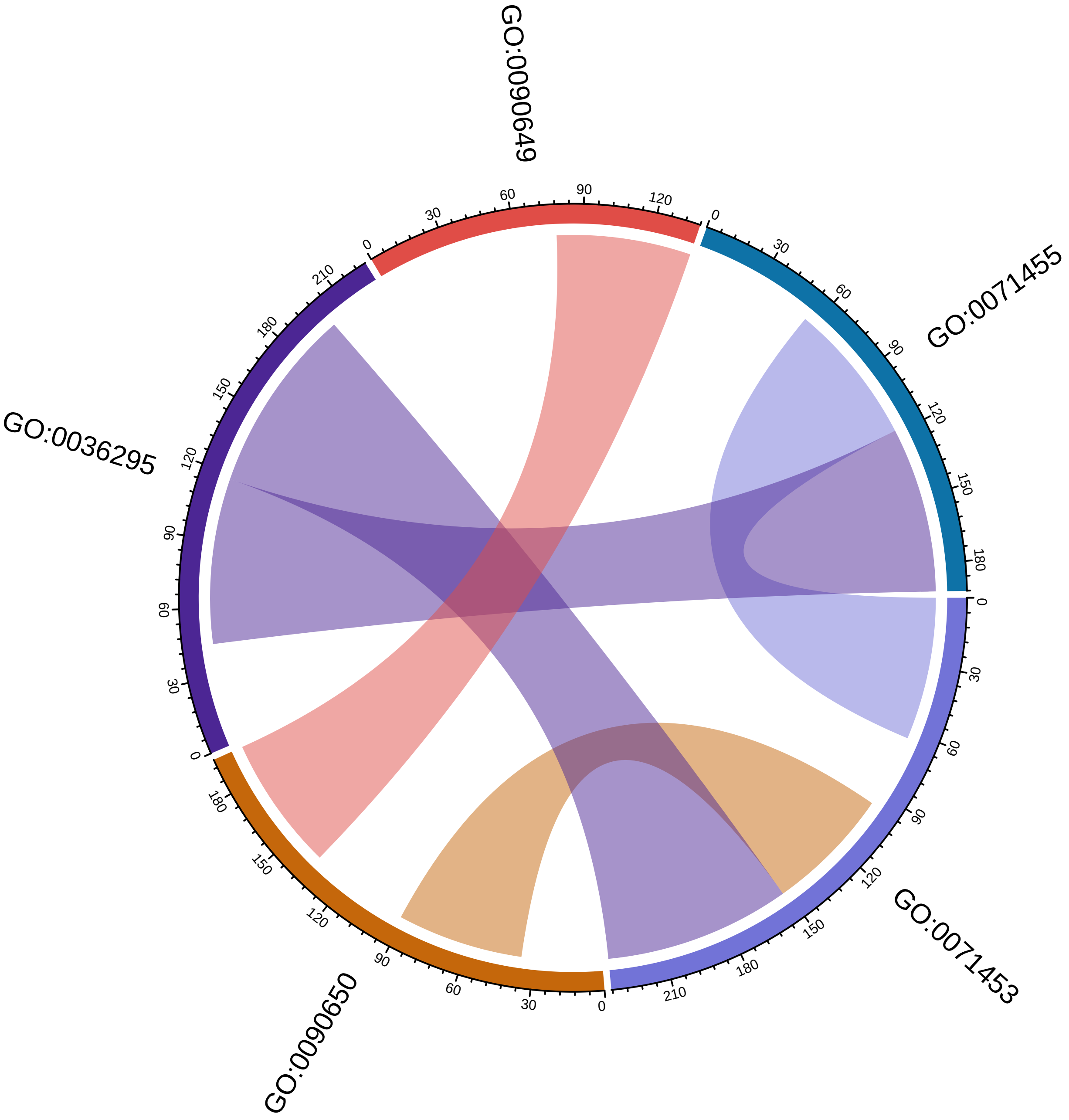

**Temp. UnderDEGs Cluster violetred2**

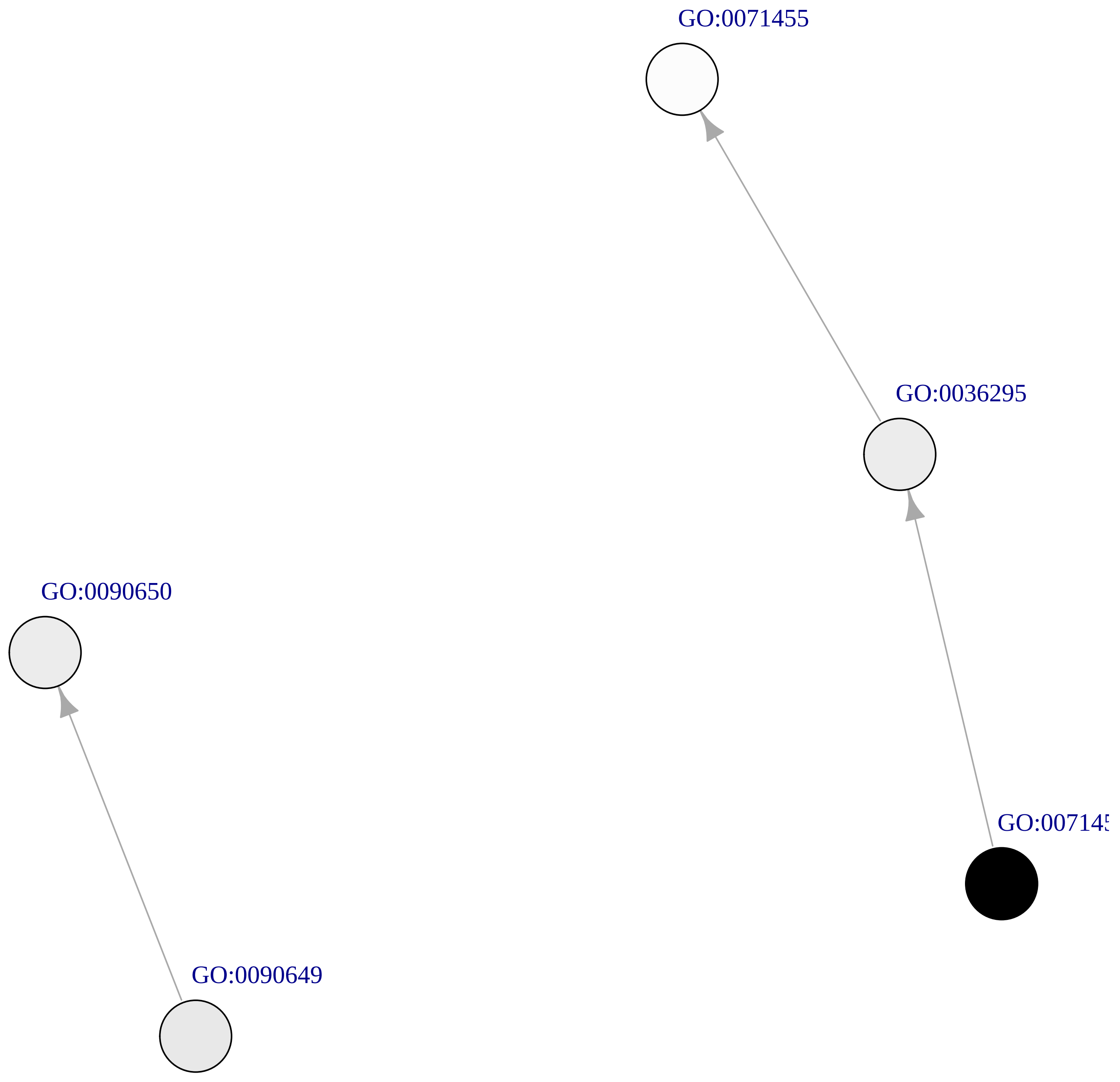
