## Supplementary Figures for "Transcriptional response of the calcification and stress response toolkits in an octocoral under heat and pH stress": Supplementary_Figures_31.pdf

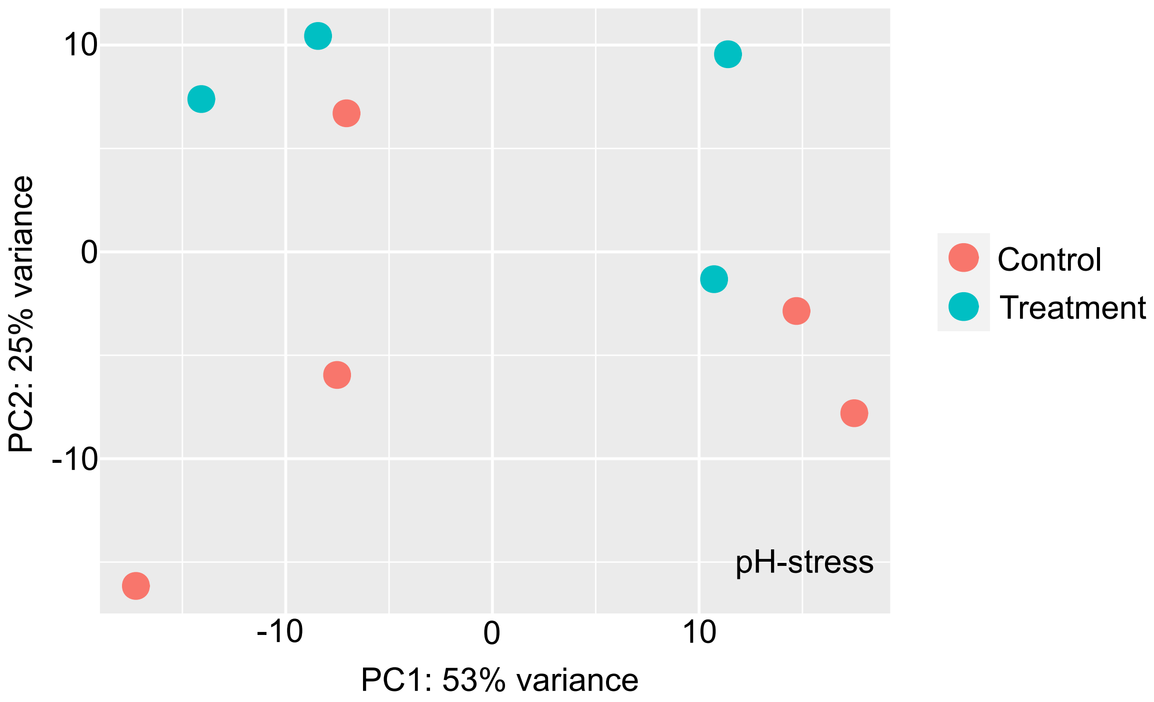


Supplementary Figure 17: Principal Component Analysis showing the global gene expression response of *Pinnigorgia flava* colonies exposed to low pH water stress.
