## Supplementary Figures for "Transcriptional response of the calcification and stress response toolkits in an octocoral under heat and pH stress": Supplementary_Figures_32-49.pdf

Chord plots (Supplementary Figures 32, 34, 36, 38, 40, 42, 44, 46, and 48) showing the semantic similarity within clusters of GO-terms enriched in the set of significantly underexpressed genes in *P. flava* colonies exposed to low pH stress. For each cluster of semantically similar GO-terms, a directed acyclic subgraph (DAsG; Supplementary Figures 33, 35, 37, 39, 41, 43, 45, 47, and 49) showing the relation between the GO-terms within a cluster in the general Biological Process gene ontology.

Supplementary Figure 32: Chord plot for cluster “darkblue”
Supplementary Figure 33: DAsG for cluster “darkblue”

Supplementary Figure 34: Chord plot for cluster “darkgreen”
Supplementary Figure 35: DAsG for cluster “darkgreen”

Supplementary Figure 36: Chord plot for cluster “mediumorchid3”
Supplementary Figure 37: DAsG for cluster “mediumorchid3”

Supplementary Figure 38: Chord plot for cluster “orange3”
Supplementary Figure 39: DAsG for cluster “orange3”

Supplementary Figure 40: Chord plot for cluster “firebrick4”
Supplementary Figure 41: DAsG for cluster “firebrick4”

Supplementary Figure 42: Chord plot for cluster “gray40”
Supplementary Figure 43: DAsG for cluster “gray40”

Supplementary Figure 44: Chord plot for cluster “violetred2”
Supplementary Figure 45: DAsG for cluster “violetred2”

Supplementary Figure 46: Chord plot for cluster “red”
Supplementary Figure 47: DAsG for cluster “red”

Supplementary Figure 48: Chord plot for cluster “blue”
Supplementary Figure 49: DAsG for cluster “blue”

NOTE: cluster names refer to the Rcolor used to plot them in the dendrogram.

**pH UnderDEGs Cluster darkblue**

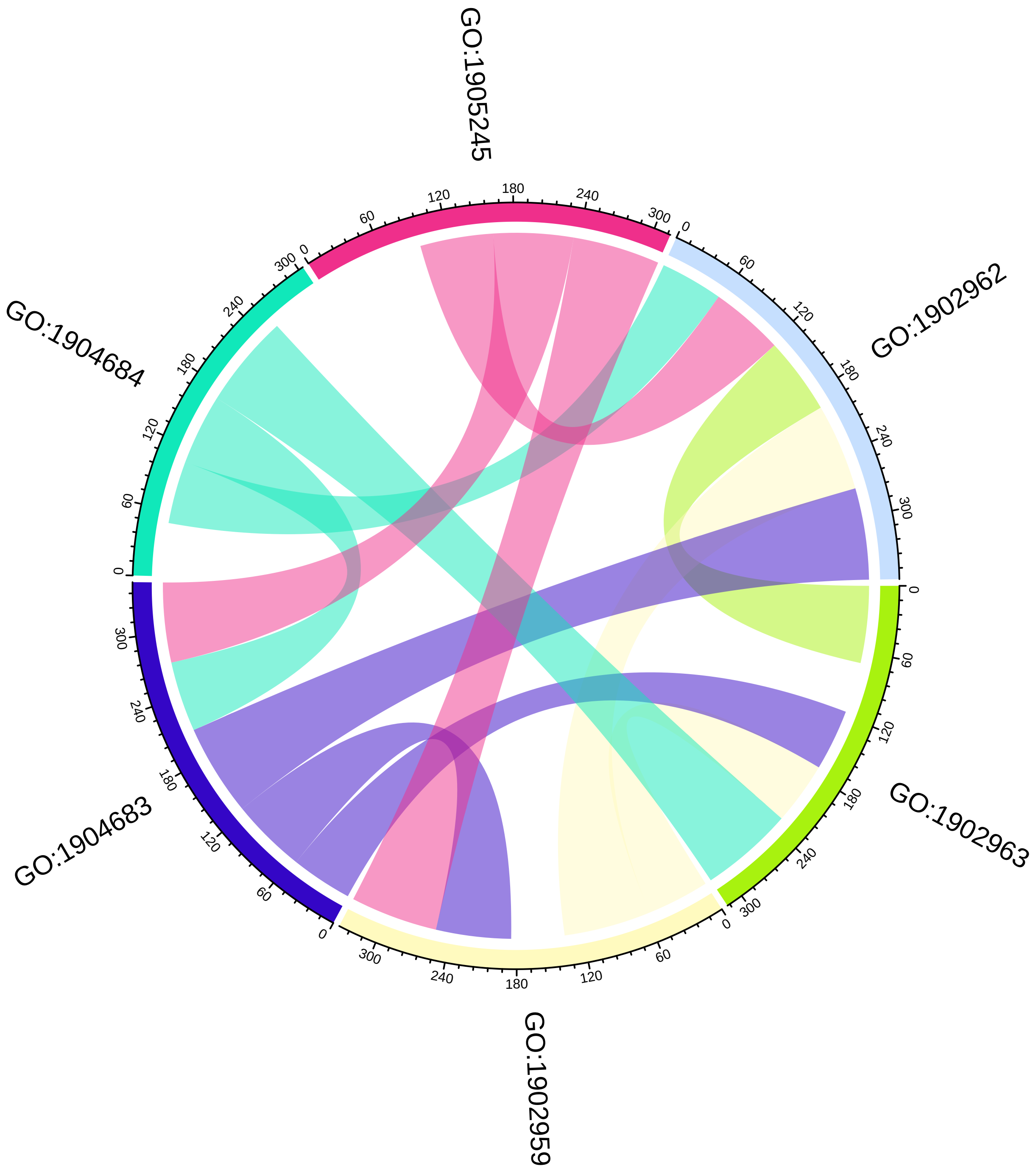

#### pH UnderDEGs Cluster darkblue

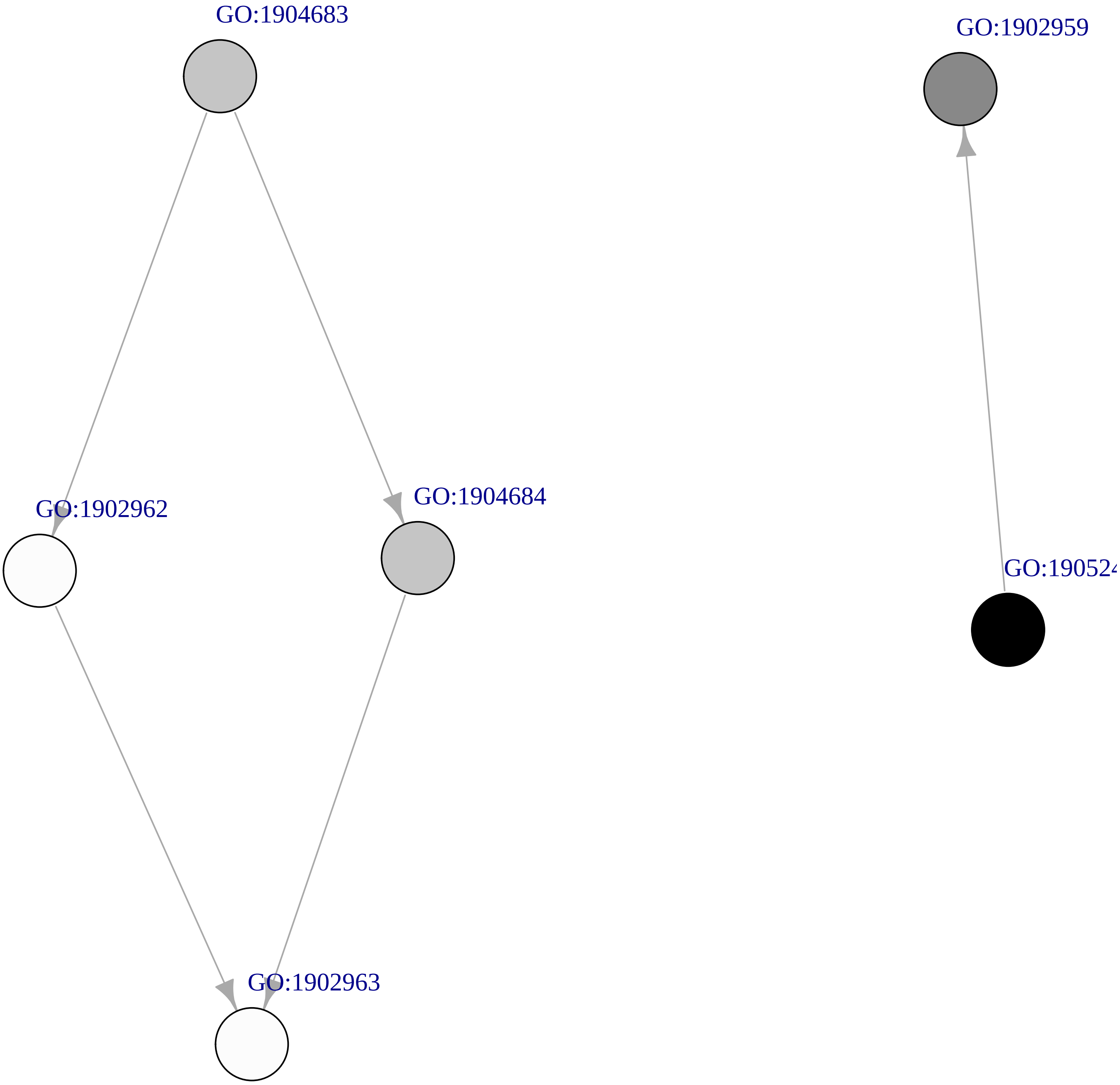

### pH UnderDEGs Cluster darkgreen

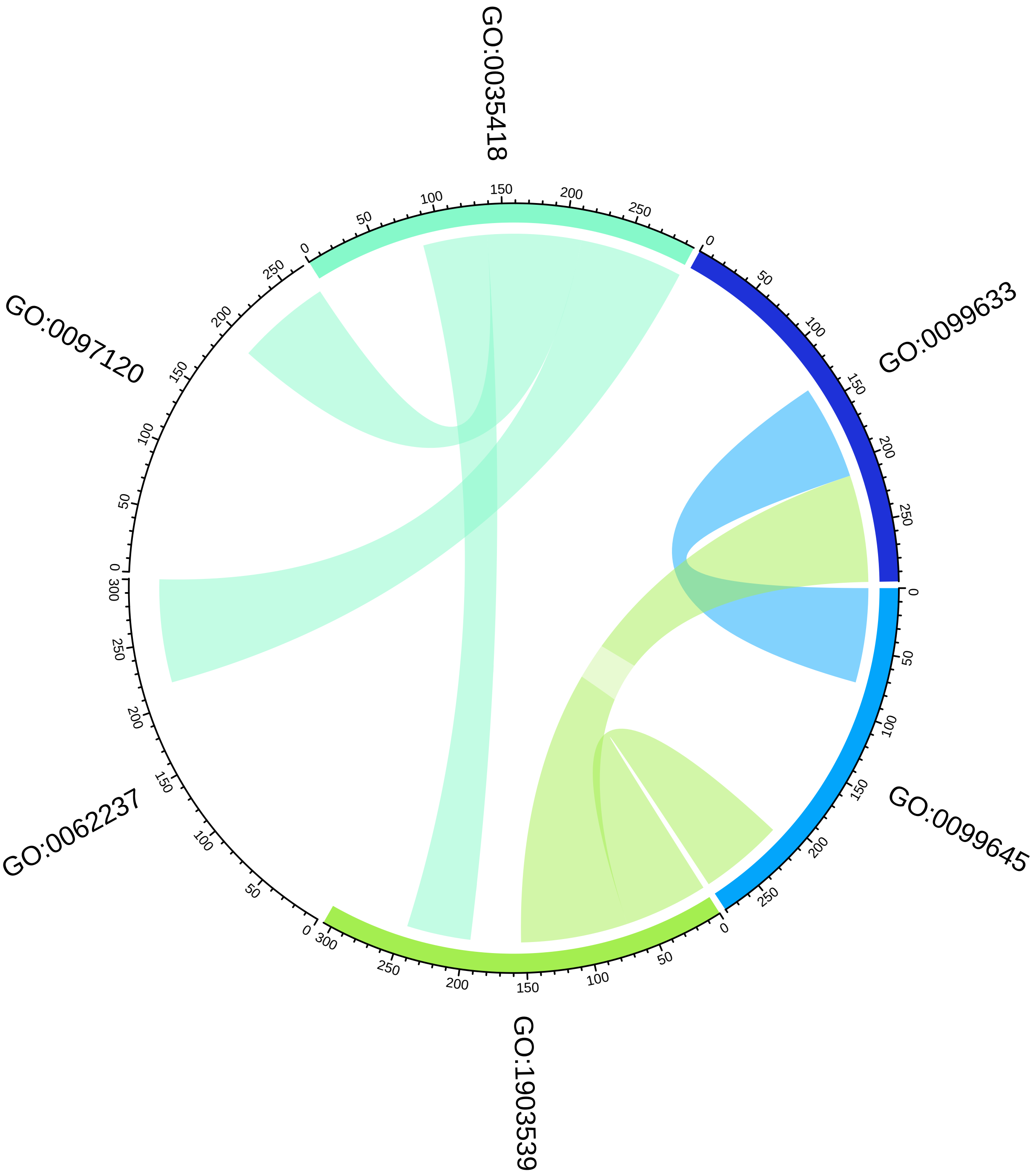

#### pH UnderDEGs Cluster darkgreen

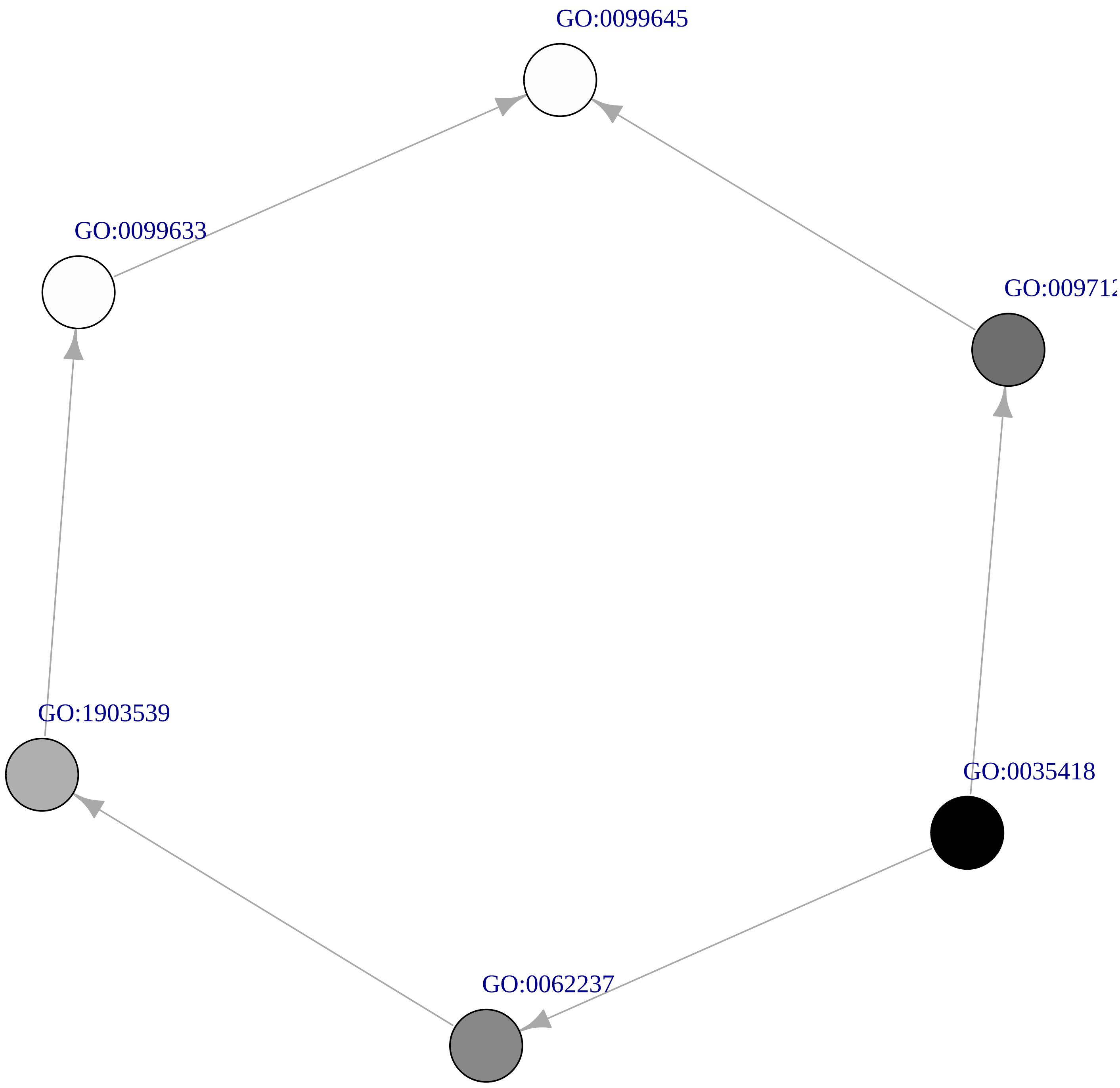

### pH UnderDEGs Cluster mediumorchid3

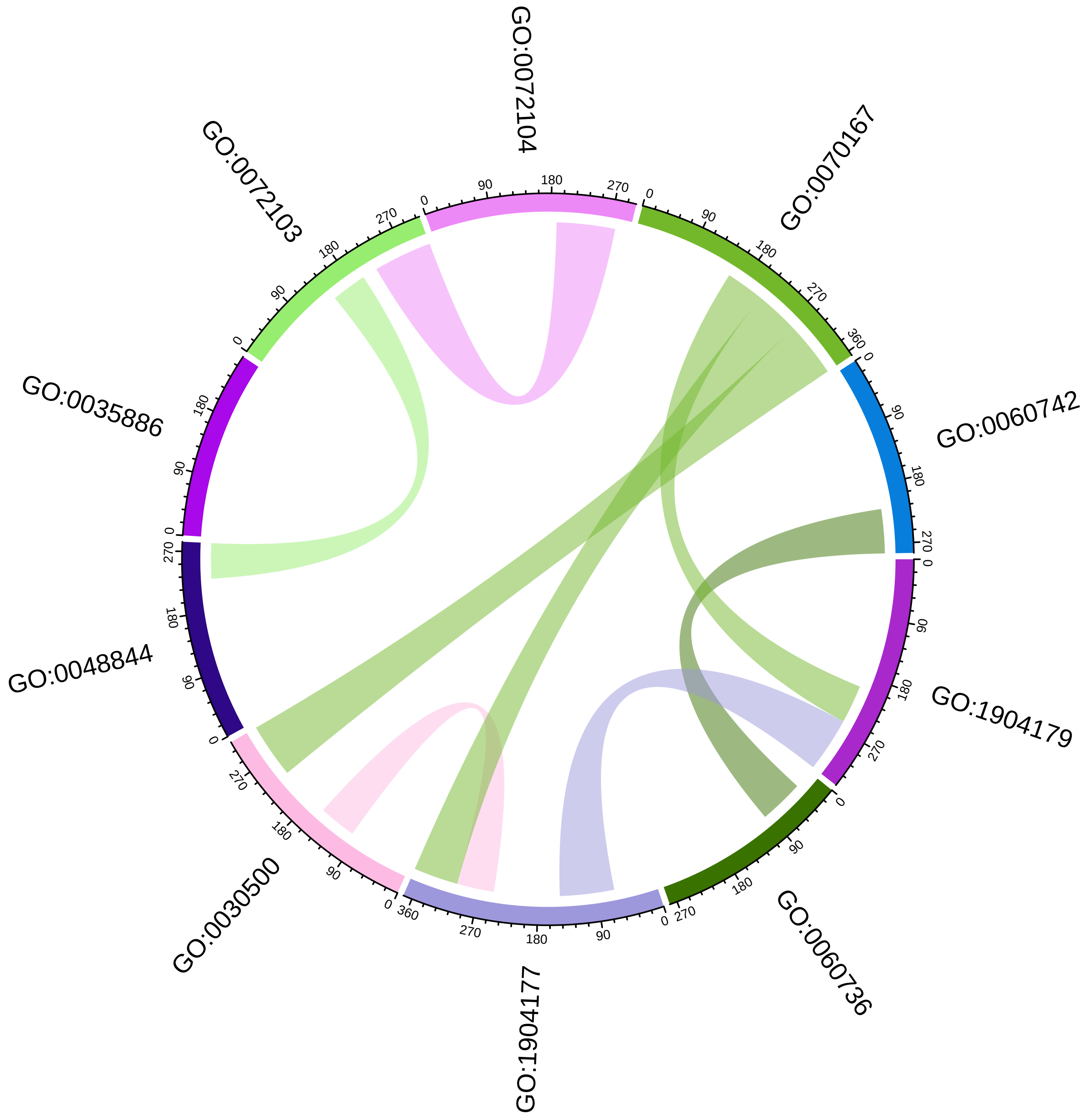

### pH UnderDEGs Cluster mediumorchid3

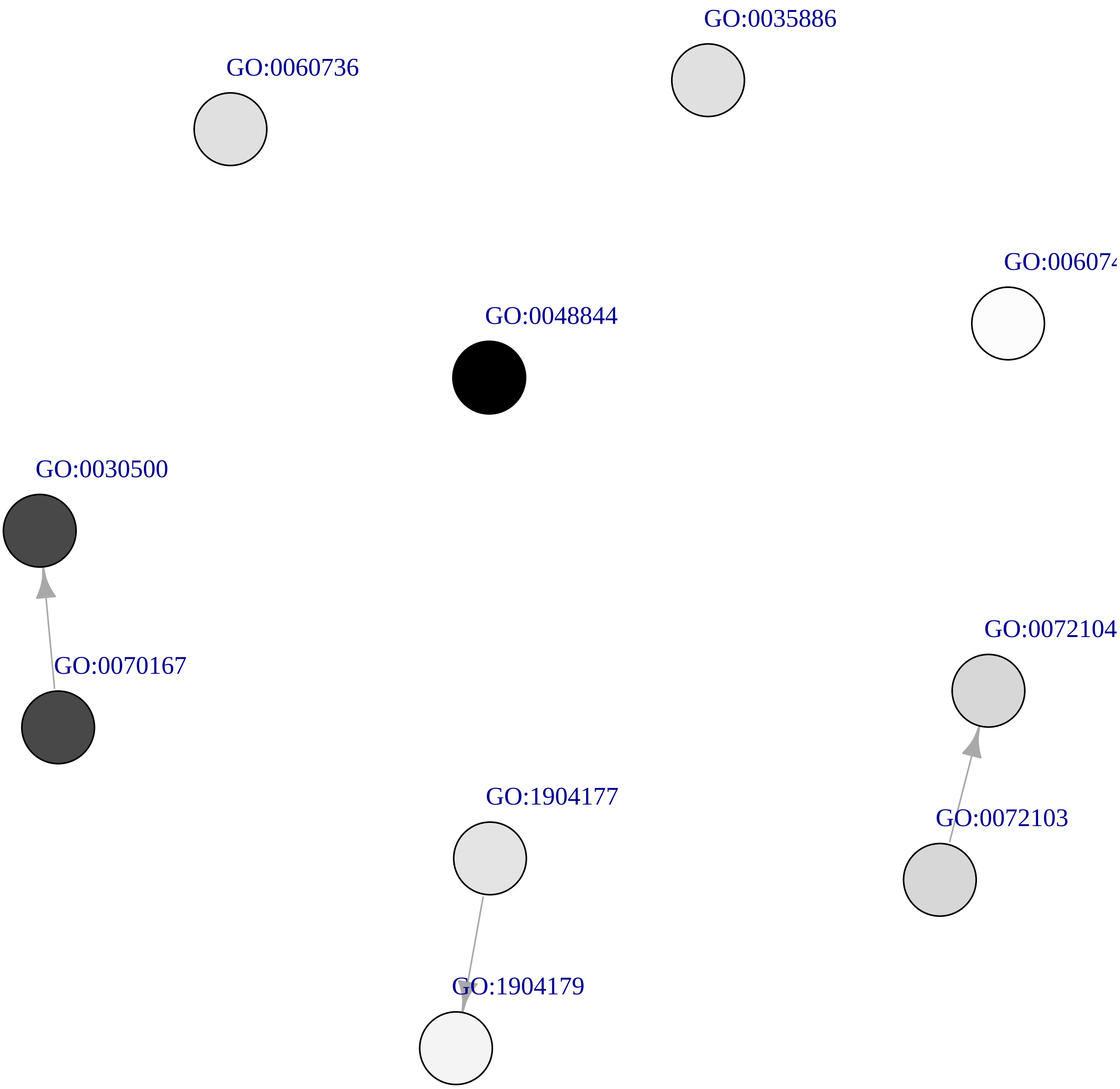

### pH UnderDEGs Cluster orange3

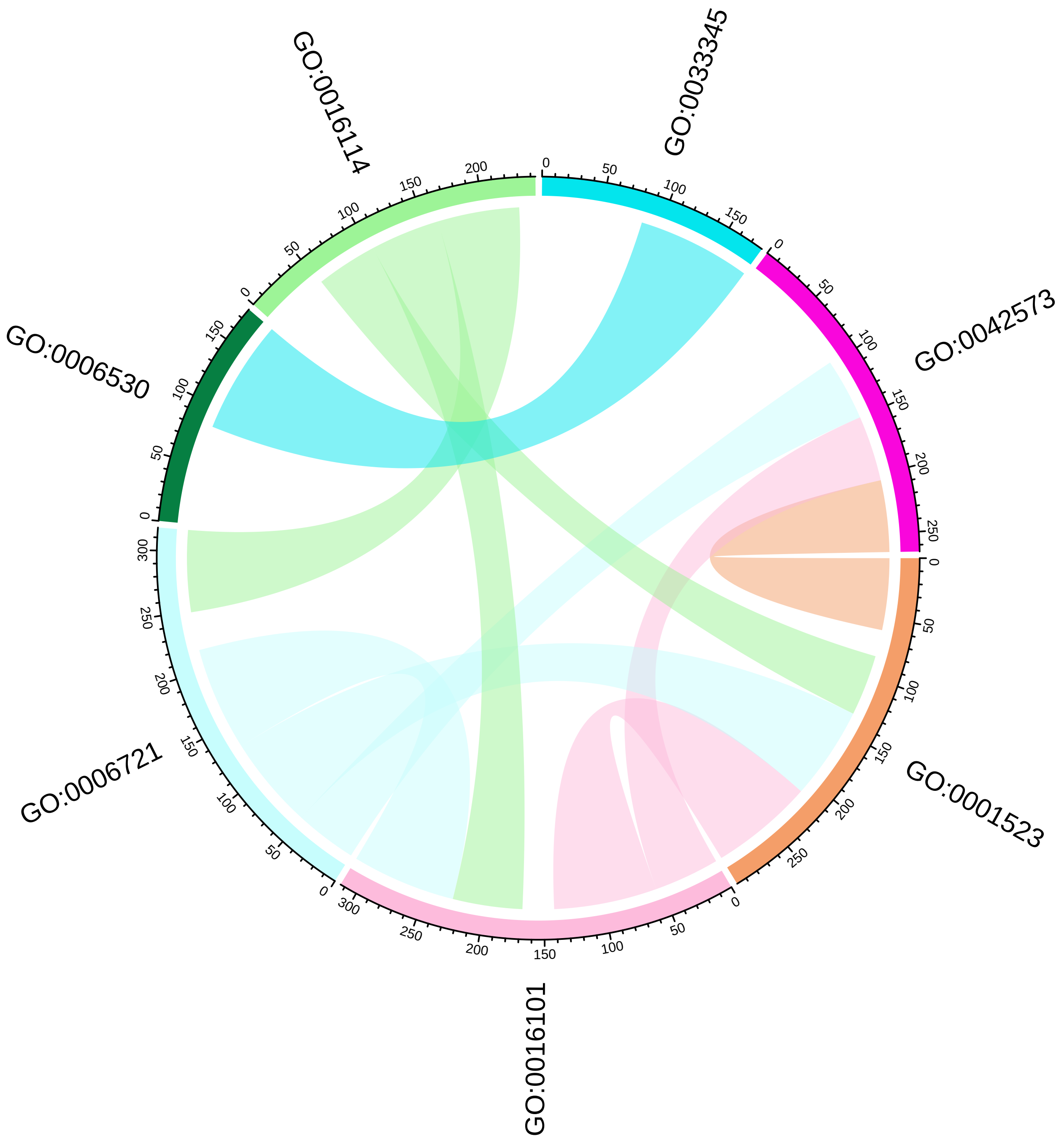

### pH UnderDEGs Cluster orange3

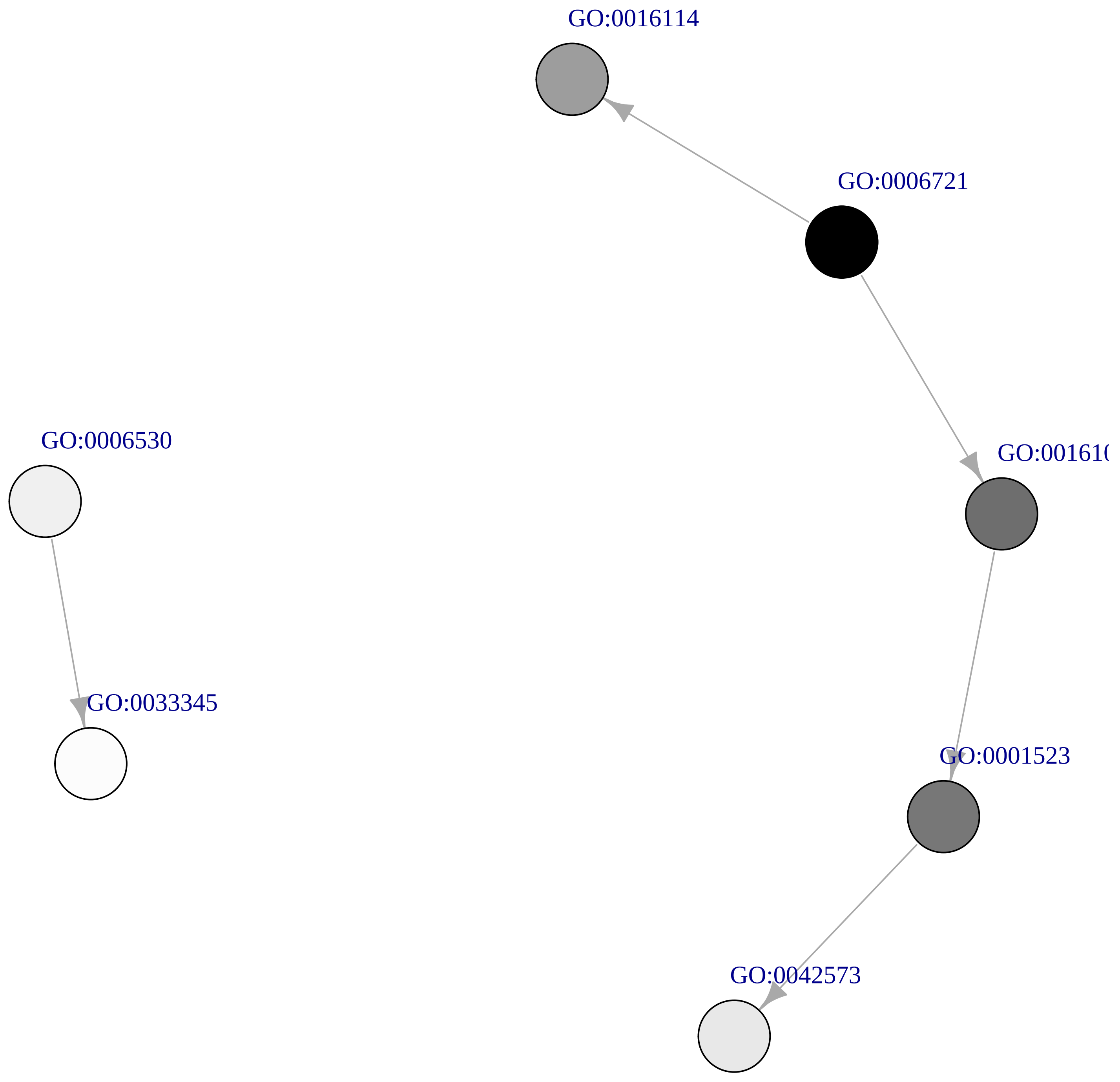

### pH UnderDEGs Cluster firebrick4

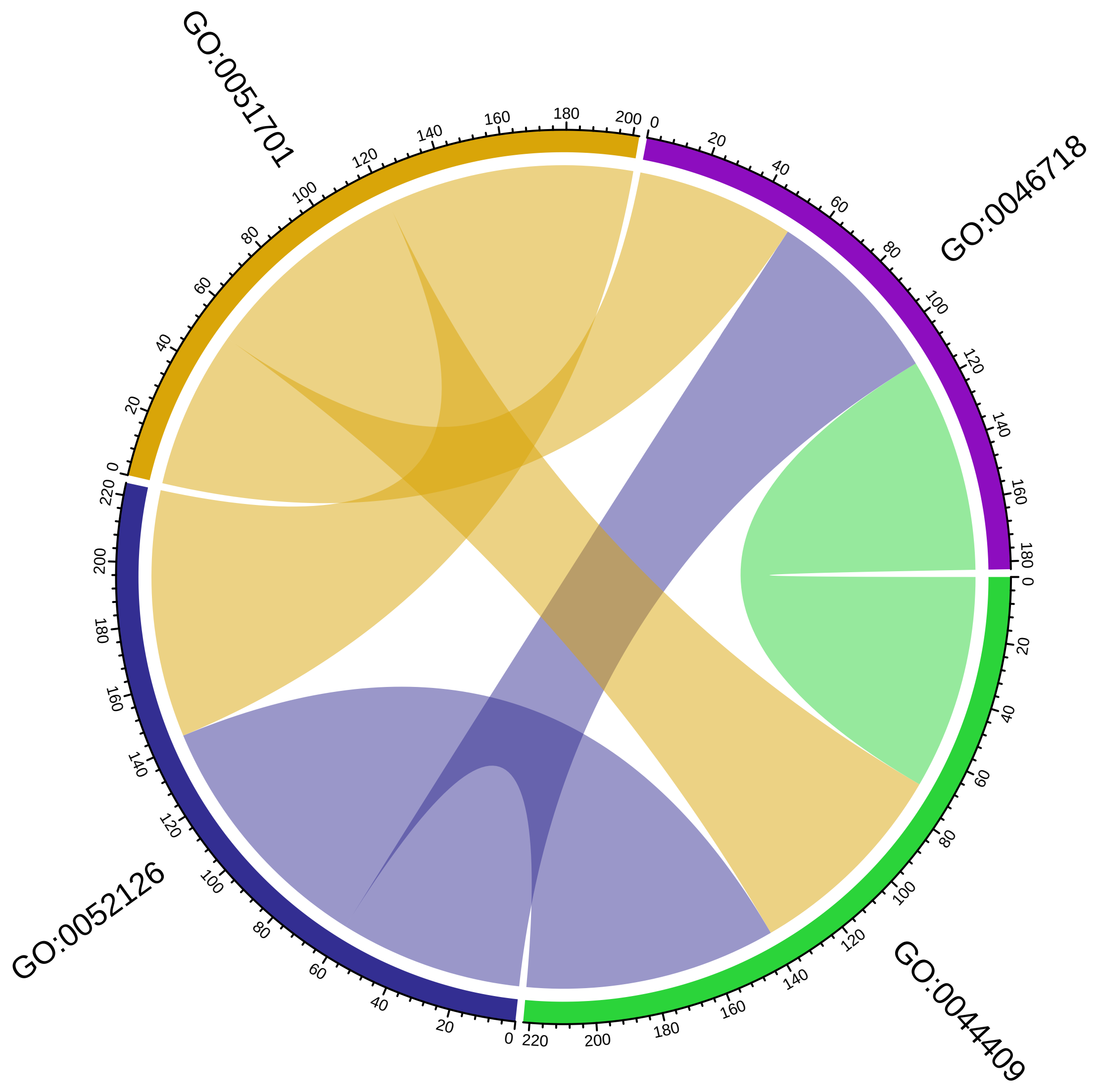

pH UnderDEGs Cluster firebrick4

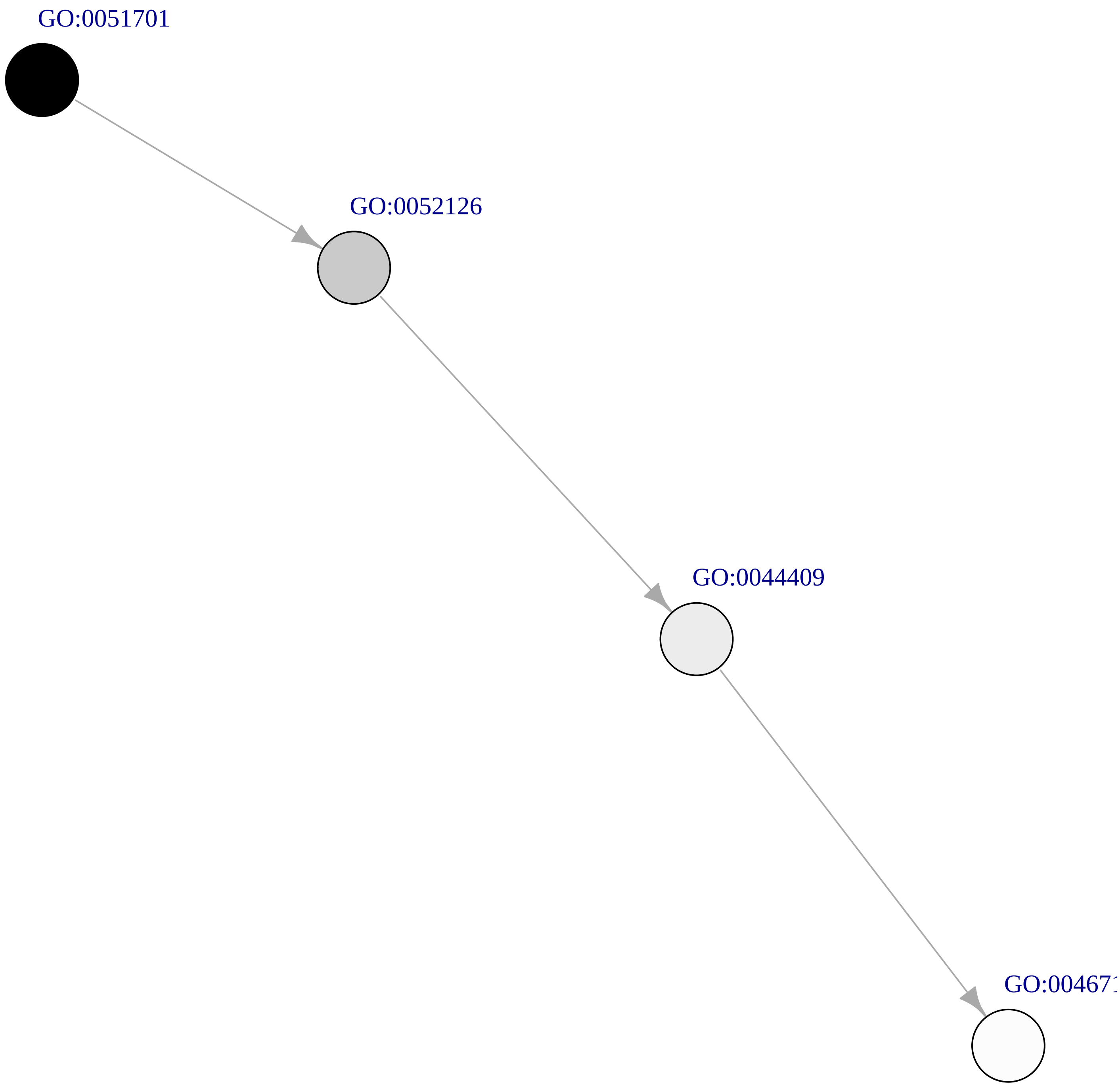

### pH UnderDEGs Cluster gray40

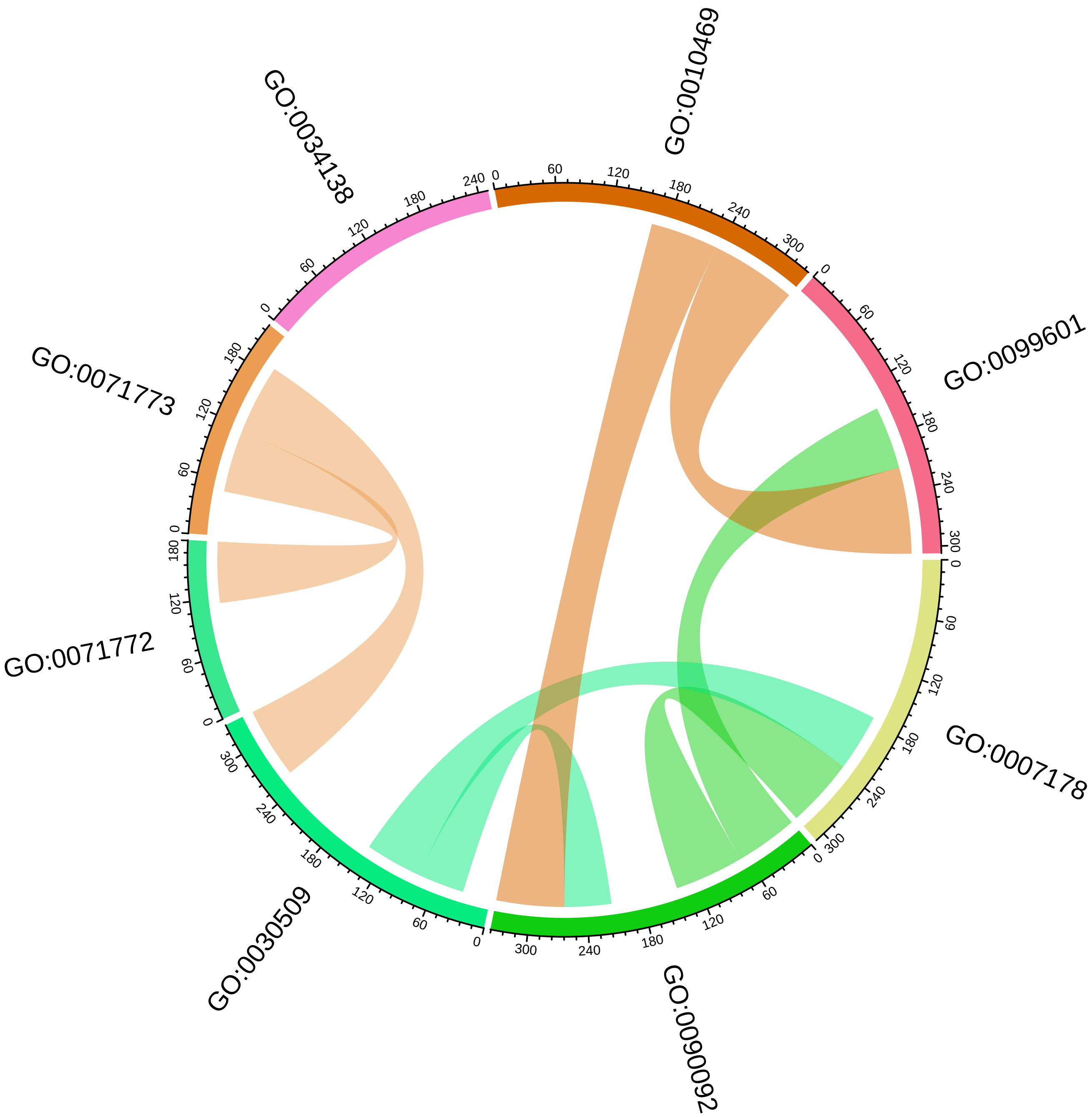

#### pH UnderDEGs Cluster gray40

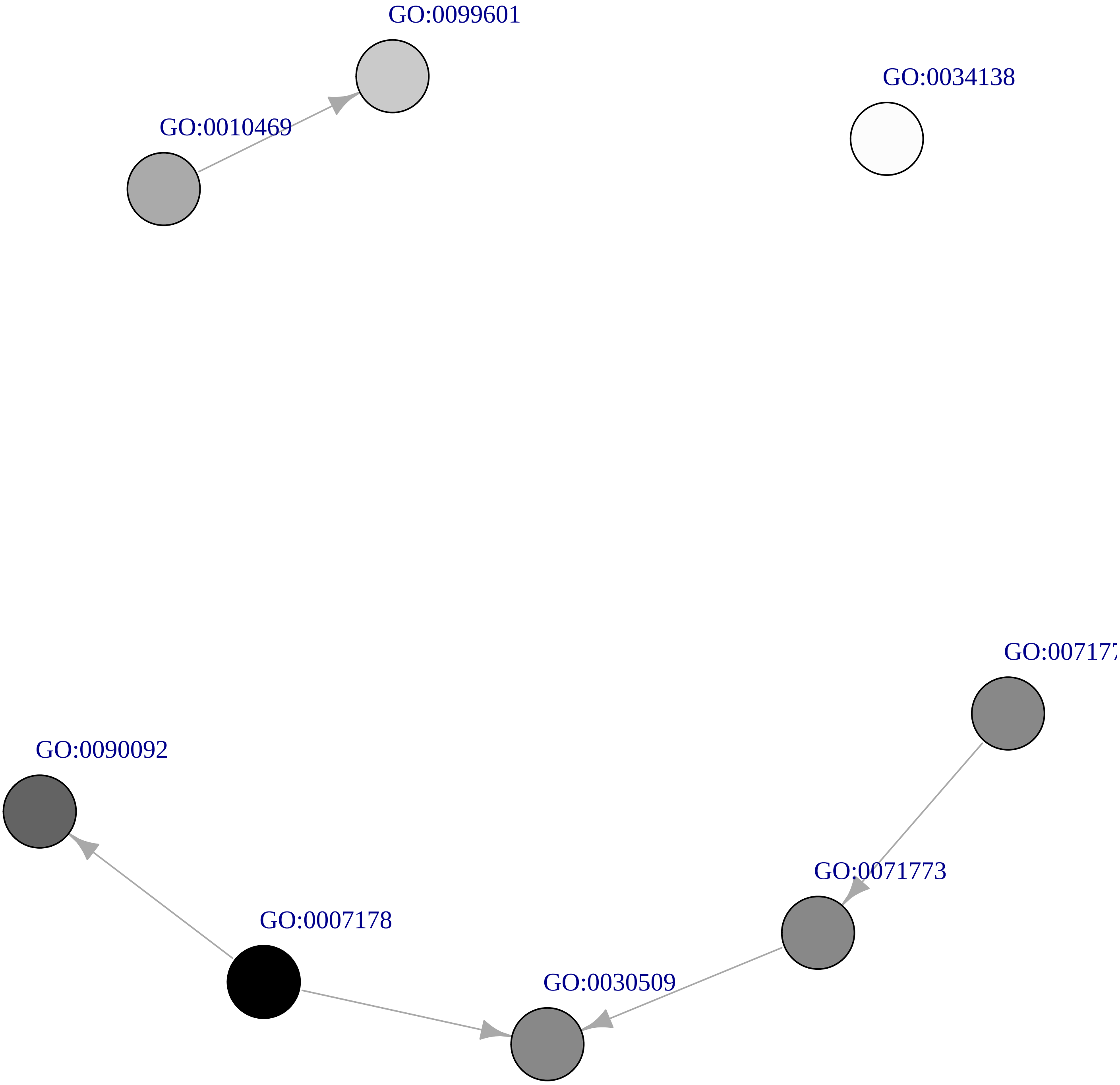

### pH UnderDEGs Cluster violetred2

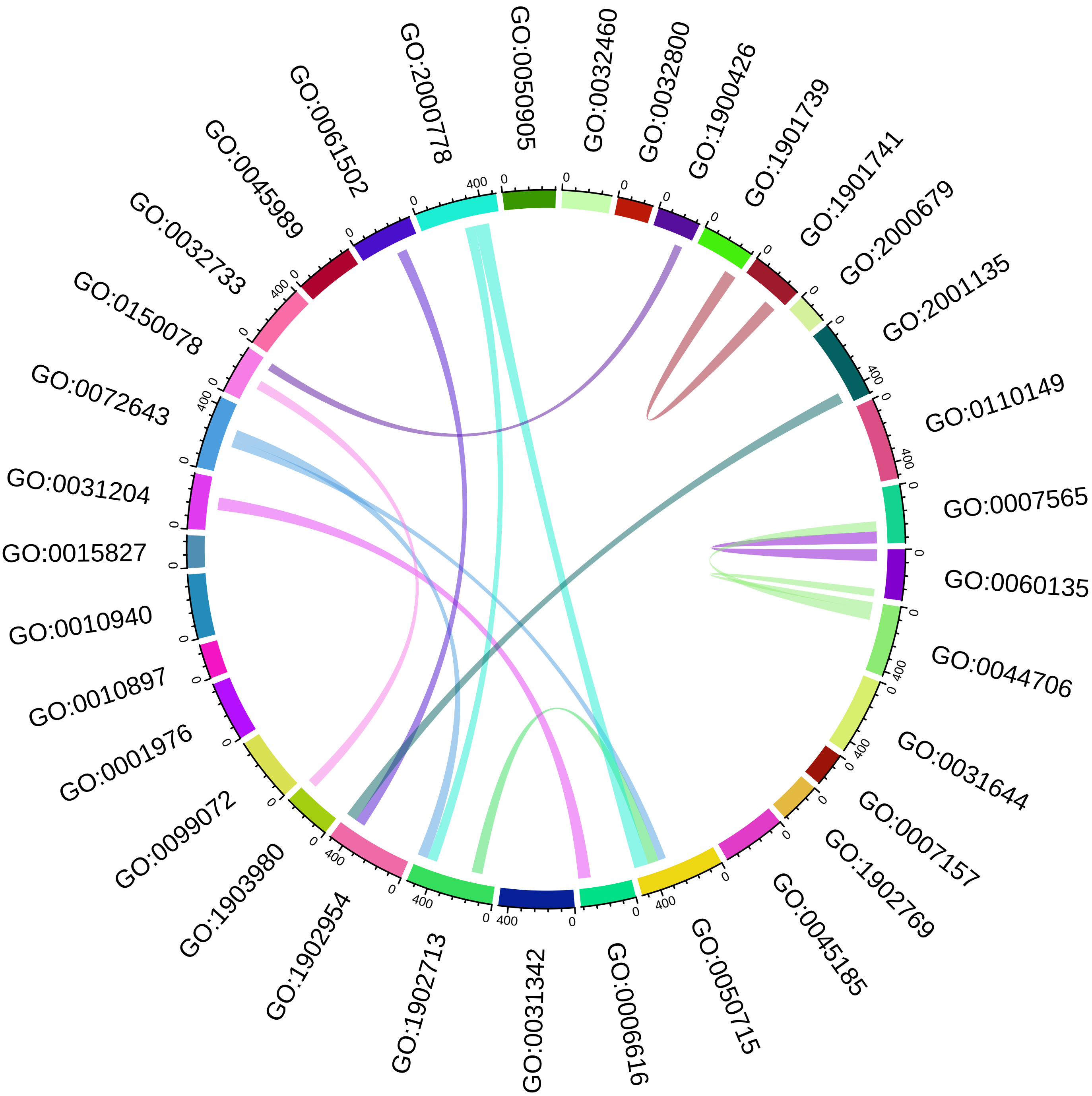

### pH UnderDEGs Cluster violetred2

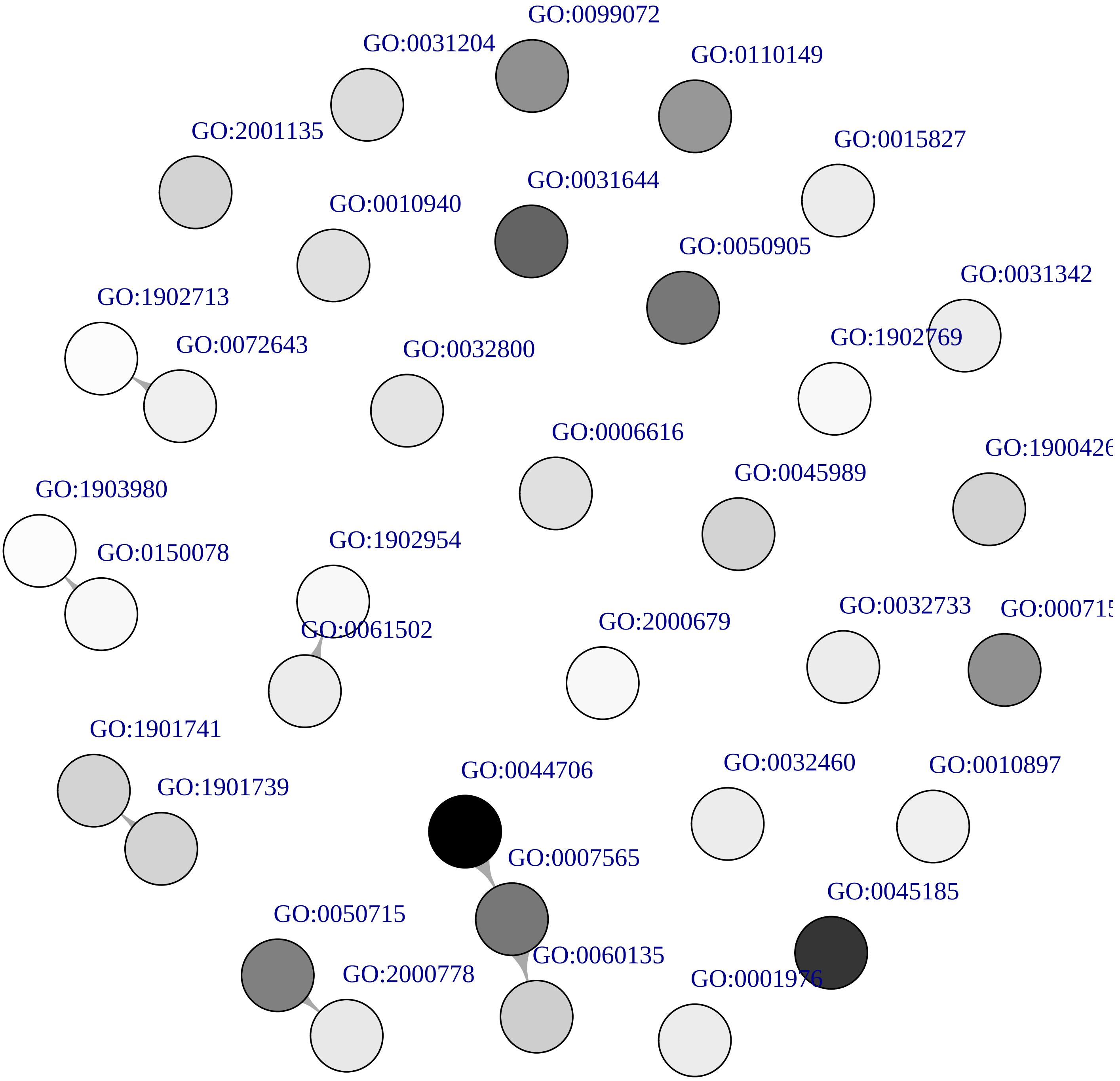

### pH UnderDEGs Cluster red

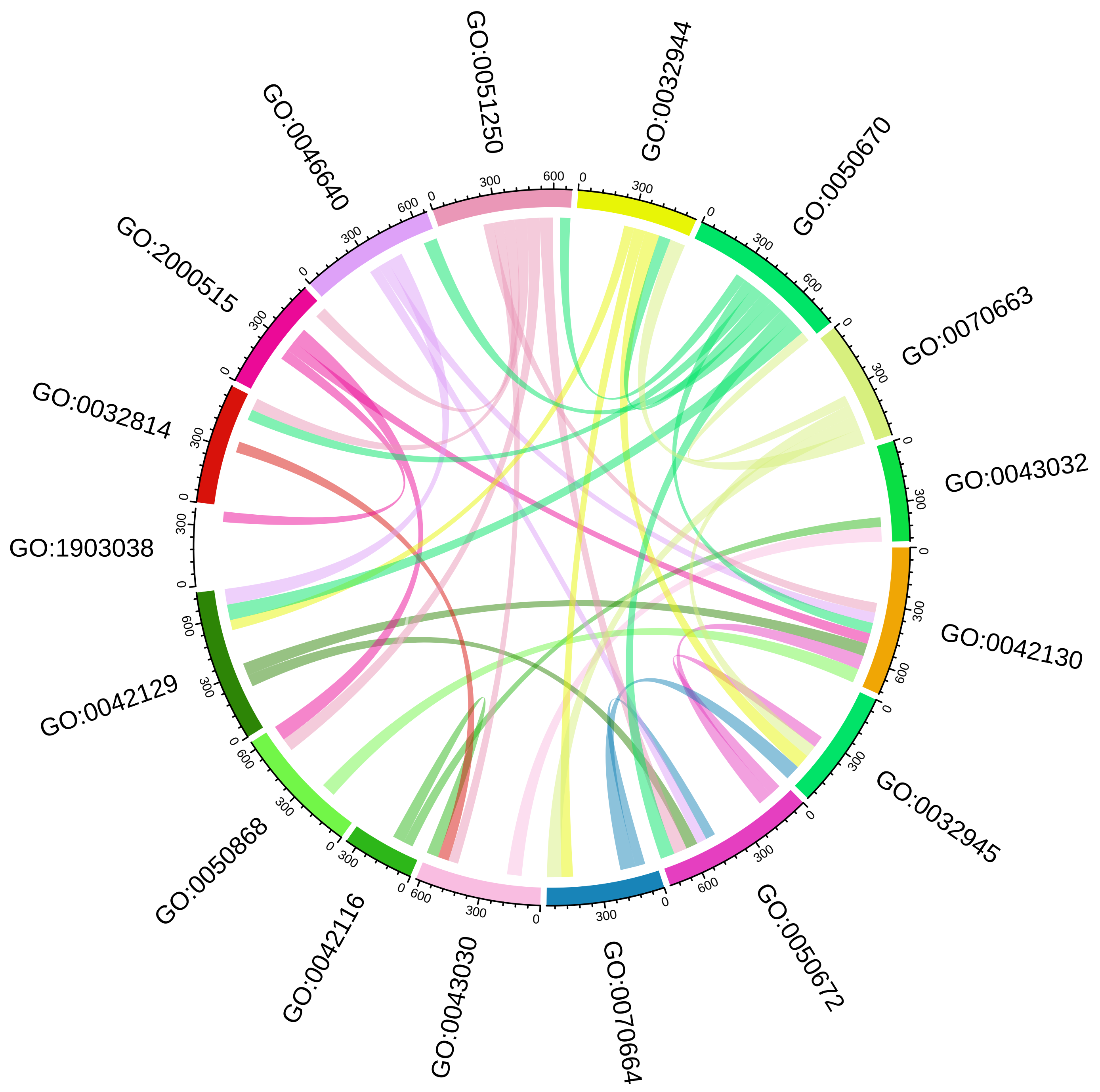

### pH UnderDEGs Cluster red

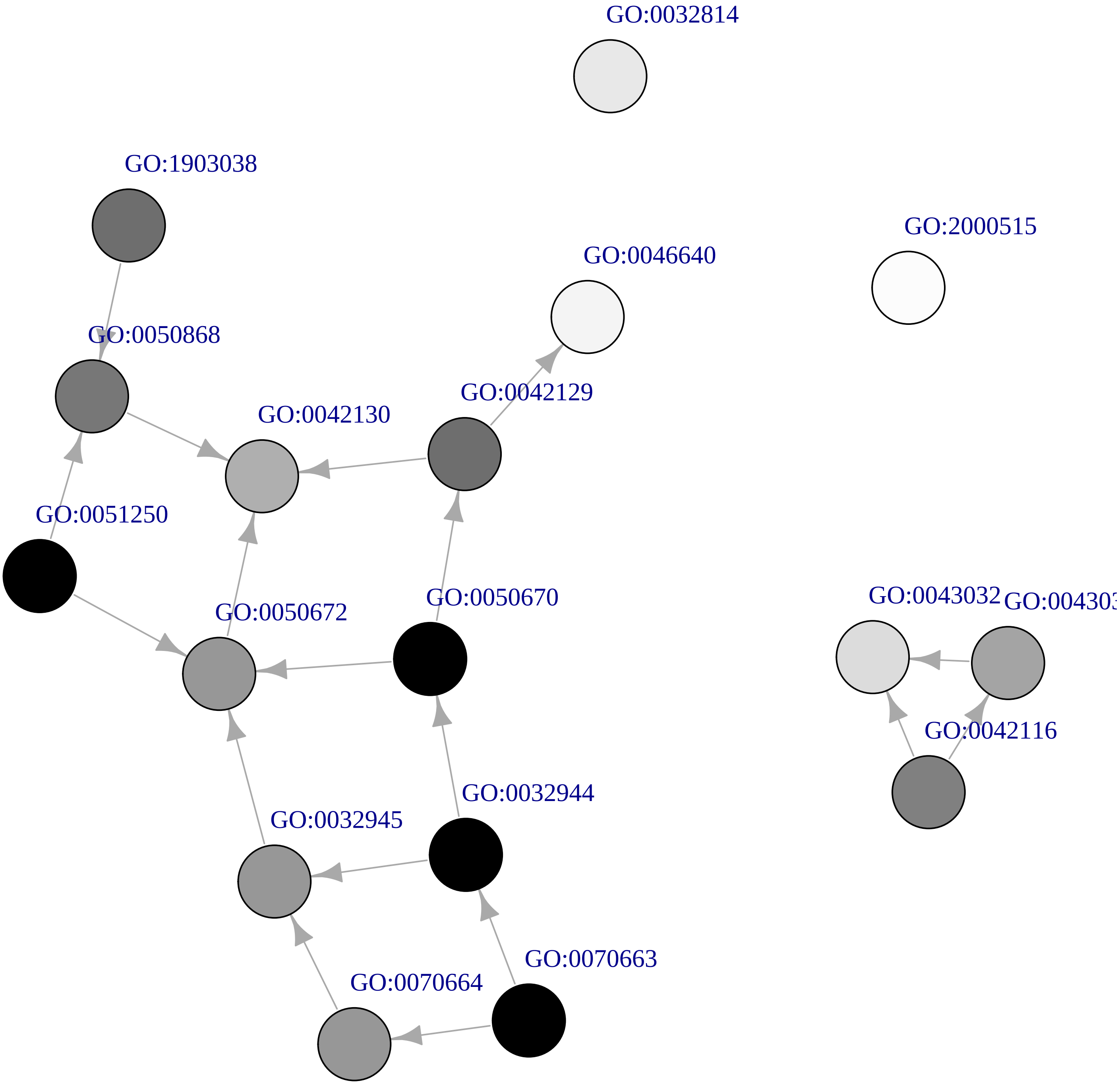

#### pH UnderDEGs Cluster blue

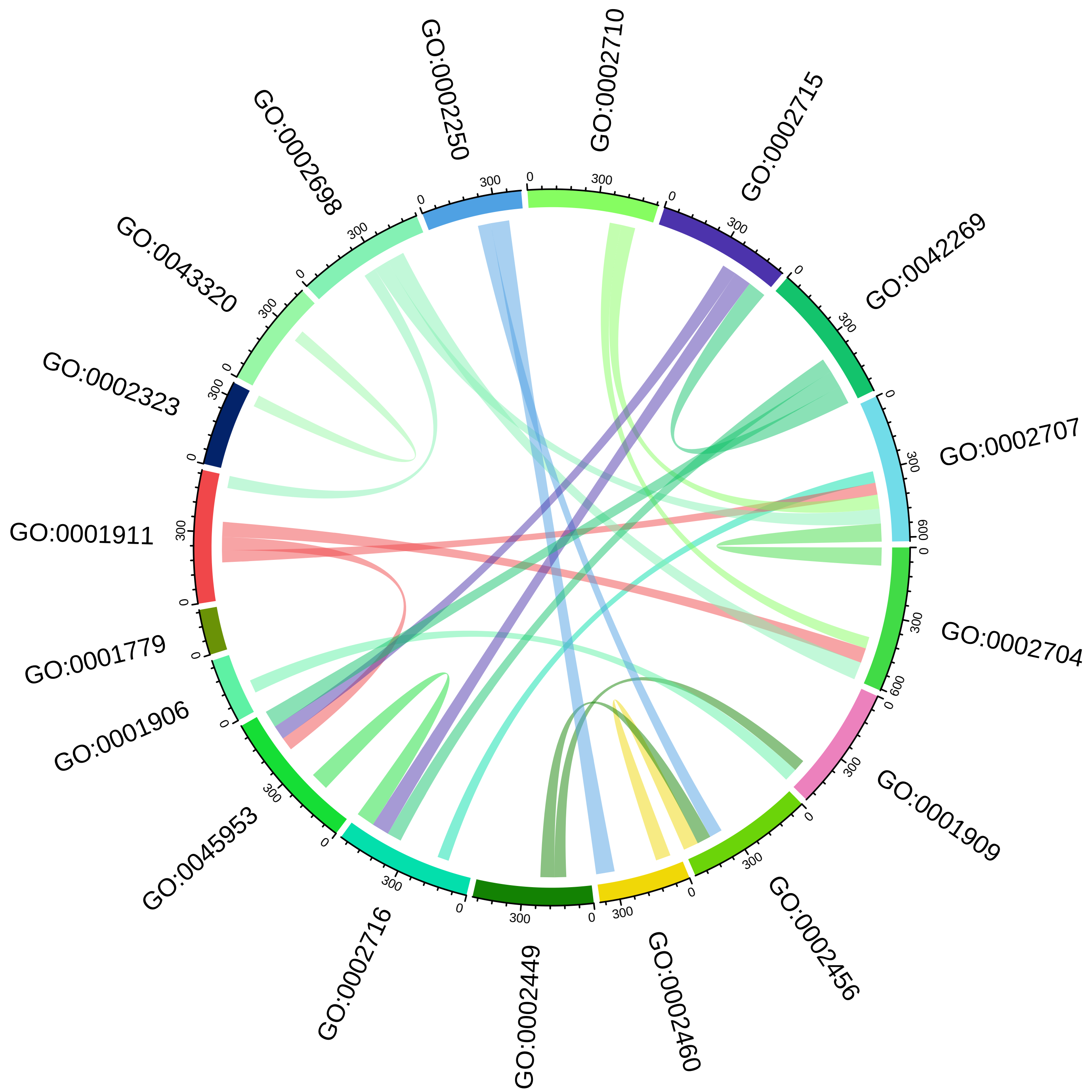

### pH UnderDEGs Cluster blue

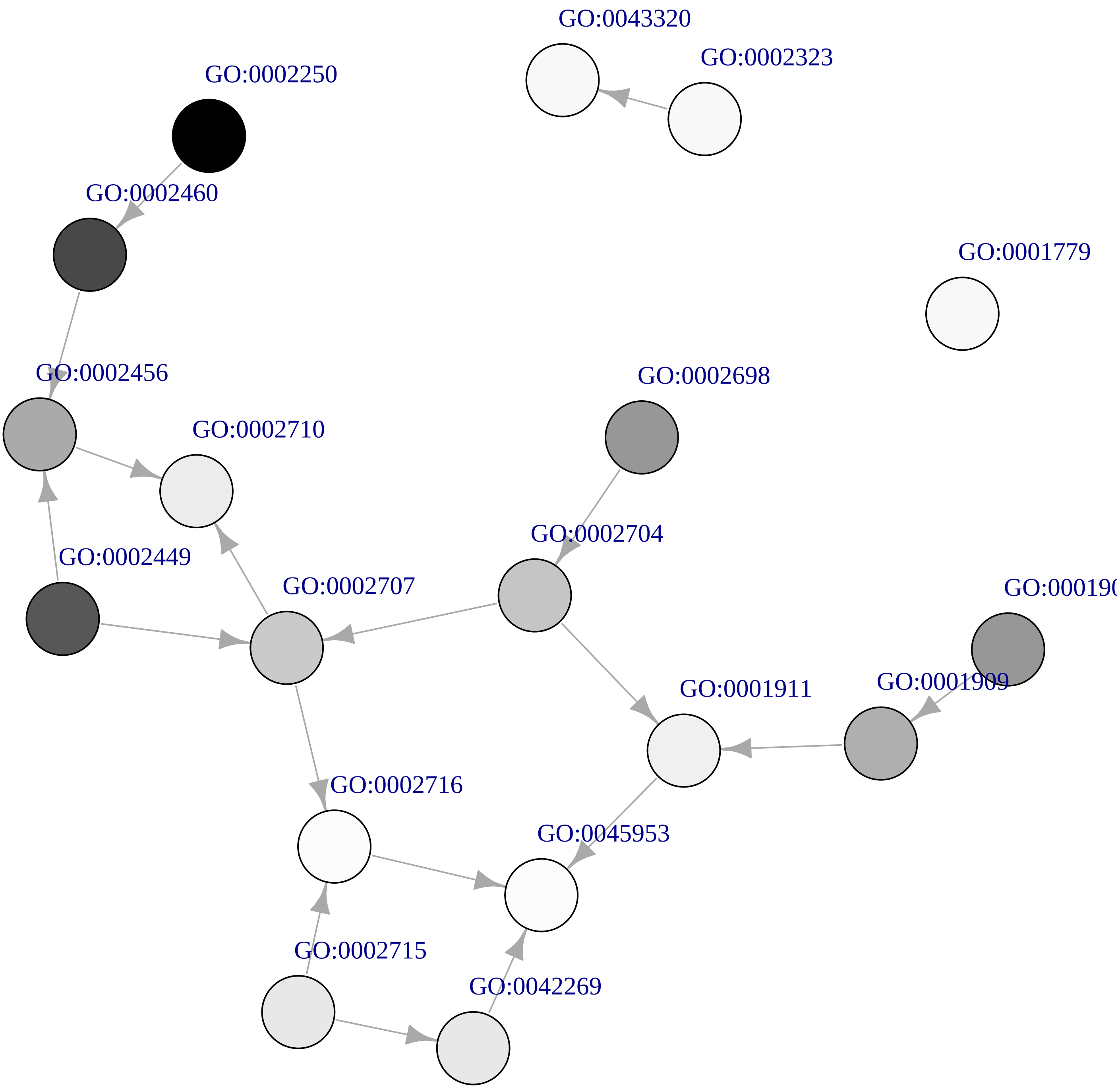
